# Lightning Fast and Highly Sensitive Full-Length Single-cell sequencing using FLASH-Seq

**DOI:** 10.1101/2021.07.14.452217

**Authors:** Vincent Hahaut, Dinko Pavlinic, Cameron Cowan, Simone Picelli

## Abstract

In the last 10 years, single-cell RNA-sequencing (scRNA-seq) has undergone exponential growth. Emulsion droplets methods^1–3^, such as those commercialized by 10x Genomics, have allowed researchers to analyze tens of thousands of cells in parallel in a robust and reproducible way. However, in contrast to SMART-based full-length sequencing protocols^4,5^, these methods interrogate only the outer portion of the transcripts and still lack the required sensitivity for analyzing comprehensively the transcriptome of individual cells. Building upon the existing SMART-seq forerunners protocols^4,5^, we developed FLASH-Seq (FS), a new scRNA-seq method which displays greater sensitivity while decreasing incubation times and reducing the number of processing steps compared to its predecessors. The entire FS protocol - from lysed cells to pooled cDNA libraries - can be performed in ~4.5 hours, is automation-friendly and can be easily miniaturized to decrease costs.

---

Switching Mechanism at the 5′ end of the RNA Template (SMART) protocols, such as Smart-seq2^4^ (SS2), consist of an initial reverse transcription reaction using a template-switching reverse transcriptase (RT) such as the murine Moloney Leukemia Virus-based (MMLV) enzyme, which adds a known sequence at the 5’-end of the cDNA. The full-length cDNA is then amplified by polymerase chain reaction (PCR) followed by fragmentation and addition of sequencing adapters, typically using a hyperactive Tn5 transposase. Throughout the workflow individual cells are processed in separate wells of a PCR plate. While more labor-intensive and expensive than emulsion droplets methods, SMART-protocols enable a much deeper analysis of the full-length transcriptome, the characterization of splice isoforms, allelic variants and single nucleotide polymorphisms at single-cell resolution^6,7^. Among SMART-protocols SS2 remained for many years the preferred method thanks to its affordability, sensitivity, robustness and ease of use^8^. Recently, Hagemann-Jensen *et al.* developed Smart-seq3 (SS3), a novel SMART-protocol which includes Unique Molecular Identifiers (UMIs) for transcript counting^5^. Combining UMIs with 5’-biased tagmentation and paired-end sequencing, allows the *in silico* reconstruction of transcript isoforms at an unprecedented resolution. However, the SS3 workflow remains labor-intensive, with the cDNA enrichment and tagmentation steps requiring careful fine-tuning to reach the appropriate balance between internal and UMI-containing reads.

In this study we present “FLASH-Seq” (FS), a novel full-length scRNA-seq protocol capable of generating sequencing-ready libraries in less than 4.5 hours while offering higher gene detection than its predecessors.

## Single-Step Reaction Displays High Sensitivity

We established the first version of the FS protocol (Fig S1) by introducing several key modifications over SS2. First, we combined RT with cDNA pre-amplification in a single step (RT-PCR). Second, we replaced the Superscript™ II Reverse Transcriptase with the more sensitive and processive Superscript™ IV Reverse Transcriptase and shortened the RT incubation time. Third, we increased by 10-fold the amount of dCTP in the reaction mix to boost the C-tailing activity of SSRTIV and maximize the number of cDNA molecules undergoing template-switching reaction^9^. Fourth, we reverted to the original Template-Switching Oligonucleotide (TSO) design, by replacing the 3’-terminal Locked Nucleic Acid Guanidine (LNA-G), shown to be prone to strand-invasion^10,11^, with riboguanosine. Fifth, we removed the PCR amplification primers, as we previously observed that the oligo-(dT)_30_VN and TSO alone are sufficient to fuel both RT and PCR reactions^12^.

The FS protocol is 2 to 3.5 hours shorter than other methods (Fig 1a) and displays greater gene detection in HEK 293T cells (Fig 1b), regardless of the sequencing depth (Fig S2a). It also shows the typical full-length gene body coverage (Fig 1c) and excellent cell-to-cell correlation (Fig 1d). Interestingly, FS generates >8-times higher cDNA amounts than SS2 and SS3 at the same number of PCR cycles (Fig S2b). The initial version of the FS protocol displayed great results in more challenging cells, such as human peripheral blood mononuclear cells (hPBMCs), similar to those obtained with the SMART-Seq Single Cell Kit (SSsc, Takara Bio) (Fig 1f & Fig S2c). Of note, we observed a higher proportion of intronic reads in SS3 compared to other full-length scRNA protocols (Fig S2d), a finding also reported by Hagemann-Jensen *et al*^5^.

**Fig. 1.**
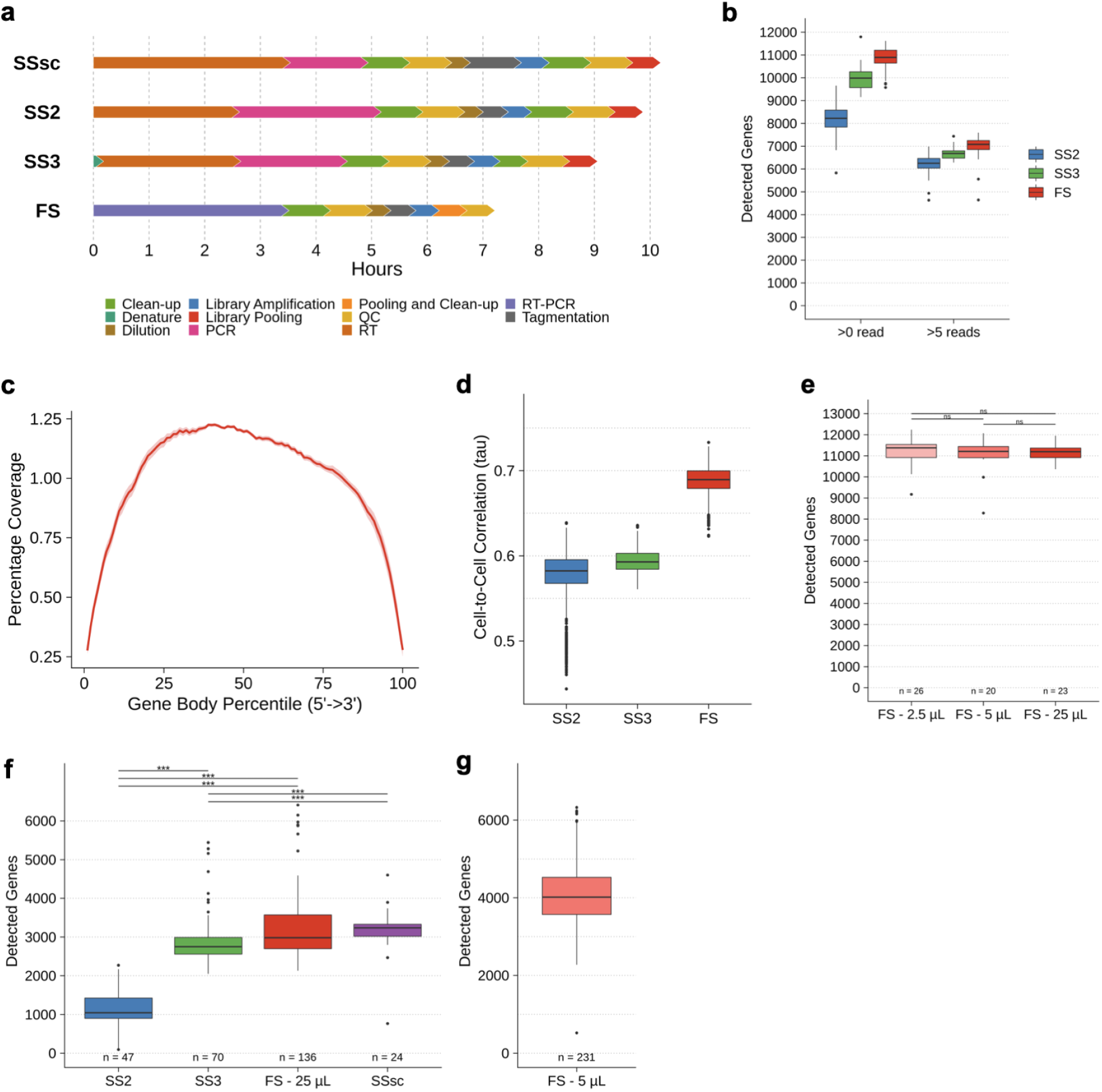
FLASH-Seq Protocol. **a.** Estimated protocol duration to process a 96-well plate of HEK 293T cells for four full-length scRNA-seq protocols. Steps are color-coded. Quality controls (QC) include concentration and size distribution measurements. **b.** Number of detected genes in HEK 293T cells processed with SS2 (n = 80 cells), SS3 (n = 42) or FS (n = 85). Gene detection threshold at two thresholds. **c.** FS gene body coverage of HEK 293T cells (n = 85). **d.** Cell-to-cell Kendall’s tau correlation between gene expressions in SS2, SS3 and FS, using genes expressed in the three methods (n = 20042). **e.** Impact of the reaction volume on the number of detected genes in HEK 293T cells (Dunn’s test, Bonferroni adj. pval > 0.05). **f.** Number of detected genes in hPBMCs processed with SS2, SS3, FS or SMART-Seq Single Cell Kit (SSsc, Takara Bio), in a 25 μL final reaction volume. Detection threshold was set to >0 read (Dunn’s test, Bonferroni adj. pval < 0.05). **g.** Impact of the reaction volume in hPBMCs. Number of detected genes in FS carried out in a final reaction volume of 5 μL (n = 231). Comparisons of gene detection in b,d,e,f,g performed on 500,000 downsampled raw reads.

## Automation and Miniaturization

We then set out to miniaturize the FS protocol to reduce costs and make it amenable to automation, with the aim of increasing reaction efficiency^13–16^. As it only requires the addition of a single reaction mix, FS is particularly suitable for automation and miniaturization, while minimizing the risk of evaporation during RT thanks to its greater starting reaction volume.

We FACS-sorted HEK 293T cells in different volumes of lysis buffer: 5 μL (96-well plates), 1 μL or 0.5 μL (384-well plates) and then dispensed 20 μL, 4 μL or 2 μL of RT-PCR mix, respectively. Decreasing the reaction volume had a limited impact on the number of detected genes compared to regular FS volume in HEK 293T cells (Fig 1e, Dunn’s test, Bonferroni adj. pval > 0.05). However, miniaturization led to a tremendous increase in the number of detected genes in hPBMCs, with FS clearly outperforming SS2, SS3 and SSsc (Fig 1f & 1g), regardless of sequencing depth (Fig S2c). We eventually settled on a 5 μL reaction volume (i.e. sorting volume of 1 μL) as sorting single cells in ≤0.5 μL requires a finer calibration of the FACS instrument, which might not be achievable by every lab. To eliminate a possible bias derived by different reaction volumes we miniaturized both SS3 and SSsc to 5 μL final (2- and 10-fold reduction over the recommended specifications, respectively) which, unfortunately, also resulted in an impaired efficiency and lower performance (Fig S3a & S3b). While these two protocols could certainly be miniaturized, further work will be needed. Since we performed this study, Smart-seq3express was published on bioRxiv and could potentially address this issue^17^.

## FS Low-Amplification

The cost reduction and automation of the miniaturized FS allowed us to test >100 new reaction conditions and additives to attempt to improve the FS protocol (extended file #1). Surprisingly, very few of them were beneficial and often displayed several limitations (e.g., increased intergenic or multi-mapped reads).

Therefore, we decided to focus our efforts on optimising the protocol duration instead. Full-length scRNA-seq protocols generally yield nanograms of cDNA after pre-amplification, which is not necessary, as libraries can be generated from sub-picogram levels of cDNA^18^. Throughout our experiments we observed a higher efficiency for FS compared to SS2 or SS3 (e.g., Fig S2c). To test the limits of our system, we varied the number of pre-amplification cycles (4, 6, 8, 10, 12), corresponding enrichment PCR cycles post-tagmentation (24, 22, 18, 16, 14) and titrated the amount of Tn5 used for each condition (Fig S4). Due to the minute amounts of cDNA in the reaction at this stage, the purification, intermediate quality control and dilution steps were skipped, saving ~2.5 additional hours. This resulted in a comparatively cheaper protocol requiring limited hands-on time (<30 minutes), all while minimizing losses derived from sample cleanup. We called this faster version, FLASH-Seq Low-Amplification (FS-LA) (Fig 2a).

**Fig. 2.**
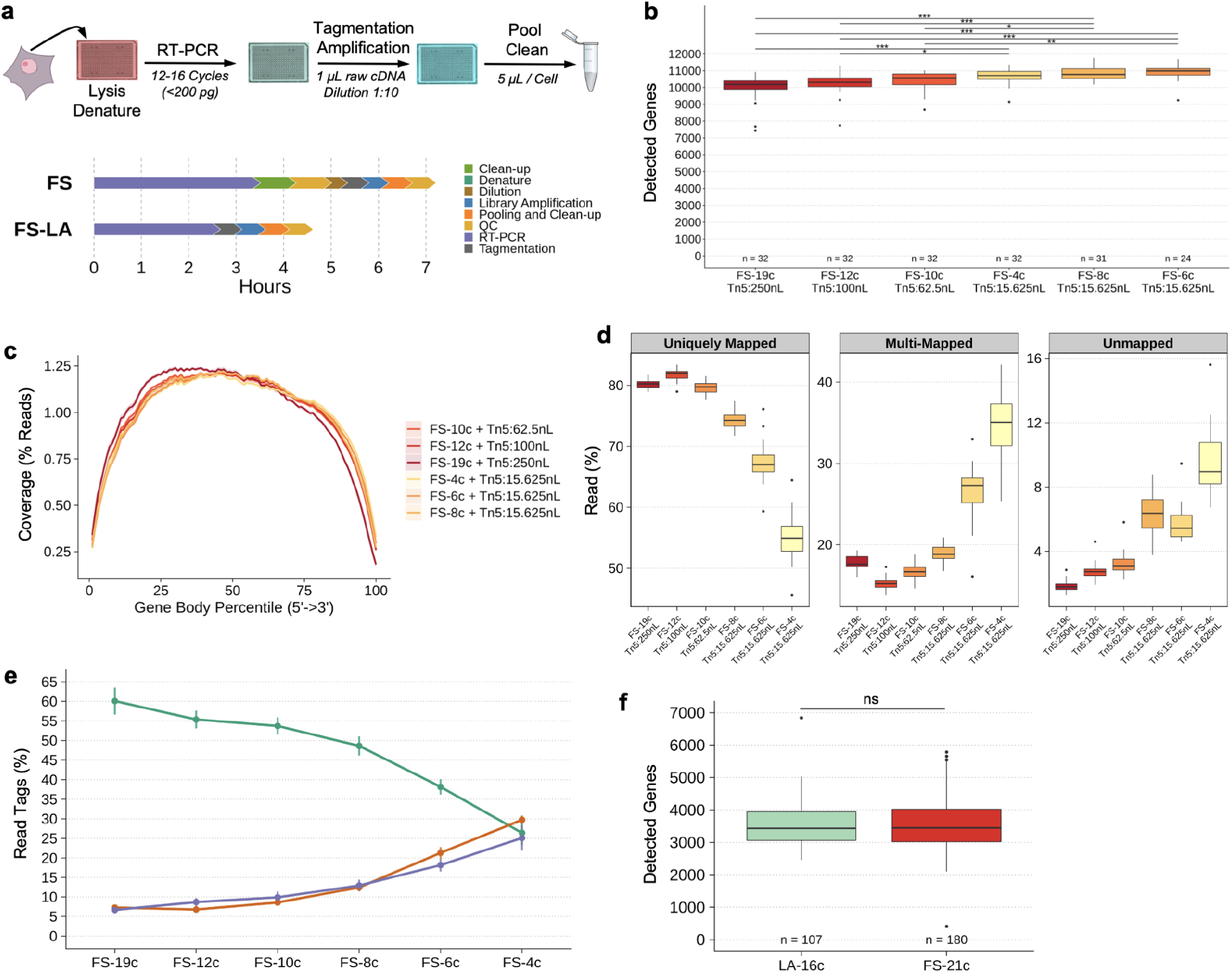
Overview of FS Low Amplification (FS-LA) protocol. **a.** Schematic representation of FS-LA and FS protocols. Steps are color-coded. QC includes concentration and length distribution measurements. **b.** Number of detected genes in HEK 293T cells processed with FS (19 PCR cycles) or FS-LA (4 to 12 PCR cycles), using 250,000 downsampled raw reads (Dunn’s test, Bonferroni adj. pval < 0.05). Detection threshold was set to >0 read. **c.** Gene body coverage of FS and FS-LA protocols (4 to 12 PCR cycles). **d.** Mapping statistics. Panels depict the percentage of uniquely mapped, multi-mapped or unmapped reads for FS and FS-LA protocols (4 to 12 PCR Cycles). **e.** Variation of the proportion of read mapped to exonic, intronic or intergenic features measured using ReSQC, in read tag percentages. **f.** Number of detected genes in hPBMCs processed with FS (21 PCR cycles) or FS-LA (16 PCR cycles), using 250,000 downsampled reads (Mann-Whitney U Test, pval > 0.05). Detection threshold was set to >0 read.

Exploration of FS-LA libraries revealed an increasing number of detected genes associated with the lower number of cycles compared to FS (19 cycles) (Fig 2b) and great gene coverage (Fig 2c & S5). However, we also observed that a lower number of cycles resulted in a higher percentage of intergenic, intronic, multi-mapped and unmapped reads (Fig 2d & 2e). The amount of Tn5 used for tagmentation had little impact on the results (Fig S6). A careful analysis of the data in Integrated Genome Viewer revealed that when performing <10 cycles these intergenic reads were randomly spread across the genome, including in centromeres (Fig S7). We hypothesized that they may be the result of gDNA tagmentation and that their abundance would be inversely proportional to the number of pre-amplification PCR cycles. In fact, the percentage of intergenic reads remained stable between ~10 and 19 pre-amplification cycles, while still showing excellent gene detection (Fig S6). Interestingly, the intergenic reads did not correlate with the number of detected genes (Fig S8a) but were inversely proportional to the percentage of uniquely mapped reads when using <10 PCR cycles (Fig S8b). These findings suggest that once a critical amount of cDNA is reached, high-quality scRNA-seq libraries can be prepared using the RT-PCR product directly, even with few PCR cycles. In the case of HEK 293T cells, FS-LA decreased the protocol duration to <4.5 hours (Fig 2a).

The RNA content of a cell determines the number of PCR cycles that are needed. Large cells such as HEK 293T contain ~16 pg RNA and as few as 10 PCR cycles are generally sufficient to generate high-quality libraries (Milteny, Average RNA yields). On the other hand, hPBMCs typically contain 2-4 times less RNA and might require more amplification. In the present study we decided to perform 16 pre-amplification and 14 enrichment PCR cycles to minimize the risk of gDNA contamination, although a lower number of cycles will most likely guarantee usable results as well. The resulting libraries had a similar number of detected genes as FS (21 cycles) (Fig 2f) while harboring minimal percentages of intergenic or unmapped reads (Fig S9). The lower number of PCR cycles did not affect the cell type assignment (Fig. S10).

We conclude that the FS protocol can be performed using a low number of cycles, reducing the experiment duration to a single afternoon while preserving comparable data quality as longer experiments. As in all single-cell experiment, a pilot experiment is recommended to determine the exact number of cycles and avoid cDNA under-amplification, as described in extended file #2.

## Modular FS

The robustness and adaptability of the FS protocol are advantages it shares with SS2^19–21^. It can therefore be expected that FS buffers and concepts will be easily implemented in existing and upcoming protocols. To illustrate this concept, we modified FS by including UMI into the TSO sequence, as previously described by SS3^5^, thus generating both 5’ UMI-containing reads and internal reads without UMI.

When designing our UMI-TSO, we hypothesised that the addition of 8-random nucleotides upstream of the 3 ribo-G (i.e. SS3 TSO) may favor strand-invasion, as it has been described in similar settings with the nanoCAGE TSO^11^. This occurs when the TSO anneals to an internal sequence of the cDNA or mRNA molecule (instead of at the 5’ end of the cDNA) that, by chance, displays sequence complementarity (Figure S11). These artefacts could affect isoform detection and bias gene counts. We therefore compared the use of SS3 TSO and FS-UMI-TSO - both with a -NNNNNNNN-rGrGrG - to seven additional TSO with the UMI separated from the ribo-Gs by a different spacer sequence in a similar fashion to the nanoCAGE protocol^11^ or the 10x 5’ kit v3 (Fig 3a). These new TSOs were tested in combination with STRT-seq2i-oligo-dT^22^, processing 16 HEK 293T cells per condition. A subset of these TSOs were also tested with SS3-^5^ or FS- oligo-dT_30_VN (full list of evaluated combinations in Fig S12). Interestingly, the cDNA yield was strongly influenced by the TSO (Fig S13). The proportion of 5’ UMI-reads varied from 10.3% (SS3) to 43.7% (TSO-ATAAC+STRT-dT) (Fig 3b), highlighting the difficulty in striking the right balance between the two read types ahead of sequencing. Considering these differences, we also included HEK 293T cells data generated by Hagemann-Jensen *et al*^5^ (E-MTAB-8735, ‘HEK 293T - 0.5 μM Forward Primer’) for comparison.

**Figure 3.**
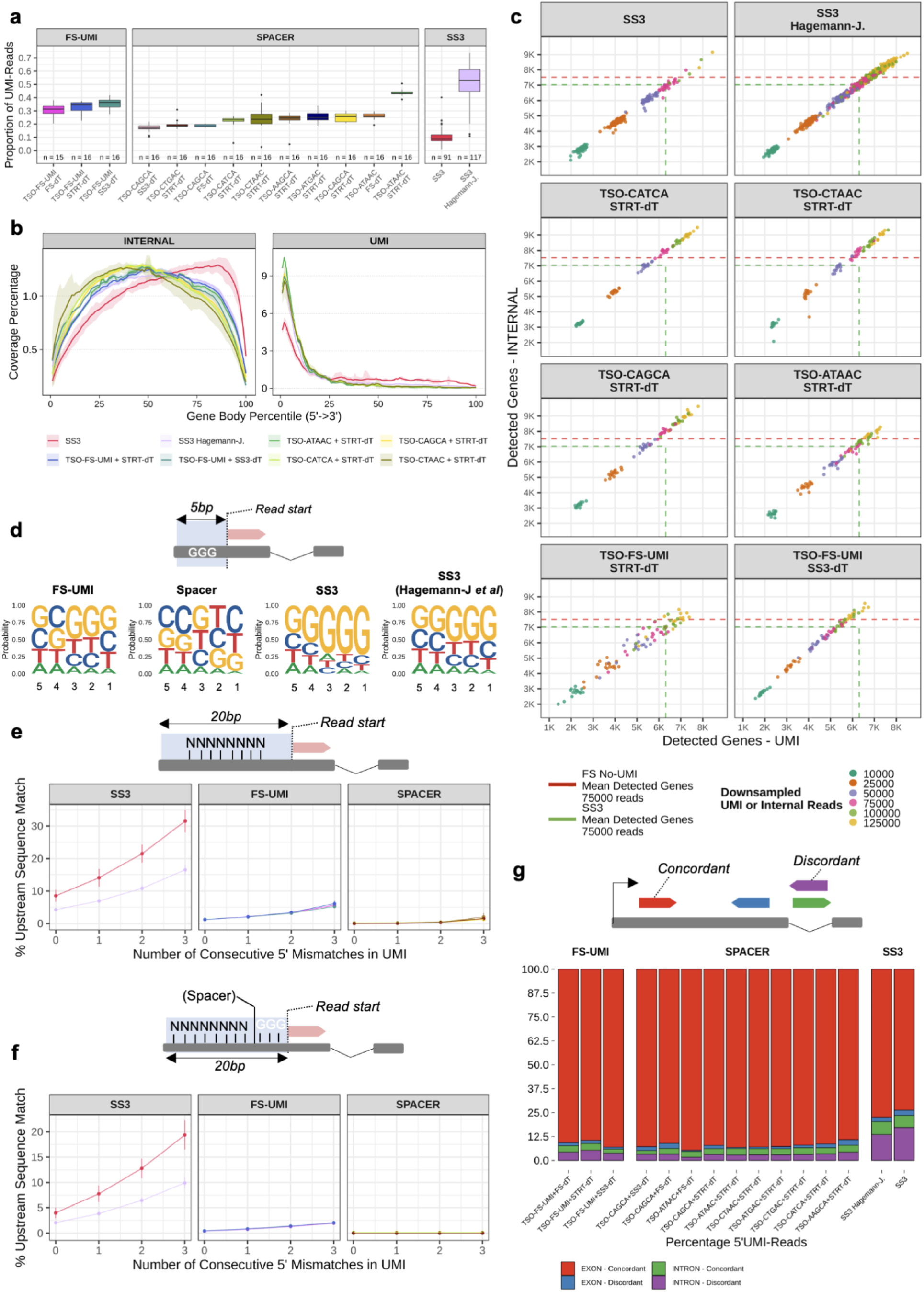
Addition of UMI to FS protocol and evaluation of oligo-dT - TSO combinations. **a.** Proportion of UMI-reads among the different combinations of TSO-UMI / oligo-dT. **b.** Gene body coverages. **c.** Relationship between the number of detected genes using UMI and internal reads. UMI and internal reads were downsampled to 10K, 25K, 50K, 75K, 100K, 125K reads. Dashed red line represents the mean number of detected genes using FS (n = 85) downsampled to 75K raw reads. Dashed green lines represent the mean number of detected genes from Hagemann-Jensen *et al.* using UMI or internal reads, downsampled to 75K raw reads. **d.** Nucleotide distribution in the 5-bp adjacent to the read start using 500K randomly selected reads from each subgroup. **e.** Percentage of deduplicated 5’ UMI reads harboring a match between the UMI and the upstream sequence within 20-bp, with 0 mismatches or 1 to 3 consecutive 5’ mismatches. **f.** Same as e. but searching for UMI+GGG or UMI+SPACER+GGG motifs. **g.** Reads distribution on exonic / intronic sequences of protein coding genes (n = 20013) accounting for the read and gene orientations, in percentage of unambiguously assigned reads. Remaining samples for b. and c. figures are depicted in Fig S14.

Overall, all conditions displayed great mapping statistics (Fig S14) and coverage (Fig 3b & Fig S15). The addition of a spacer sequence increased by up to 15% (TSO-CATCA) the number of detected genes compared to SS3 (Fig 3d & Fig S15). Six out seven TSOs with a spacer even outperformed the regular FS in respect of the number of detected genes. As hypothesized, we observed several elements suggestive of strand-invasion events when using UMIs upstream of the ribo-Gs motif (Fig S16). First, a ‘GGG’ motif was more often observed adjacent to the first base of deduplicated 5’ UMI reads (Fig 3d). Second, a perfect match between the UMI and the sequence upstream of the read was observed in >4.25% of SS3 deduplicated 5’ UMI reads (Fig 3e). When accounting for the partial 5’ match of the UMI and upstream sequence, this value rose to >10.9% with two mismatches (Fig 3e). The percentage of matches remained higher in SS3 data even when considering the presence of the ‘GGG’ motif and spacer sequence (Fig 3f). Third, SS3 showed an increased percentage of UMI reads mapping to intronic features and a decrease in 5’ UTR reads compared to FS (Fig S17). Fourth, an important proportion of SS3 deduplicated 5’ UMI reads mapping to intronic regions of protein coding genes were mapped in antisense orientation compared to the gene reference (Fig 3g). The addition of a spacer sequence between the ribo-Gs and the UMI appeared to prevent most strand-invasion events associated with the UMI, regardless of the spacer sequence. In addition, the presence of the spacer facilitated the detection of the remaining events (Fig S18).

These results highlight the versatility of FS and confirm that UMIs can also be used in our RT-PCR setup. They also suggest that better TSO sequences can still be designed and highlight the importance of spacer sequences in UMI-TSO design.

## Discussion

In this study we introduced FLASH-Seq, a new full-length single-cell RNA-sequencing method which is faster and more sensitive than the existing single-cell protocols, thanks to a novel combination of additives and enzymes. FS is a robust protocol that can be easily upgraded, as demonstrated by the addition of UMIs to count individual molecules^5^. To decrease protocol duration we aimed at generating just enough cDNA to directly use it in the tagmentation reaction, removing the need for sample cleanup. We could then combine the superior gene detection of FS with a much faster and easier workflow, capable of generating sequencing-ready libraries in half a day of work. Thanks to its ease of use, rapid execution and automation-friendly design, we expect FS-LA to be particularly useful when processing a large number of samples, i.e. in genomics core facilities or single-cell atlases. While this manuscript was ready for submission a new method called Smart-seq3xpress was posted on bioRxiv^17^. Although Smart-seq3xpress and FS-LA share several similarities, we believe that the addition of oil to the reaction as well as the small reaction volumes described in Smart-seq3xpress make the protocol less user-friendly and more difficult to carry out without dedicated nanodispensers, as the pipetting volumes fall below the specs of most liquid handling robots. Moreover, the duration of the RT reaction in Smart-seq3xpress is more than twice as long as FS and the PCR mix has to be added in a separate step. While manageable when working on a few cells, these additional steps will have a strong impact when processing a large number of samples in parallel.

Compared to older or commercial alternatives, both FS and SS3 delivered excellent results at an affordable cost/cell (SS2, ~7$/cell; SS3, <$1; FS, ~$1; SSsc, ~$120). However, we observed that strand-invasion events occurred more often when the UMI was located upstream of the ribo-G stretch in the TSO. The excess of discordant intronic reads in SS3 5’ UMI reads suggests that strand-invasion could happen preferentially during the reverse transcription but we cannot formally exclude other mechanisms. For instance, SS3-TSO is able to prime the cDNA in the enrichment PCR reaction^5^. Strand-invasions could therefore occur during PCR even in the presence of an excess of competing forward primers. However, the low melting temperature of the UMI-rGrGrG binding sequence should make this event less likely, especially as the annealing temperature in the PCR is 65-67°C. The exact level of strand-invasion is difficult to estimate due to the diversity of underlying mechanisms and could vary from one sample to the other. In this study, strand-invasion events varied from ~1.2% of the deduplicated 5’ UMI reads in FS-UMI to >4% in the SS3 samples we analysed. Due to their non-stranded and fragmented nature, the impact of strand-invasion on the internal reads could not be evaluated. These results raise concerns about generation of false positives in isoform detection and biased gene counts when using TSOs with UMI located upstream of the ribo-Gs. The addition of a spacer sequence isolating the UMI from the ribo-Gs reduced the risk of strand-invasion, displayed higher cDNA yield and superior gene detection. We therefore recommend the use of a spacer when including UMI to the TSO sequence in combination with FS buffers for superior gene detection and a significant decrease in strand-invasion events.

In conclusion, we believe that FLASH-Seq can become the tool of choice when looking for an efficient, robust, modular, affordable and automation-friendly full-length scRNA-seq protocol.

## Material and Methods

### Single-cell isolation

Single hPBMC were obtained from healthy human donors after signing the informed consent form. Leucocytes were isolated from whole blood by density gradient centrifugation using Ficoll-Paque (Sigma). HEK 293T cells were cultivated in Dulbecco’s Modified Eagle Medium (DMEM, ThermoFisher Scientific) supplemented with 10% FBS and 1% penicillin/streptavidin. Both hPBMCs and HEK 293T cells were cryopreserved in FBS + 10% DMSO (Sigma-Aldrich). Prior to FACS sorting, hPBMCs were resuspended in 1x PBS + 2% FBS and stained with a FITC-conjugated Mouse Anti-Human CD45 Antibody (Clone HI30, BD Biosciences) for 25 min on ice. Unbounded antibodies were washed away by adding 3 mL of Roswell Park Memorial Institute 1640 medium (RPMI, ThermoFisher Scientific) + 2% FBS. Prior to FACS-sorting both hPBMCs and HEK 293T cells were centrifuged for 5 min at 300 × g, resuspended in 1x PBS + 0.05% BSA, and strained through a 40-μm filter (pluriSelect) then stained for 5 min with Propidium Iodide (1 mg/mL, ThermoFisher Scientific) at room temperature to label dying cells.

Single cells were sorted on a FACSAria Fusion (BD Biosciences) with a 100-μm nozzle, using the following strategy: particles smaller than cells (debris) were eliminated with an area plot of Forward-Scatter (FSC-A) versus Side-Scatter (SSC-A) by gating for cell-sized particles inside the gate. A 2-steps doublet discrimination was carried out to remove cell aggregates by making plots of Height versus Width, both in the Forward- and Side-Scatter channel, FSC-H vs FSC-W and SSC-H vs SSC-W, respectively. For the hPBMCs we plotted FITC-A (for CD45) vs TexasRed-A (for PI), sorting all singlets that were FITC+PI-. For HEK293T cells we plotted FSC-A vs TexasRed-A (for PI), sorting all singlets that were PI-.

Cells were sorted in different volumes of lysis buffer, according to the plate type: in 5 μL, when using 96-well plates; 1 or 0.5 μL, when using 384-well plates. For all experiments we used LoBind twin.tec plates (Eppendorf). Lysis buffer was distributed with the I.DOT instrument (CellLink) ahead of sorting and plates were stored at −20℃ until needed.

### Lysis buffer composition

For the sake of brevity only the volumes used for sorting cells in 96-well plates (5 μL/well) are indicated. When miniaturising the protocol we decreased the volume of each reaction accordingly.

- Standard Triton Buffer (0.2% Triton final): 0.1 μL Triton X-100 (10% v/v, Sigma-Aldrich), 1.2 μL dNTP mix (25 mM each, Roth), 0.46 μL oligo-dT_30_VN (5’Bio-AAGCAGTGGTATCAACGCAGAGTACT_30_VN-3’; Bio = C6-linker biotin; 100 μM, IDT), 0.15 μL Recombinant RNAse Inhibitor (40 U/μL, Takara Bio), 0.06 μL Dithiothreitol (DTT, 100 mM; part of the Superscript IV kit, ThermoFisher Scientific), 1 μL betaine (5M, Sigma-Aldrich), 0.45 μL dCTP (100 mM, ThermoFisher Scientific), 0.46 μL Template-Switching Oligonucleotide (TSO, 5′ Bio-AAGCAGTGGTATCAACGCAGAGTACrGrGrG-3′, Bio = biotin with C6-spacer; 100 μM, IDT) and water to 5 μL final.
- Standard Triton Buffer (0.2% Triton final) with lower oligo-dT_30_VN: only 0.092 μL (⅕ compared to the buffer indicated above) or 0.046 μL (1/10) oligo-dT_30_VN were used for hPBMC to minimize dimer artefacts. The other components are the same as in the Standard Triton Buffer.
- BSA-Triton Buffer (1 mg/mL BSA + 0.2% Triton): 0.5 μL BSA (10 mg/mL, ThermoFisher Scientific) were added to the Standard Triton Buffer described above.
- BSA Buffer (1 mg/mL BSA final): 0.5 μL BSA (10 mg/mL, ThermoFisher Scientific) were added to the Standard Triton Buffer described above but in this case Triton was not included.
- Guanidine-based Buffer (250 mM final): 0.03125 μL guanidine hydrochloride (8 M, Sigma-Aldrich) were added to the Standard Triton Buffer described above but in this case Triton was not included.

After cell sorting the plates were sealed and immediately stored at −80°C until ready for further processing. Lysed cells can be stored in these conditions for several months without any appreciable decrease in RNA quality. Freeze-thaw cycling should be avoided.

### RT-PCR reaction

Plates were taken out of the −80°C freezer, immediately transferred to a Mastercycler thermocycler (Eppendorf), incubated either for 3 min at 72°C or for 3 min at 95°C, then immediately transferred to a metal block place on an ice bucket. RT-PCR mix was added using the I.DOT instrument (Dispendix). When sorting cells in 5 μL of the lysis buffer and using 96-well plates, 20 μL RT-PCR mix were added. In the miniaturization experiments the volumes were scaled down accordingly, i.e. adding 4 μL RT-PCR mix when cells were sorted in 1 μL lysis buffer and 384-well plates (or 2 μL RT-PCR mix when cells were sorted in 0.5 μL lysis buffer). The RT-PCR mix had the following composition: 1.19 μL Dithiothreitol (DTT, 100 mM; part of the Superscript IV kit, ThermoFisher Scientific), 4 μL betaine (5 M, Sigma-Aldrich), 0.23 μL magnesium chloride (1 M, Ambion), 0.48 μL Recombinant RNAse Inhibitor (40 U/μL, Takara Bio), Superscript IV Reverse Transcriptase (200 U/μL, ThermoFisher Scientific), 12.5 μL KAPA HiFi Hot-Start ReadyMix (2x, Roche), nuclease-free water to a final volume of 20 μL. The plate content was briefly vortexed and spun down, before starting the following program: 60 min at 50°C, 98°C for 3 min, then N cycles of (98°C for 20 sec, 67°C for 20 sec, 72°C for 6 min). The number of cycles to the cell type: when using HEK 293T cells we generally used N=19. When using hPBMC we used N=21. The following modifications were tested during protocol optimization:

- Short RT: the RT reaction was shortened to 30 min;
- Short extension: in each PCR cycles the extension step was shortened to 4 min;
- Low-Amplification-FLASH-Seq (see below for details): when optimizing the “Low-Amplification” protocol we tested N=4, N=6, N=8, N=10, N=12, N=16. These experiments were carried out only in 384-well plates, sorting cells in 1 μL of Standard Lysis Buffer and adding 4 μL of the RT-PCR mix described above.

Several additives, enzymes and buffers were also tested, alone or in combination:

- T4 gene 32 protein (10 mg/mL, NEB or EPFL “homemade” T4g32p, 10 mg/mL). In 96-well plates, we added per reaction either 5 μg, 3.75 μg, 2.5 μg, 1.25 μg or 0.625 μg. As recommended by the manufacturer, the RT reaction was performed at 37°C when using T4 gene 32 protein.
- Higher concentrations of TSO. In the attempt of increasing the number of cDNA molecules which could successfully be amplified in the following pre-amplification reaction, we supplemented the RT-PCR mix with extra TSO, testing 1.5x, 2x, 3x, 4x and 5x increase over the standard protocol.
- A combination of KAPA HotStart ReadyMix and Pfu DNA Polymerase (2 U/μL, Promega) was used in some tests to try to increase enzyme processivity. In all those tests we used 0.5 μL Pfu/ 25 μL reaction.
- Ficoll-400 (20% w/v, Sigma-Aldrich) was tested as a macromolecular crowding agent to a final concentration of 4%. Due to volume constraints, including Ficoll-400 in the reaction required betaine to be removed.
- Maxima H- Reverse Transcriptase (200 U/μL, ThermoFisher Scientific) replaced Superscript IV in some tests, all the other components of the standard RT-PCR mix remaining the same.
- Addition of 20 mM, 30 mM or 40 mM of NaCl (Sigma-Aldrich) to the RT-PCR mix.
- Extreme Thermostable Single-Stranded DNA Binding Protein (ET SSB, 500 ng/μL, NEB) was tested in 4 different amounts: 40 ng protein/reaction, 80 ng, 160 ng or 250 ng.

### Sample cleanup

Experiments carried out in 96-well plates or 384-well plates were cleaned-up using a Fluent 780 Automated Workstation (Tecan, Switzerland). SeraMag SpeadBeads™ (GE Healthcare) containing 18% w/v Polyethylene glycol MW=8000 (Sigma-Aldrich) were used. A detailed protocol is described elsewhere^12^. Amplified cDNA cleanup was not performed for all experiments of this study. When using the FS-LA protocol, cells were sorted and processed in 384-well plates (5 μL final volume after RT-PCR mix addition) and 1 μL was used for direct tagmentation without cleanup.

When performing sample cleanup, unless otherwise indicated, we skipped ethanol wash while the cDNA was immobilized on magnetic beads, to avoid material losses and simplify the workflow. Purified cDNA was generally eluted in 10 or 15 μL nuclease-free water.

### Sample QC

Sample concentrations were quantified by using the Quant-iT™ PicoGreen™ dsDNA Assay Kit (ThermoFisher Scientific) and performed in black Nunc™ F96 MicroWell™ Polystyrene Plates (ThermoFisher Scientific), according to the manufacturer’s instructions, except that all reaction volumes were cut by half to reduce costs. Fluorescence measurements were recorded on a Hidex Sense instrument (Hidex). In order to assess cDNA size distribution, 11 samples from each plate were randomly chosen, loaded on a High Sensitivity DNA chip and run on a 2100 Bioanalyzer System (Agilent Technologies). As a rule of thumb, the criteria for deciding whether an experiment was worth to be followed up were the following: average size distribution around 1.5-2.0 Kb, low proportion of short fragments (<400 bp) and absence of residual primers/primer dimers (<200 bp).

### Sample normalization

Samples were often too concentrated for the following tagmentation reaction and required normalization. The concentration returned by PicoGreen™ measurement was used as input for normalizing each sample to a final concentration of 150 pg/μL. Briefly: one microliter of purified cDNA was transferred to a new 96- or 384-well plate by using the Tecan Fluent robot and transferred to the I.DOT instrument, where different amounts of nuclease-free water were added to reach the desired concentration. This plate represented our working dilution cDNA plate, which was used in the following library preparation step.

### Standard tagmentation and Enrichment PCR

Tagmentation was carried out using 3 different versions of the Tn5 transposase:

- Library preparation with the Nextera^®^ XT kit (Illumina). Briefly: 1 μL from each well of the working dilution cDNA plate was transferred to a new plate, before adding 1 μL of Amplicon Tagment Mix (ATM) and 2 μL of Tagment DNA Buffer (TD). The plate content was gently vortexed, spun down and incubated for 8 min at 55°C. The Tn5 was released from the tagmented DNA by adding 1 μL of Neutralization Buffer (NT), followed by a 5 min incubation at room temperature. We then added 3 μL of Nextera PCR Master Mix (NPM) and 2 μL of pre-mixed N7xx and S5xx Nextera v2 Index Primers (sets A, B, C, D). Given the low amount of cDNA used for the tagmentation we pre-diluted each vial of Nextera primers 1:5 using low-EDTA TE buffer (10 mM Tris-HCl, 0.1 mM EDTA, pH 8.0), before generating four 96-well plates, with each well containing a unique combination of N7xx + S5xx (layout available upon request). The enrichment PCR reaction was carried out as follows: 72°C for 3 min, 95°C for 30 sec, then 14 cycles of (95°C for 10 sec, 55°C for 30 sec, 72°C for 30 sec), 72°C for 5 min, 4°C hold.
- Library preparation using homemade Tn5 transposase (EPFL). The Tn5 synthesis used the pTXB1 plasmid, available from Addgene (#60240). The original protocol and further modifications were used for protein preparation^18,23^. The tagmentation reaction was carried out as follows: 1 μL from each well of the working dilution cDNA plate was transferred to a new plate, before adding 1 μL of 100% dimethylformamide (DMF, Sigma-Aldrich), 1 μL of 5X TAPS buffer (50 mM TAPS-NaOH pH 7.3 at 25°C, 25 mM MgCl_2_; 0.22 μm filtered), 0.025 μL Tn5 transposase (~21 μM) and nuclease-free water to a final volume of 5 μL. The plate content was gently vortexed, spun down and incubated for 8 min at 55°C. The Tn5 was released from the tagmented DNA by adding 1.25 μL of 0.2% SDS, followed by a 5 min incubation at room temperature. The Enrichment PCR Mix added in the following step had the following composition: 0.25 μL of KAPA HiFi Enzyme, 0.375 μL dNTP mix (10 mM), 2.5 μL KAPA HiFi Buffer (5x) (all part of the KAPA HiFi kit, Roche), and water to a final volume of 3.75 μL. Similarly to what described above, we then added 2.5 μL of pre-mixed N7xx + S5xx Index Primers. However, this time we used a custom set of 32 N7xx and 48 S5xx (IDT, sequences and plate layout available on request). All primers were diluted to 10 μM upon receipt by using low-EDTA TE buffer and then mixed in equal volumes on 96-well or 384-well plates by using the I.DOT instrument, so that each one of them had a working concentration of 5 μM. The enrichment PCR reaction was carried out as described in the previous section.
- plexWell™ Rapid Single Cell kit (seqWell). This time the purified cDNA was diluted to a final concentration of 250 pg/μL and 4 μL (corresponding to 1 ng cDNA) were used as input. We followed the manufacturer’s recommendations and transferred 4 μL from each sample to a different well of the SBL96 Plate to carry out the Sample Barcode reaction. The tagmentation reaction was carried out at 55°C for 30 min, before adding 6 μL of X Solution for quenching. The plate was then incubated at 68°C for 20 min. Nine microliter solution from each well of the SBL96 plate were pooled in a 1.5-mL Eppendorf tube, 1 volume of MAGwise Paramagnetic Beads were added and the tagmented cDNA was eluted in 40 μL of 10 mM Tris-HCl pH 8. Five microliters of PB Reagent (Pool Barcode) were added alongside with 3X Coding Buffer and a second tagmentation was carried out at 55°C for 15 min. The reaction was quenched again with 33 μL X Solution, incubated at 68°C for 10 min and purified again with 1X MAGwise Paramagnetic Beads. The cDNA was eluted from the beads in 23 μL of 10 mM Tris-HCl pH 8, followed by the addition of 4 μL of Library Primer Mix and 27 μL of KAPA HiFi HotStart ReadyMix. The enrichment PCR reaction was carried out as follows: 72°C for 3 min, 95°C for 3 min, then 12 cycles of (98°C for 30 sec, 64°C for 15 sec, 72°C for 30 sec), 72°C for 3 min, 4°C hold. The final library was then purified using a 0.8x of MAGwise Paramagnetic Beads.

### Tagmentation of “Low-Amplification” samples

In the “Low-Amplification” protocol, several aspects of the workflow had to be adjusted, depending on the number of pre-amplification PCR cycles used. Hereafter some general guidelines are described but appropriate reaction conditions need to be adjusted according to the RNA content of each cell type.

For HEK 293T cells we performed the RT-PCR reaction only in 384-well plates and in a final volume of 5 μL. We then used 1 μL of unpurified amplified cDNA as input for the tagmentation reaction. In pilot experiments we noticed that leftover primers, dNTPs and salt can completely inhibit the activity of the Tn5 transposase (data not shown). An acceptable compromise was to dilute the cDNA 10 times, i.e. adding 1 μL and 9 μL tagmentation mix to a final volume of 10 μL. The associated cost of such a large volume is negligible when using homemade Tn5 transposase.

We recommend users to perform a titration experiment in order to determine the most appropriate amount of Tn5 to use to achieve the desired library size. Guidelines can be found in extended file #2.

### Sample pooling and sequencing

An aliquot from each well of the plate was pooled together, transferred to a 1.5 mL LoBind tube (Eppendorf) and cleaned up with a 0.8:1 SeraMag SpeadBeads™ containing 18% w/v Polyethylene glycol MW=8000. Elution was performed in nuclease-free water and 1 μL was used for measuring the concentration with the Qubit™ dsDNA High Sensitivity Assay Kit (ThermoFisher Scientific), while the size was assessed with a High Sensitivity DNA chip on an 2100 Bioanalyzer.

Libraries prepared with different sets of index primers were pooled together and sequenced using a NextSeq™ 500/550 High Output Kit v2.5 (75 Cycles) with read mode 75-8-8-0. FS-UMI libraries were sequenced using NextSeq™ 500/550 Mid Output Kit v2.5 (150 Cycles) with read mode 100-8-8-50.

### Other single-cell methods

Smart-seq2 libraries were generated according to Picelli *et al*^12^. Smart-seq3 libraries were generated using the latest version of the protocol (V3) on protocol.io (dx.doi.org/10.17504/protocols.io.bcq4ivyw). SMART-Seq Single Cell Kit (Takara Bio) libraries were prepared following the manufacturer instructions, with the exception that all reaction volumes were halved.

### Cell-to-cell correlations

Cell-to-cell correlations were calculated using Kendall’s tau rank correlation to better handle ties (pcaPP, v1.9-74). Only genes expressed in SS2, SS3 and FS were used to calculate correlations (n = 20042).

### Single-Cell Data Analysis

Nextera and FS/SS2/SS3 adapters were trimmed using BBDUK (BBMAP, v38.86) in all reads. Umi-tools (v1.1.0) was used to extract UMI from 5’ UMI-reads and when available to trim spacer sequence, FS/SS2/SS3 adapter and GGG motifs originating from the TSO. Trimmed reads >60bp (SS2, SSsc, FS) or >50bp (SS3, FS-UMI) were aligned against hg38 (ensembl v100) using STAR (*--limitSjdbInsertNsj 2000000 --outFilterIntronMotifs RemoveNoncanonicalUnannotated*, v2.7.3) in single-pass mode and assigned to a genomic feature from Gencode v34 annotation (primary_assembly) using featureCounts (*-t exon -g gene_name*, v1.6.5). Downsampling were performed on the FASTQ files using seqkt (*-s42*, v1.3-r106). Gene body coverage and genome-wide read distribution were both estimated using ReSQC (v3.0.1).

When analysing the impact of UMI on FS-UMI/SS3, internal reads were separated from the 5’ UMI-reads (detected with umi-tools) using *filterbyname.sh* (BBMAP, v38.86) and mapped separately as described above. Internal/UMI reads with a MAPQ <5 were removed using samtools (v1.9). UMI-containing reads were assigned to a feature using featureCounts (*-t exon -g gene_name -s 1 --fracOverlap 0.25*). UMI were counted per gene using *umi_tools counts* (v1.1.0). As SS3 data from Hagemann-Jensen *et al* were sequenced using single-end reads, only read 1 of FS-UMI was used for the comparisons. All resulting data were parsed using R (v3.6.1). Marker genes and cell types were explored in hPBMCs using scran (v1.14.6) and scater (v1.14.6). Only cells with >100,000 raw reads and <25% unmapped reads were selected. Mitochondrial, ribosomal, MALAT1 and lowly expressed (rowSums < 10) genes were excluded from the analysis (n_total_genes_=21408). Two extra cells were removed due to low quality (*quickPerCellQC*). Libraries were scaled using *quickCluster/calculateSumFactors* and log-normalized. The per-gene mean-variance was modelized with *modelGeneVar*. PCA dimension reduction was performed using the top 10% most highly variable genes, selecting the first 10 components. Uniform Manifold Approximations and Projections were computed on the PCA components. Distinct clusters were identified by building a k-nearest neighbors graph (*buildSNNGraph*, k=6) and identifying individual communities with the louvain algorithm (*igraph*, v1.2.6) Cell types were assigned using commonly described hPBMCs markers: B-Cells (CD79A, JCHAIN, MS4A1, IGHM, IGHD), CD4+ T-cells (CD4, IL7R, CD3, PRDM1), CD8+ T-cells (CD8, PRF1, GZMA, CD3, PRDM1), NK-cells (FCGR3A, NKG7, GNLY, NCAM1), Naïve T-cells (CCR7, NOG, LRRN3, CD3, LEF1, TCF7), CD14+ Monocytes (CD14, LYZ), Non-classical monocytes (FCGR3A, CDKN1C).

### Strand-invasion

A GTF file regrouping the collapsed exonic and intronic genomic sequences of protein coding genes (n = 20016) was created using custom R scripts (GenomicRanges v1.38). Overlapping exonic-intronic sequences were considered as exonic. This GTF was used to unambiguously assign mapped 5’ UMI reads to exonic or intronic features in an unstranded manner with featurecounts (-s 0 --largestOverlap --fracOverlap 0.25 -R BAM). The annotated BAM file was parsed using custom R scripts to retrieve the read position, read strand, UMI sequence and gene annotation. The sequence adjacent to the read start (20-bp) was extracted using *getSeq* (BSgenome v1.41, hg38) accounting for the read orientation. To avoid overcounting PCR duplicates, UMI reads harboring the same UMI sequence, mapping position and adjacent read-start sequence were collapsed (= deduplicated 5’ UMI reads). The adjacent sequence was compared to the UMI looking for a perfect or partial match in the presence or absence of an additional ‘GGG’ or ‘SPACER-GGG’ motif. A schematic description of the strand-invasion analysis is depicted in Fig S19. Seqlogo graphics were produced using ggseqlogo (v0.1).

### Data Availability

All data supporting this study will be uploaded to a public repository upon publication.

### Code Availability

Scripts used to process the data will be soon available at: https://github.com/Radek91/FLASH-Seq.

## Acknowledgements

We thank Dr. Kevin Lau and Dr. Florence Pojer from the Swiss Federal Institute of Technology Lausanne for providing the homemade Tn5 transposase and T4 gene 32 protein.

## Contributions

V.H. developed the reaction chemistry, generated scRNA-seq libraries, performed computational analysis, prepared figures and wrote the manuscript text. D.P. generated scRNA-seq libraries. C.C. advised on the data analysis. S.P. conceived the method, planned and supervised work, developed the reaction chemistry, generated scRNA-seq libraries and wrote the manuscript.

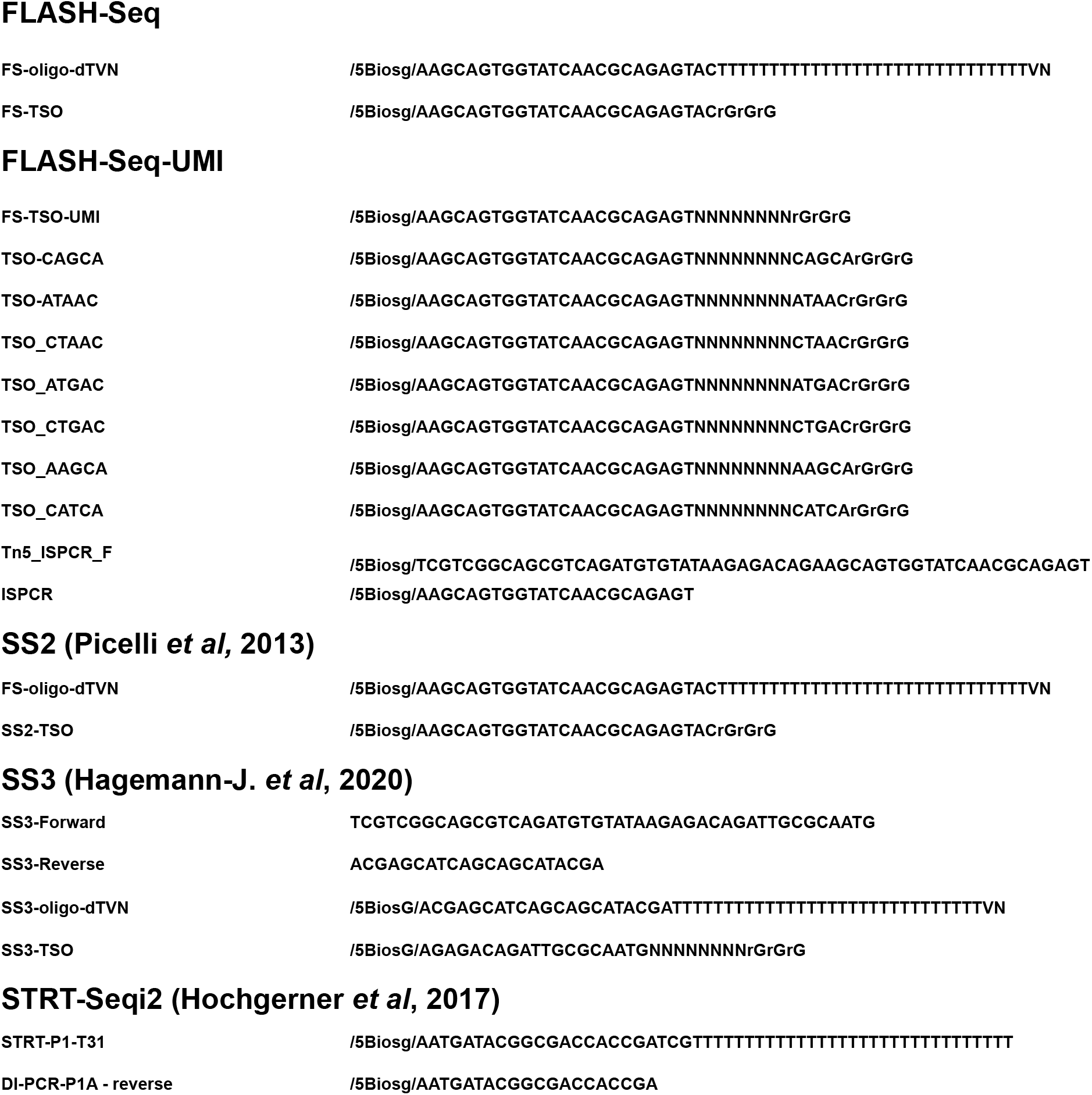

---

**Fig. S1.**
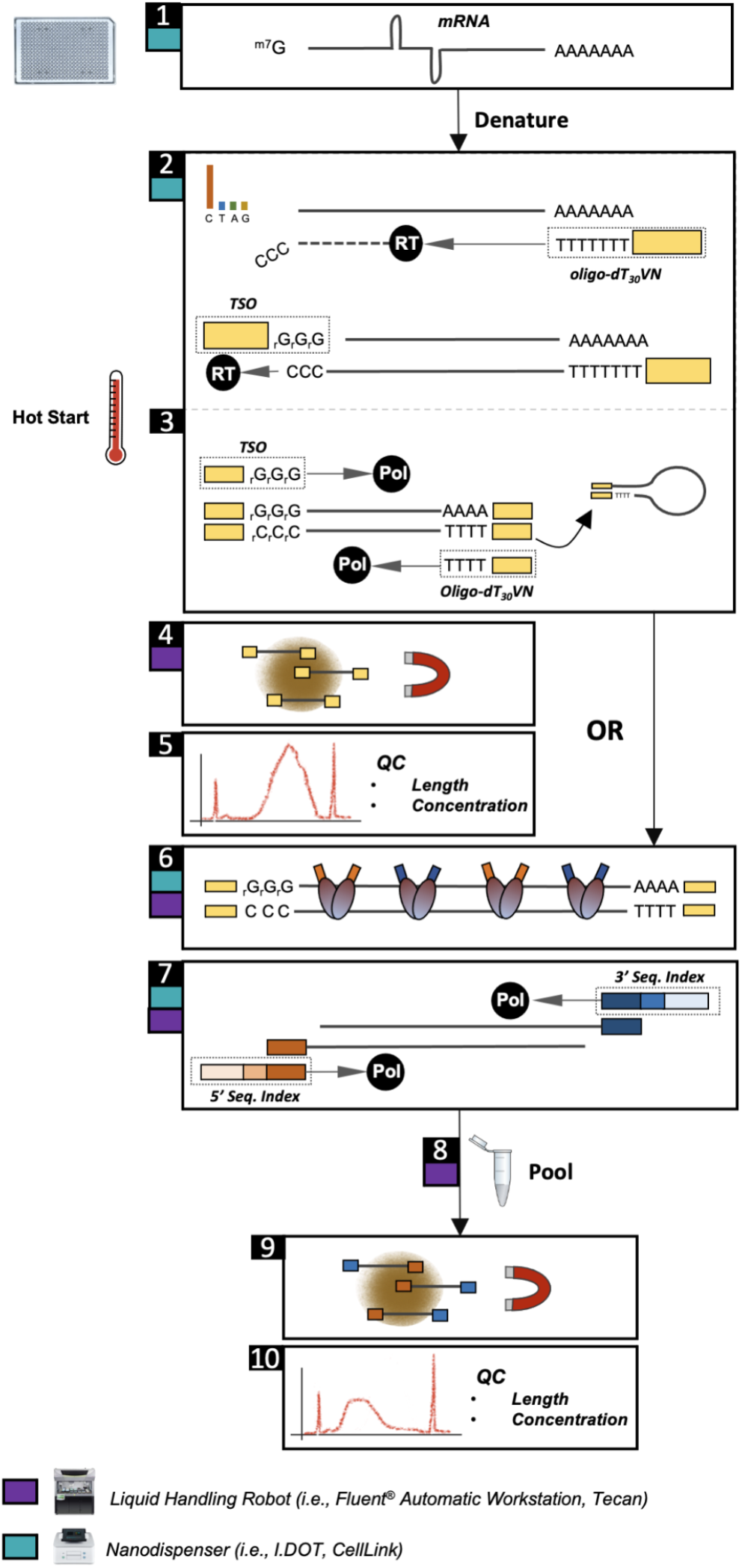
10 Steps of FLASH-Seq Workflow. Steps automated with a liquid handling robot or nanodispenser are marked by a purple and teal square, respectively. **Step 1**, cells are individually sorted in plates in the lysis buffer. **Step 2**, after denaturation, mRNAs are reverse transcribed using a template-switching reverse transcriptase (RT) in the presence of an excess of dCTP. **Step 3**, cDNA is amplified by semi-suppressive PCR. **Step 4,** cDNA is purified using magnetic beads. **Step 5**, cDNA concentration and fragment size are measured and the samples are diluted to a final concentration of ~100-200 pg/μL. **Step 6**, the cDNA is tagmented using a hyperactive Tn5 transposase which introduces known adaptor sequences. **Step 7**, the tagmented cDNA is amplified by PCR and sequencing indices are added. **Step 8**, all samples are pooled together. **Step 9,** the library is purified using magnetic beads. **Step 10**, concentration and average fragment size is measured in preparation for the sequencing.

**Fig. S2.**
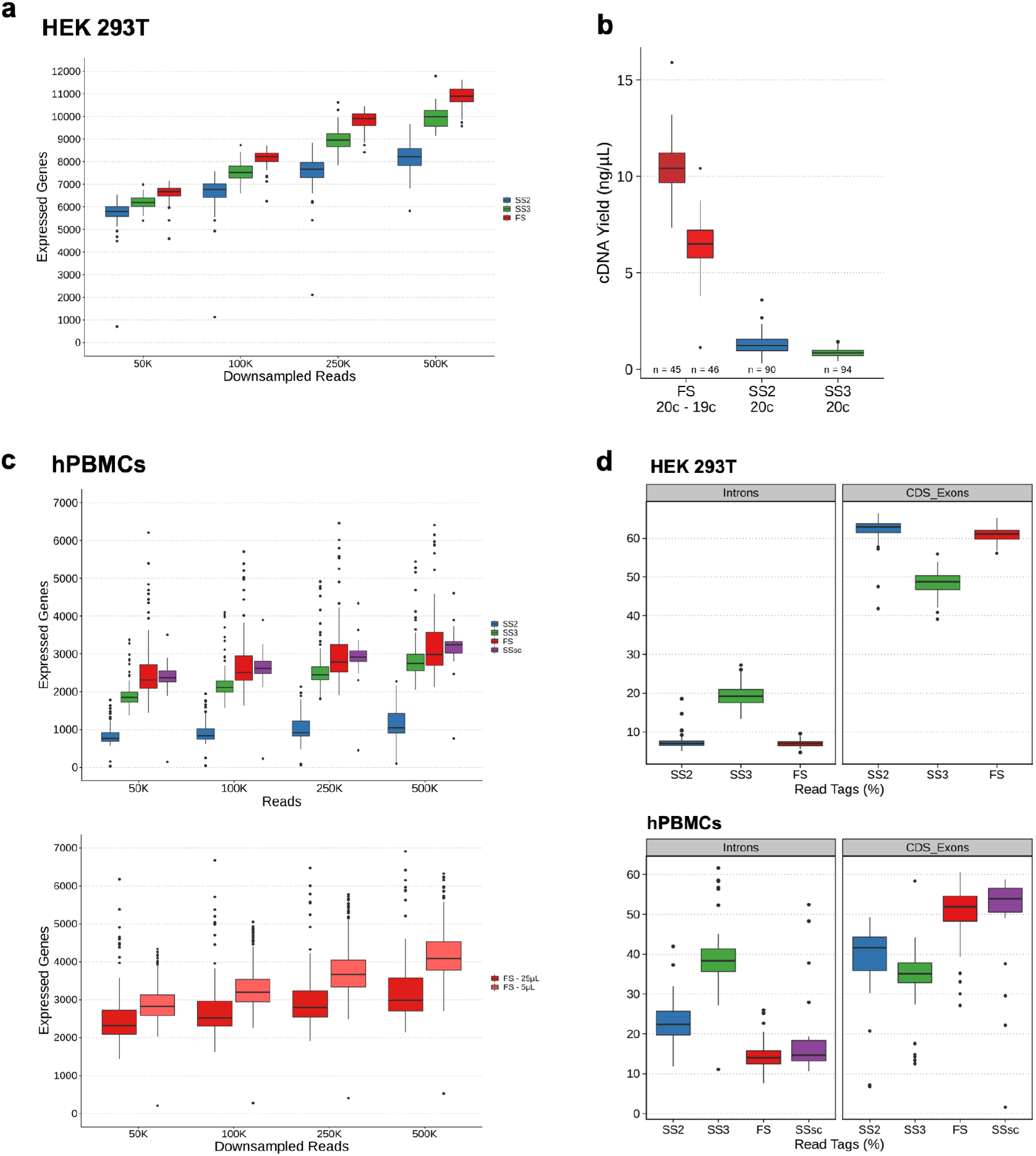
FLASH-Seq gene detection using downsampled reads and cDNA yield. **a.** Number of detected genes in HEK 293T cells processed with SS2 (n = 80), SS3 (n = 42) or FS (n = 85) using 50,000, 100,000, 250,000 or 500,000 downsampled reads. Gene detection threshold was set to >0 read. **b.** cDNA Yield, in ng/μL for SS2 (20 PCR cycles), SS3 (20 PCR cycles) or FS (20 or 19 PCR cycles) **c.** Upper part, number of detected genes in hPBMCs processed with SS2 (n = 47), SS3 (n = 76), FS (n = 136) or SMART-Seq Single Cell Kit (SSsc, Takara Bio, n = 24). FS, SS2, SS3 and SSsc reaction volumes were respectively: 25, 25, 10 and 50 μL. Bottom part, number of detected genes in hPBMCs processed with FS (n = 136, 25 μL) and miniaturized FS (n = 231, 5 μL). Raw reads were downsampled to 50,000, 100,000, 250,000 or 500,000. Gene detection threshold was set to >0 read. **d.** Estimated proportion of read mapped to intronic or CDS-exonic features, in HEK 293T cells and hPBMCs, in read tag percentages.

**Fig. S3.**
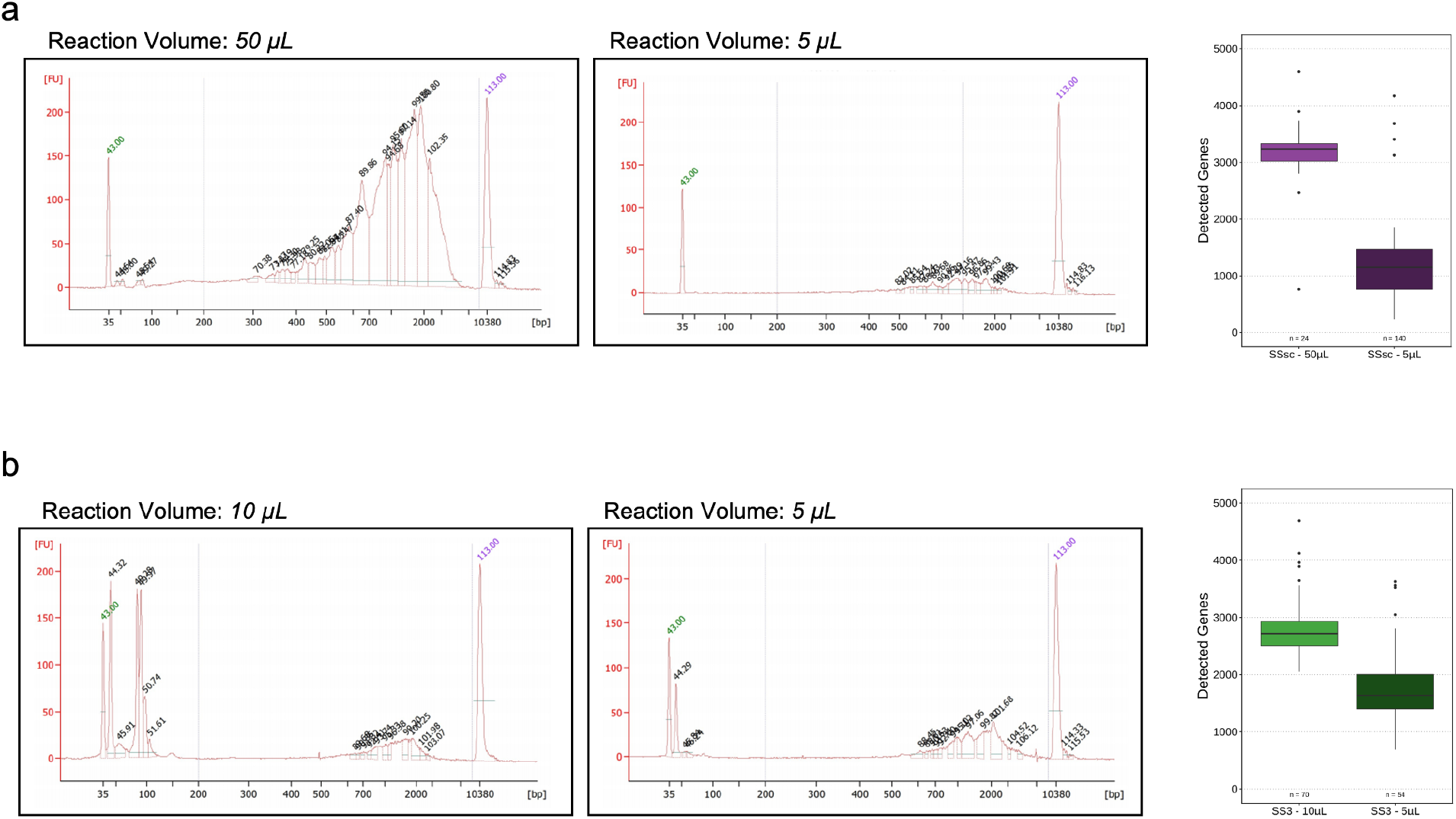
SSsc and SS3 miniaturization to 5 μL. **a.** Bioanalyzer traces illustrating the cDNA length distribution of regular (50 μL) or miniaturized (5 μL) SSsc. The right-panel shows the number of detected genes (>0 read) in both conditions. **b.** Bioanalyzer traces showing the cDNA length distribution in regular (10 μL) or miniaturized (5 μL) SS3. The right-panel shows the number of detected genes (>0 read) in both conditions using 500,000 downsampled raw reads.

**Fig. S4.**
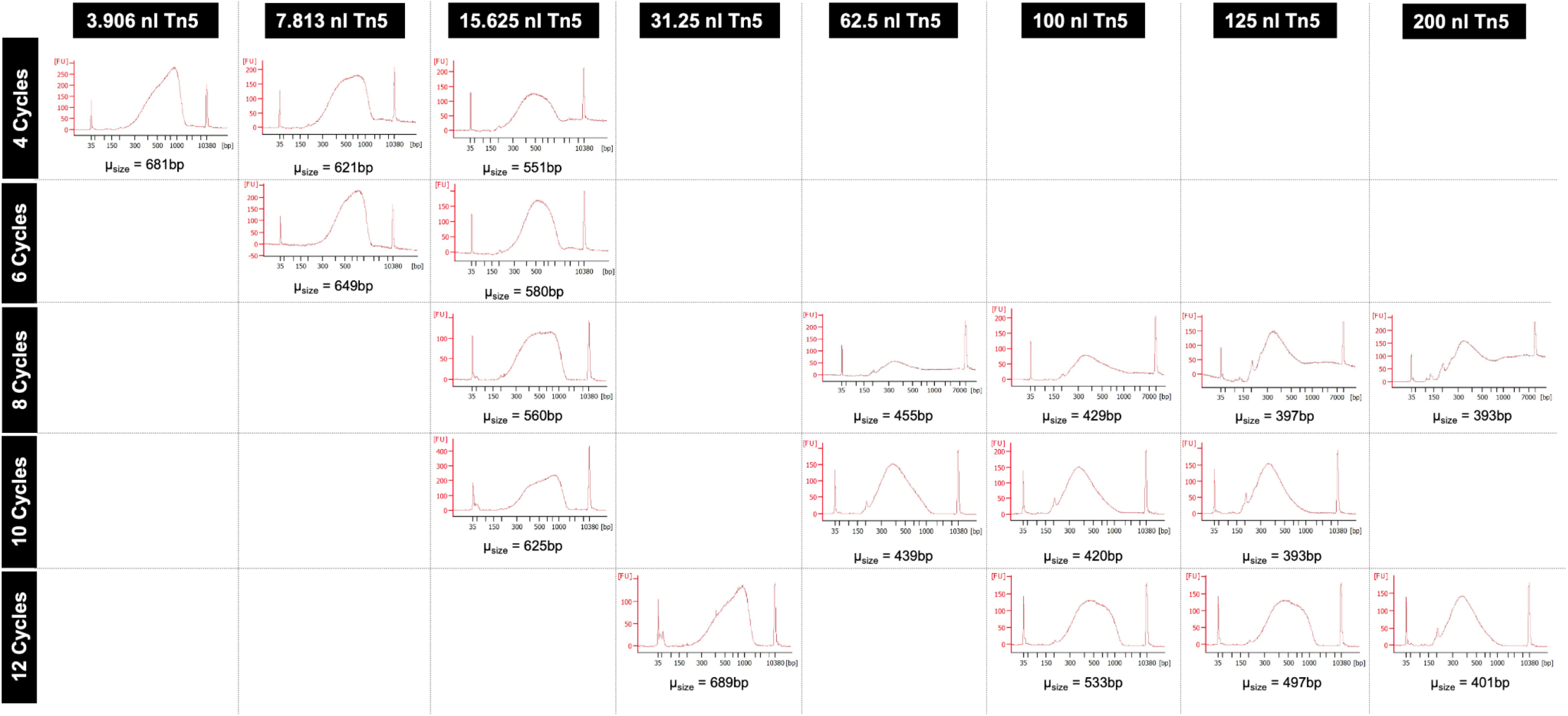
Impact of the number of PCR cycles and Tn5 amount on FS-LA cDNA size distribution. Bioanalyzer traces of the final FS-LA libraries. The number of PCR cycles used for cDNA pre-amplification is depicted on the rows (4 to 12 cycles). The amount of Tn5 added to each cell for tagmentation is depicted on the top of the chart. The mean cDNA size distribution between 200 bp and 9000 bp is displayed below each trace.

**Fig. S5.**
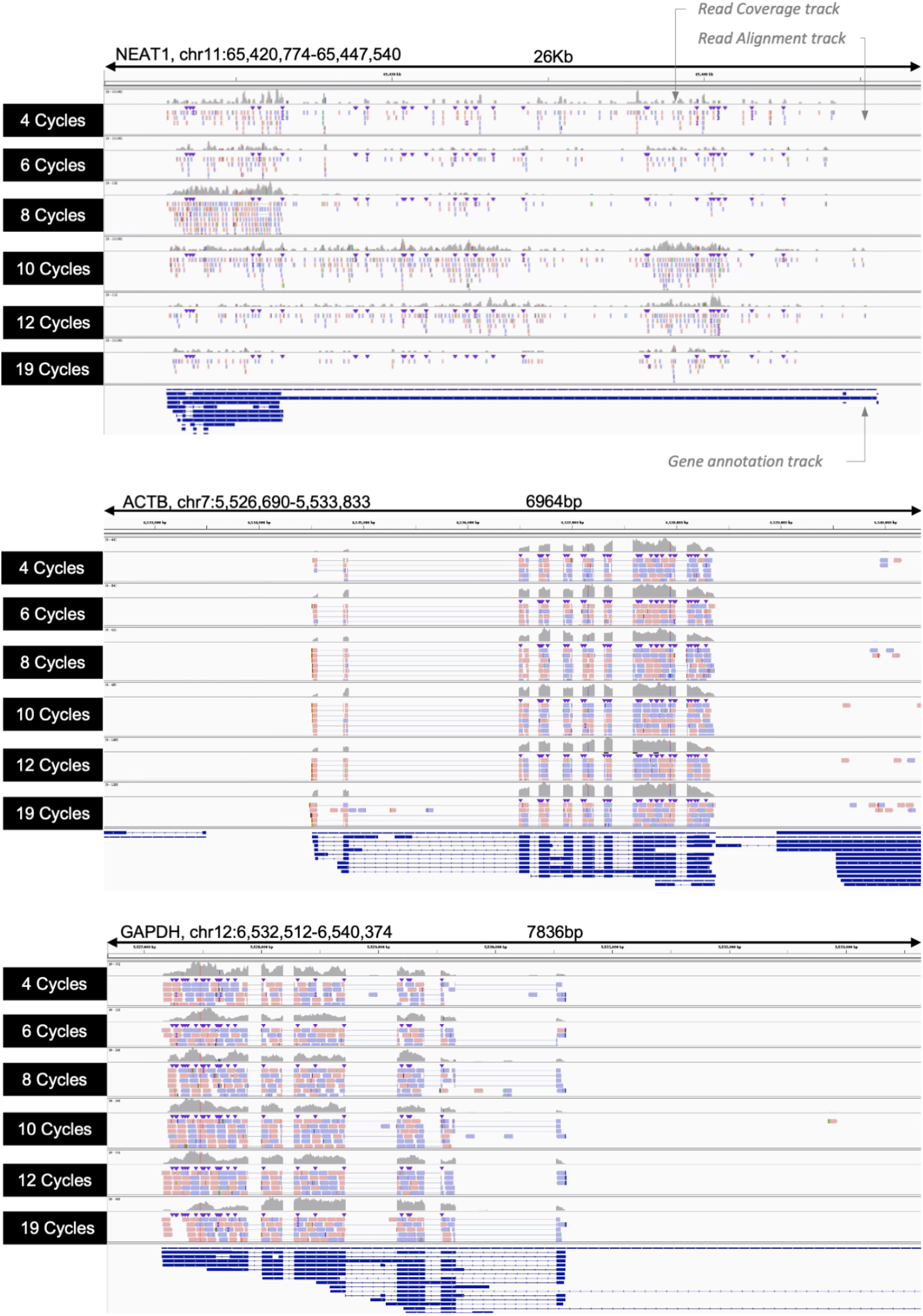
Integrated Genome Viewer visualization of selected genes. Each panel contains six representative samples which were downsampled to 500,000 raw reads and mapped onto hg38. As depicted in the first panel, each sample consists of a gene coverage, a read mapping and gene annotation track. In the read mapping track, each blue or red bar corresponds to a single mapped read. The color of the bar depicts the read orientation compared to the reference. Fine lines highlight split reads. In the gene annotation track, fine blue lines correspond to introns while bold blue lines represent exons. Each panel shows a different gene (from top to bottom: lncRNA-NEAT1, ACTB, GAPDH).

**Fig. S6.**
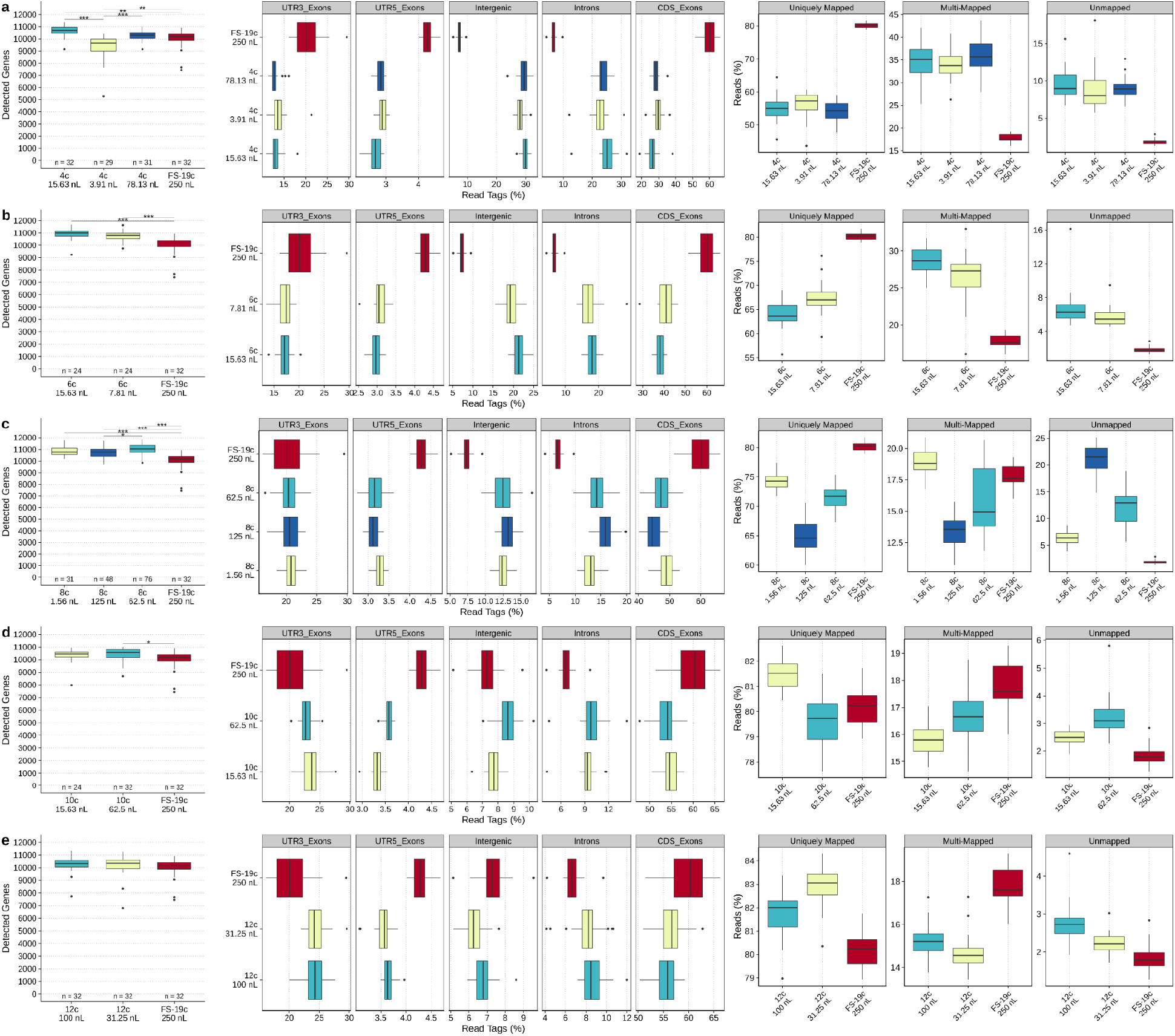
Overview of FS-LA gene detection and mapping statistics. Each panel from right to left: First, the number of detected genes in HEK 293T cells using FS (19 PCR cycles) or FS-LA, 250,000 downsampled reads, gene detection threshold >0 read. Significance level was evaluated using Dunn’s test. P-values were corrected for multiple testing using Bonferroni correction (pval < 0.05). Second, the proportion of read mapped to exonic, intronic or intergenic features measured using ReSQC, in read tag percentages. Third, the percentage of uniquely mapped, multi-mapped or unmapped reads. Each panel compares regular FS (19 PCR cycles, in red) with **a.**4 PCR cycles FS-LA **b.**6 PCR cycles FS-LA **c.**8 PCR cycles FS-LA **d.**10 PCR cycles FS-LA **e.**12 PCR cycles FS-LA.

**Fig. S7.**
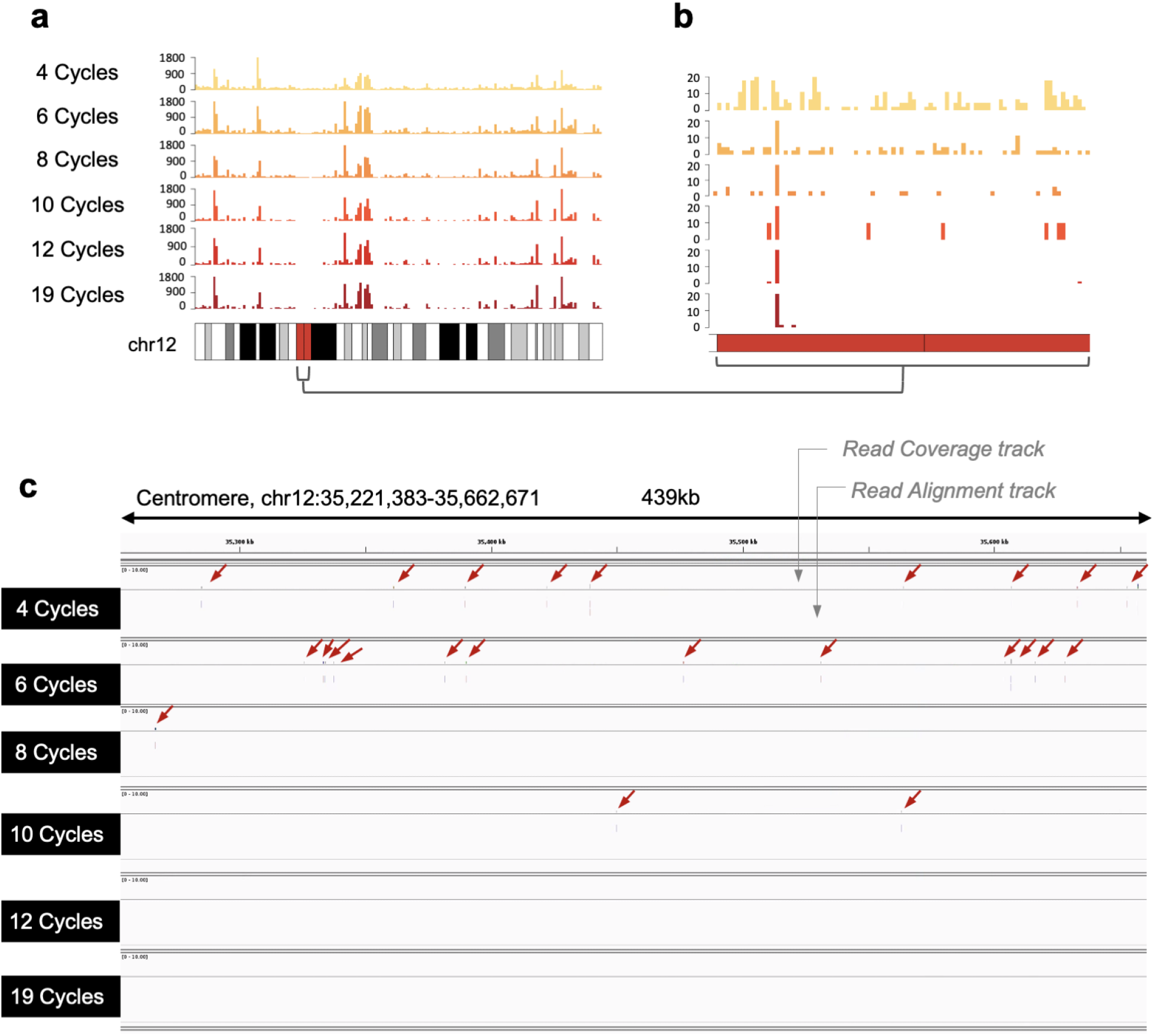
Intergenic reads generated during low-amplification. **a.** Density of mapped reads on chromosome 12, averaged to 750,000 bp windows for 6 representative samples, amplified with 4, 6, 8, 10 or 12 PCR cycles (FS-LA) or 19 PCR cycles (FS) and downsampled to 500,000 raw reads. **b.** Zoom on the centromeric region. **c.** Integrated genome viewer screenshot showing an example of unexpected read mapping on a 439Kb region of chromosome 12 centromere. Mapped reads are marked by a red arrow. Each line represents a different sample processed with 4, 6, 8, 10, 12 or 19 PCR cycles.

**Fig. S8.**
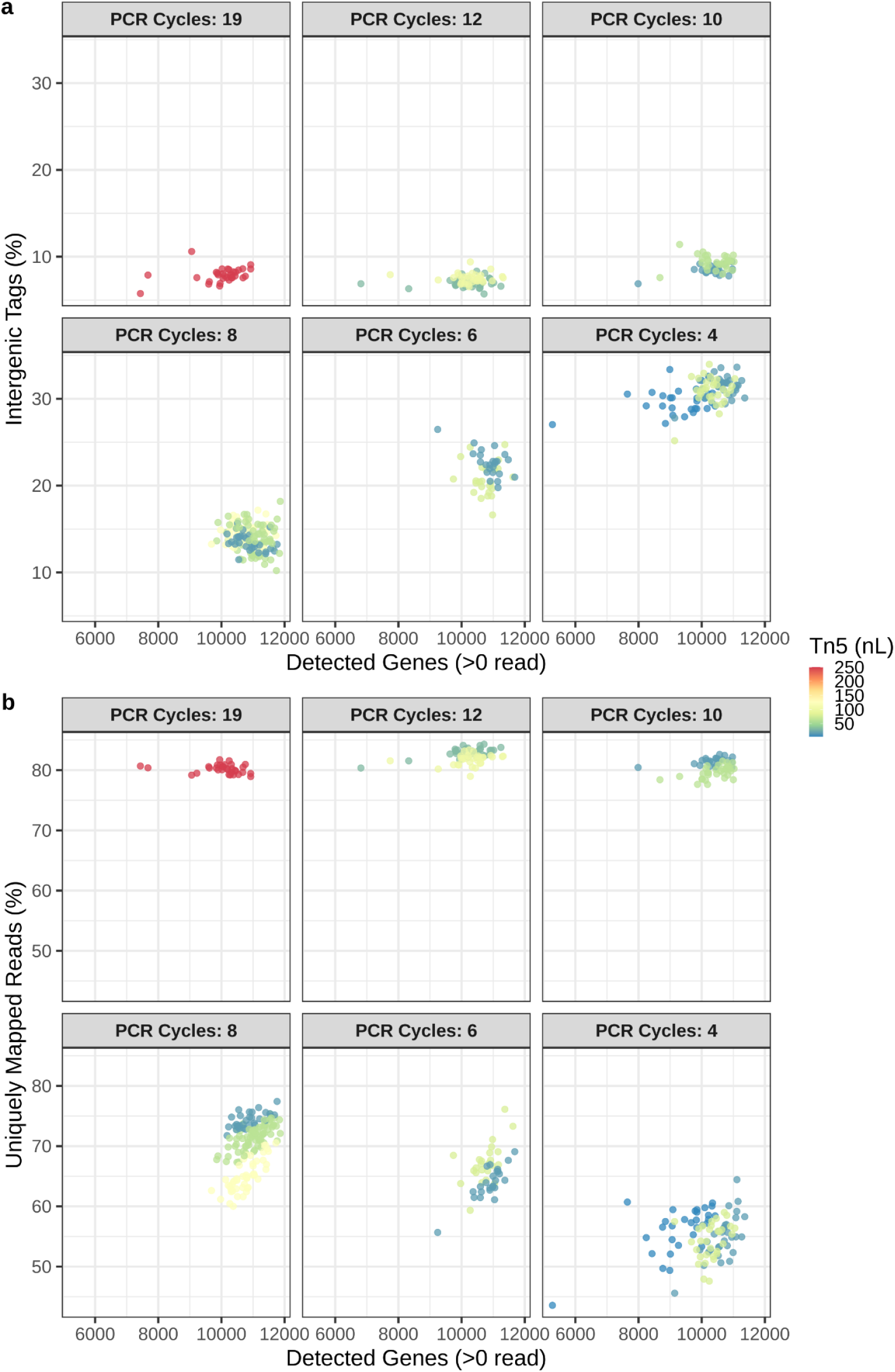
Relation between the number of detected genes. **a.** with the percentage of intergenic reads **b.** the percentage of uniquely mapped reads. Color scale represents the amount of home-made Tn5 used for tagmentation, in nL.

**Fig. S9.**
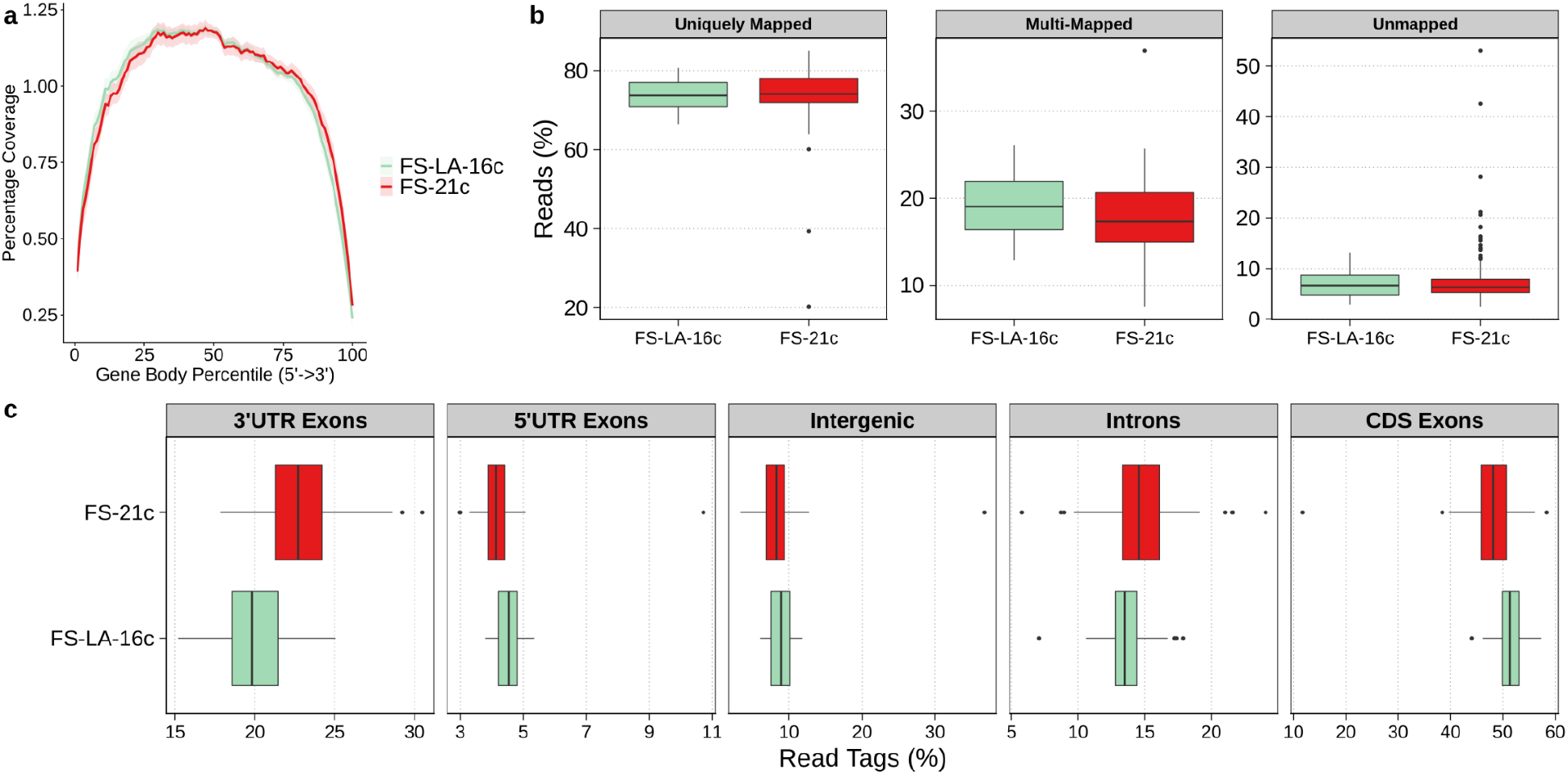
Overview of low amplification hPBMCs mapping statistics. **a.** FS gene body coverage of hPBMCs processed with FS (21 cycles, n = 180) or FS-LA (16 cycles, n = 107). **b.** STAR mapping statistics showing the percentage of uniquely mapped, multi-mapped and unmapped reads. **c.** Distribution of mapped reads between introns, intergenic regions or 3’-UTR / 5’-UTR / Coding sequence (=CDS) exons. Displayed in percentage of read tags and computed using ReSQC.

**Fig. S10.**
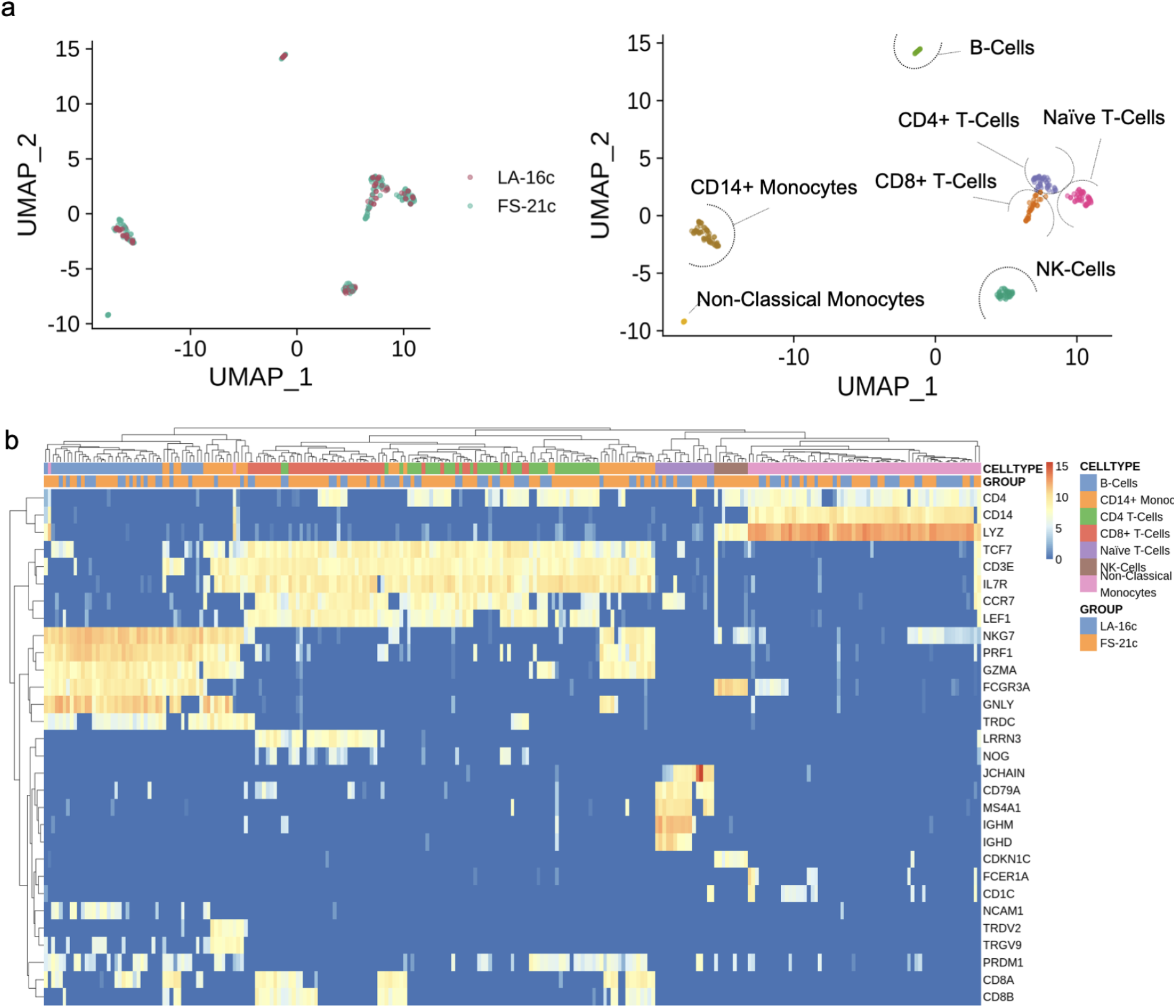
Dimension reduction and cell type assignment of hPBMCs processed with FS or FS-LA. **a.** Dimension reduction of hPBMCs processed with FLASH-Seq (FS-21c, n = 160) or FLASH-Seq low amplification (LA-16c, n = 95) protocol, highlighted by the method (left) or assigned cell types (right). Only cells with >100,000 raw reads and <25% unmapped reads were selected for this analysis. **b.** Heatmap of selected marker genes expressed by cells processed with either FS-21c or FS-LA-16c.

**Fig. S11.**
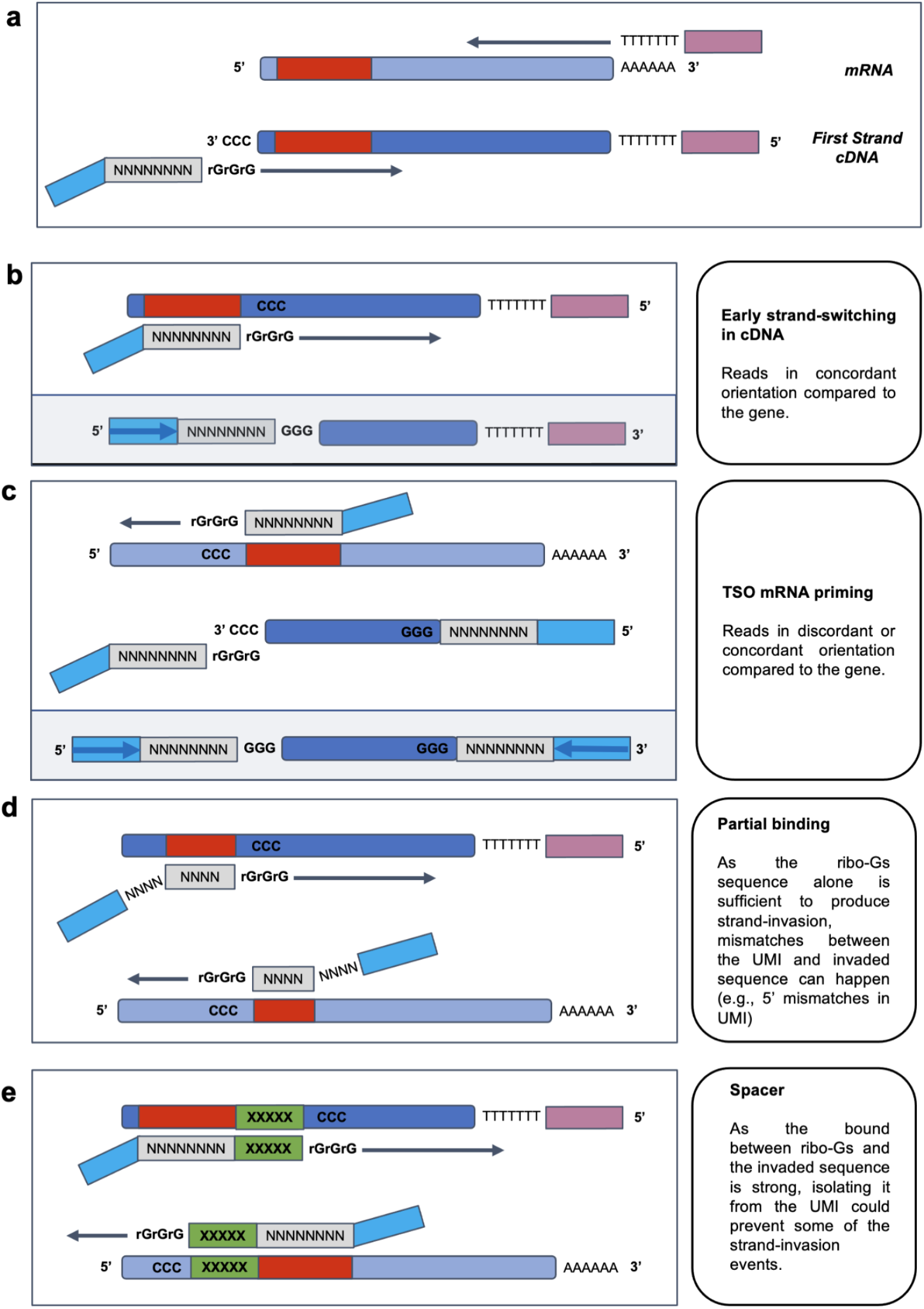
Strand-Invasion mechanisms. **a.** Regular RT and first-strand cDNA synthesis in SS3 protocol. **b.** cDNA Strand-invasion. The UMI and rGrGrG motif bind to a sequence inside of the cDNA **c.** TSO mRNA priming. The TSO replaces oligo-dT to prime the RT. **d.** Example of 5’ partial match between the UMI and the cDNA or mRNA. **e.** Example of TSO invasion when a spacer sequence is used to separate the rGrGrG motif from the UMI.

**Fig. S12.**
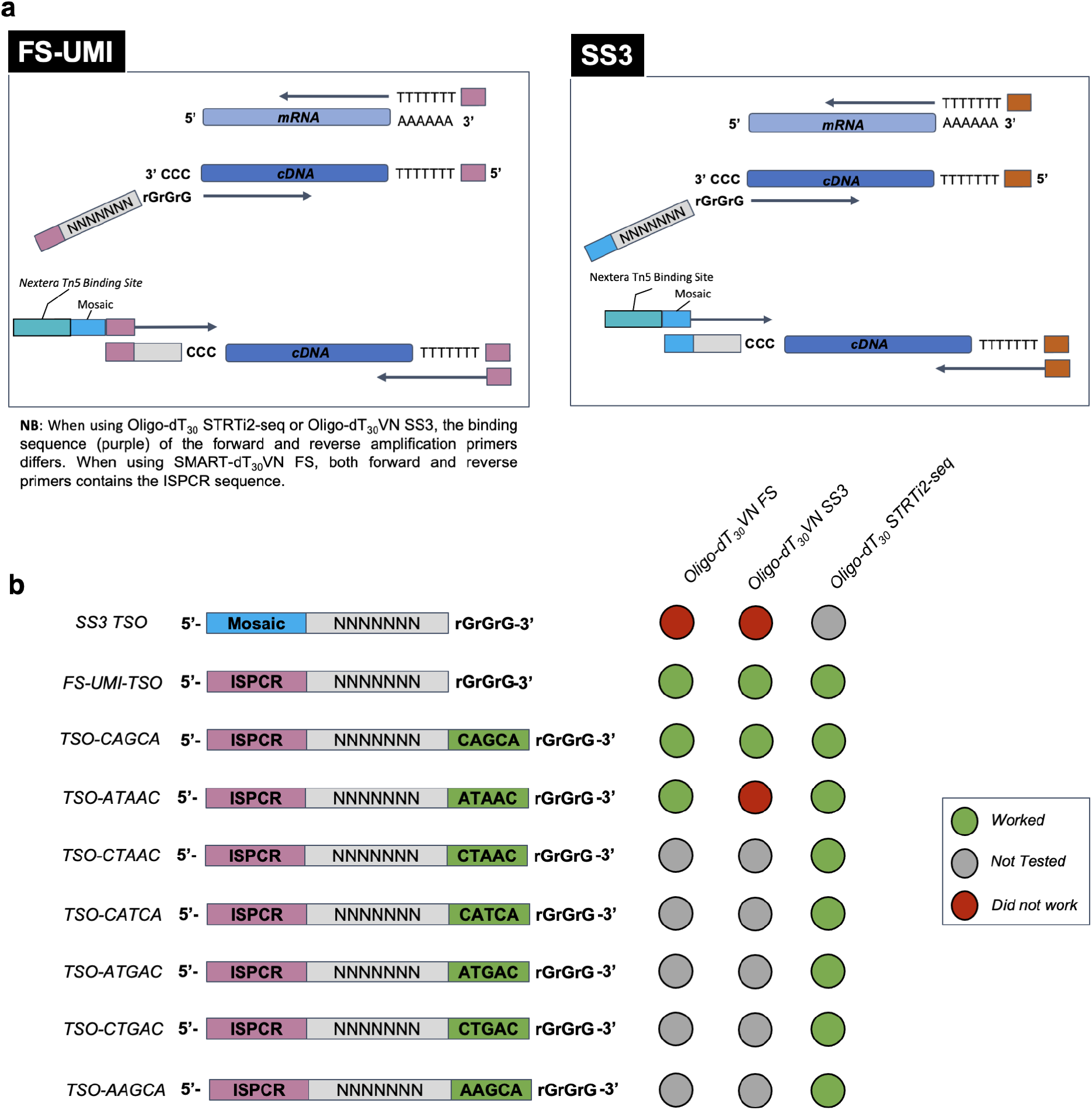
Schematic representation of the tested oligo-dT / TSO combinations. **a.** Reverse transcription and template-switching for the FS-UMI and SS3 protocols. **b.** Combinations of TSO and oligo-dT tested in this study.

**Fig. S13.**
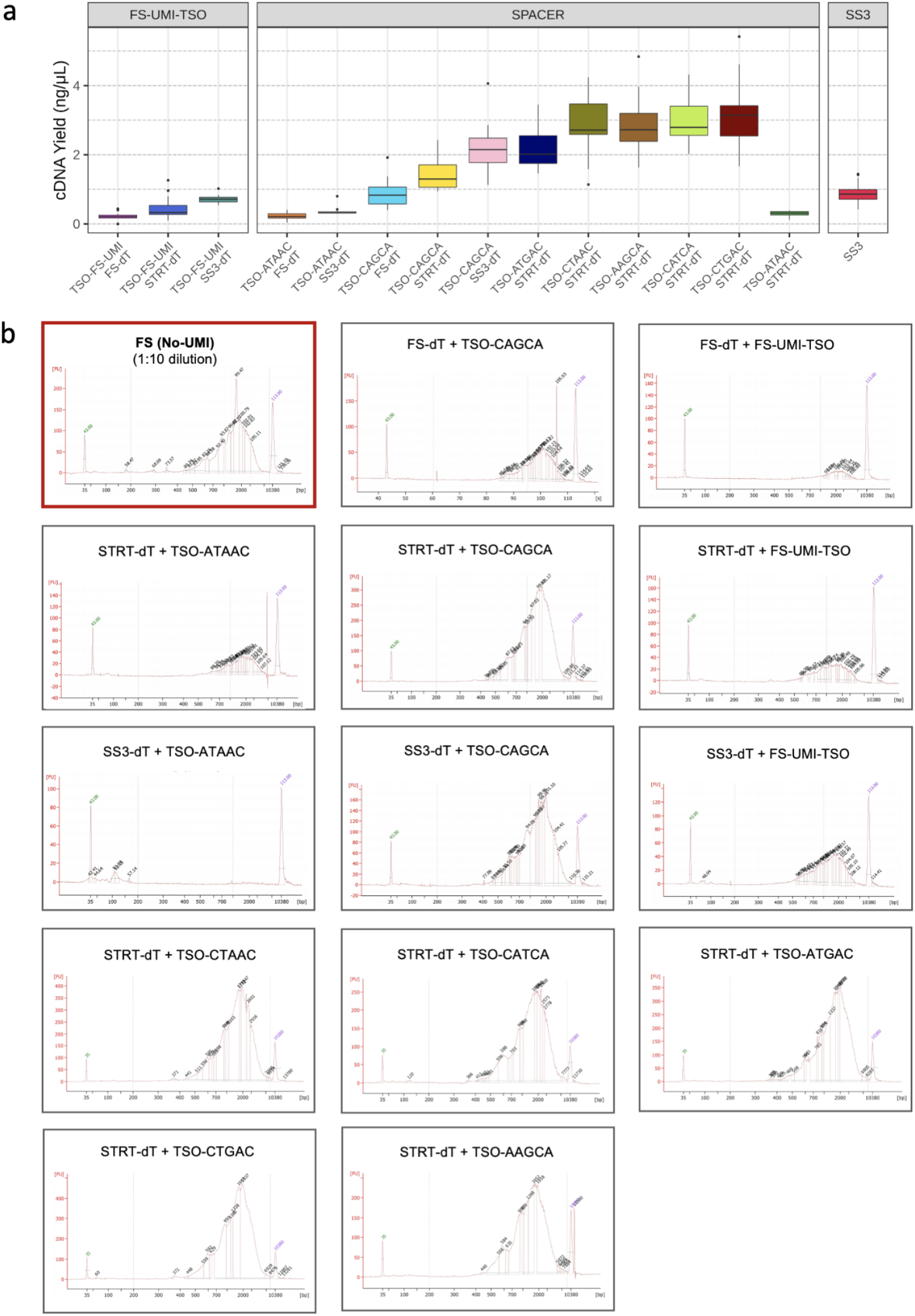
cDNA yield and Length distribution of the UMI-TSO / Oligo-dT combinations. **a.** cDNA yield **b.** Bioanalyzer traces of a representative sample of each UMI-TSO / oligo-dT combination. Control FS performed using regular TSO (= devoid of UMI) is highlighted in red.

**Fig. S14.**
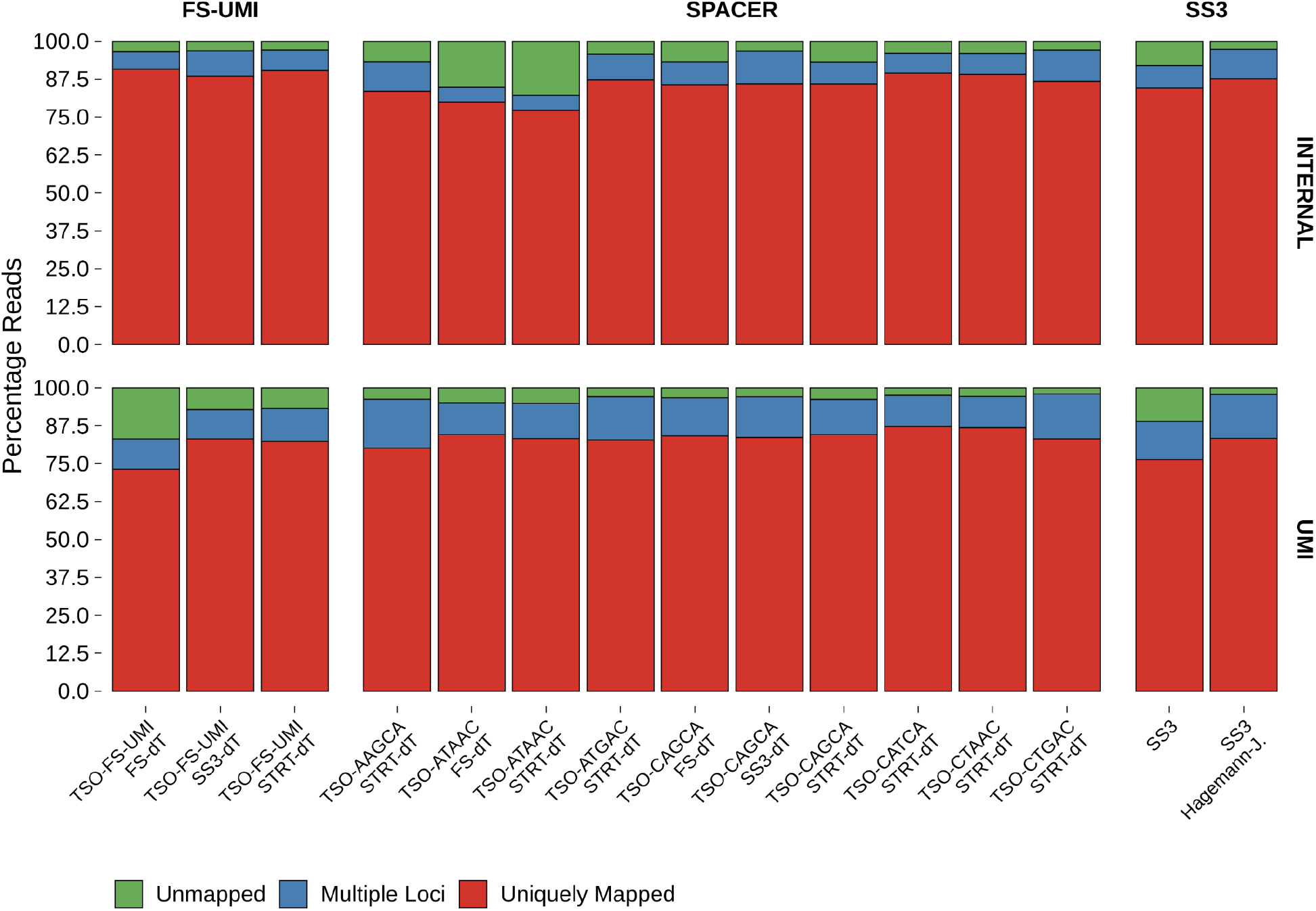
Internal and UMI reads mapping statistics of the different UMI-TSO / oligo-dT combinations. STAR mapping statistics showing the percentage of uniquely mapped, multi-mapped and unmapped reads using either internal or UMI-reads.

**Fig. S15.**
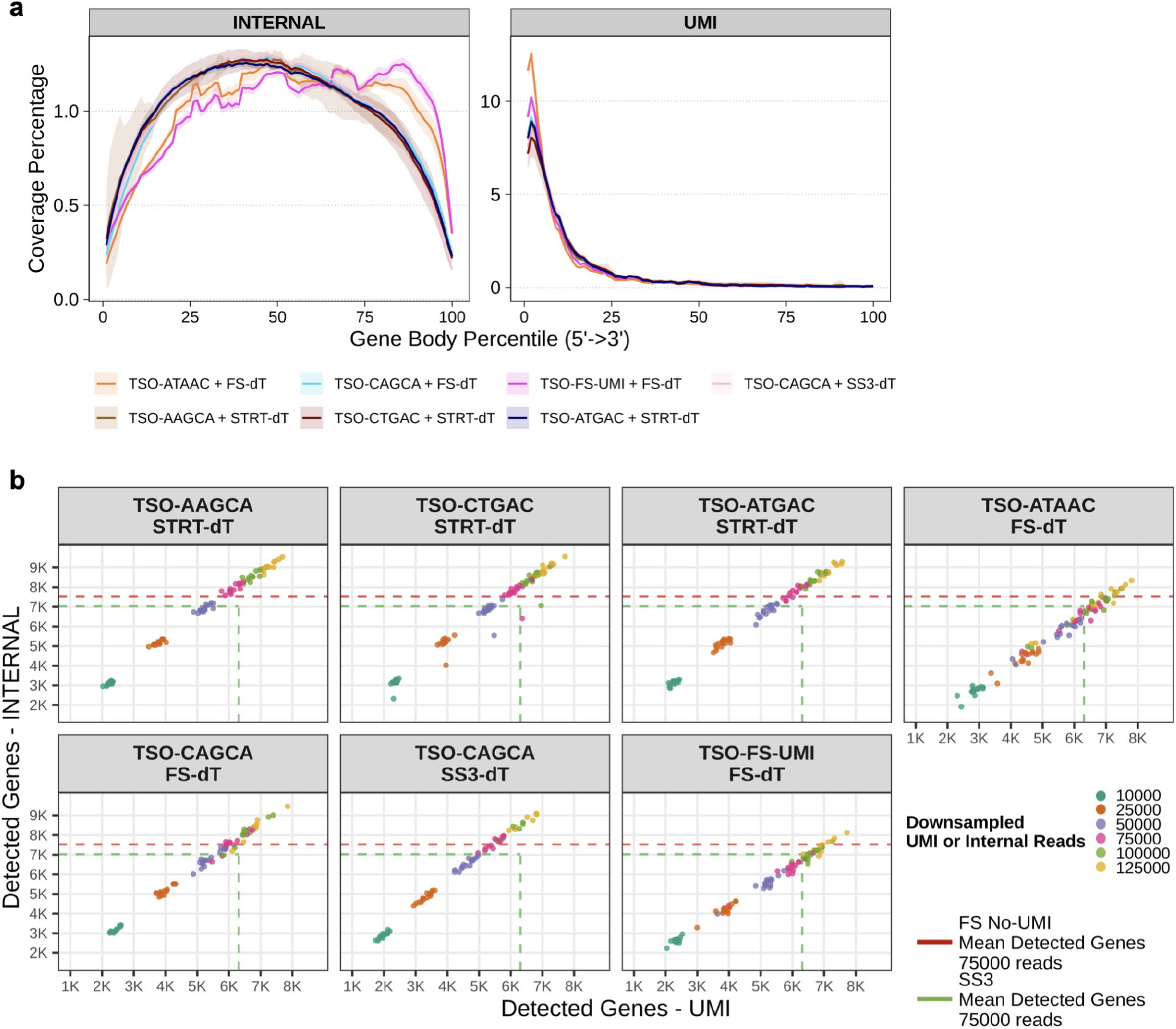
Addition of UMI to FS protocol and evaluation of oligo-dT - TSO combinations (remaining conditions of Fig 3). **a.** Gene body coverages. **b.** Relationship between the number of detected genes using UMIs and internal reads. UMIs and internal reads were downsampled to 10K, 25K, 50K, 75K, 100K, 125K reads. Dashed red line represents the mean number of detected genes using FS (n = 85) downsampled to 75K raw reads. Dashed green lines represent the mean number of detected genes from Hagemann-Jensen *et al* using UMI or internal reads, downsampled to 75K raw reads.

**Fig. S16.**
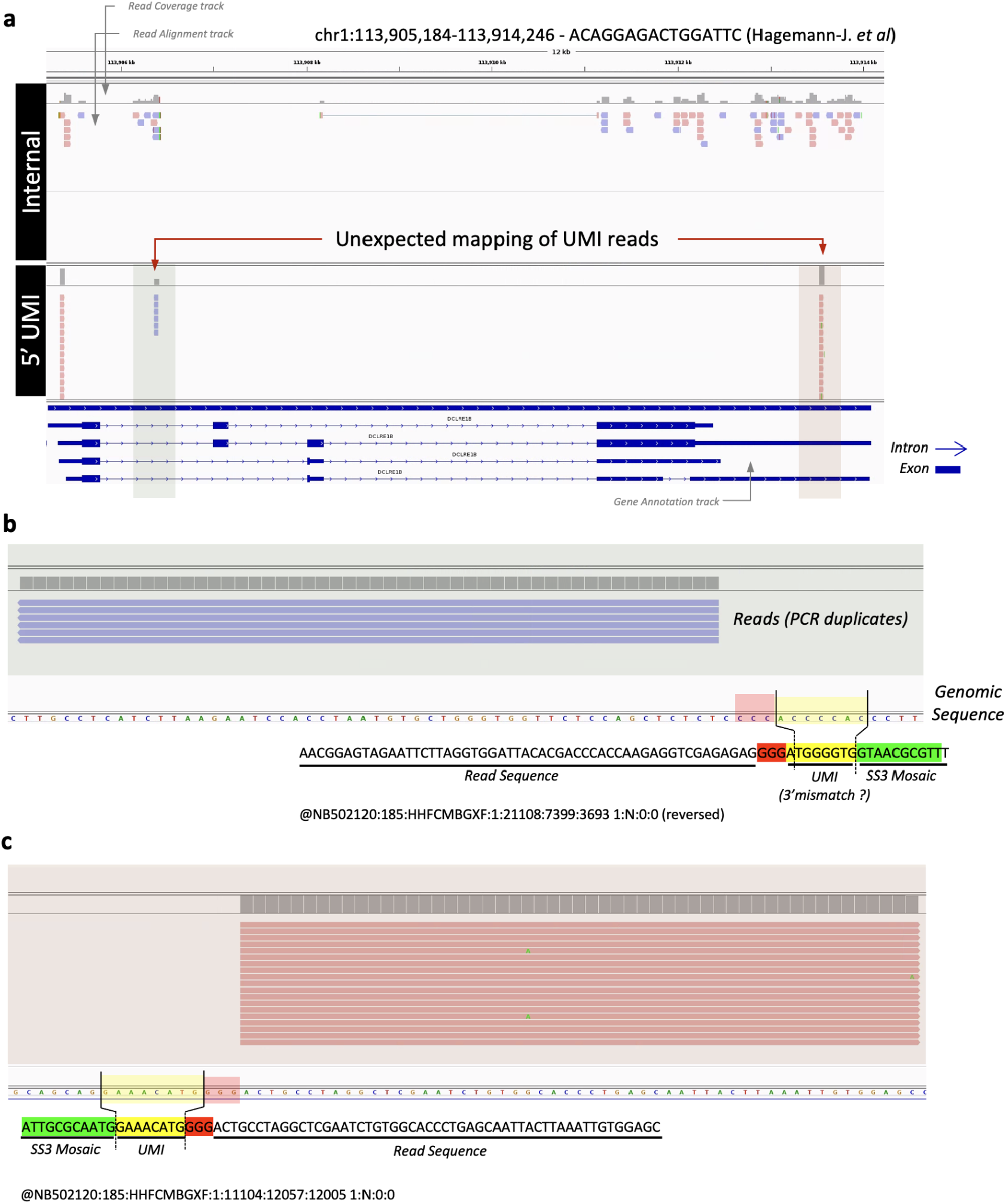
Strand-invasion example. **a.** Integrated Genome Viewer snapshot depicting the reads of the HEK 293T cell ‘ACAGGAGACTGGATTC’ (Hagemann-J. *et al*) mapping to gene DCLRE1B. The upper part of the picture shows the internal reads (= no UMI). The bottom part of the picture displays the 5’ UMI reads. Two piles of pcr duplicated 5’ UMI reads are located inside the gene body rather than at the 5’ end. **b.** Zoom on the first read pile (green in a.). These reads are located in an intronic sequence and are in discordant orientation compared to the gene. The read sequence is displayed at the bottom. The read’s GGG motif and UMI show almost a perfect complementarity with the genomic sequence. **c.** Same as b. but zooming on the second pile of PCR duplicated reads (orange in a.). A perfect match between the GGG motif and UMI the genomic sequence is observed.

**Fig. S17.**
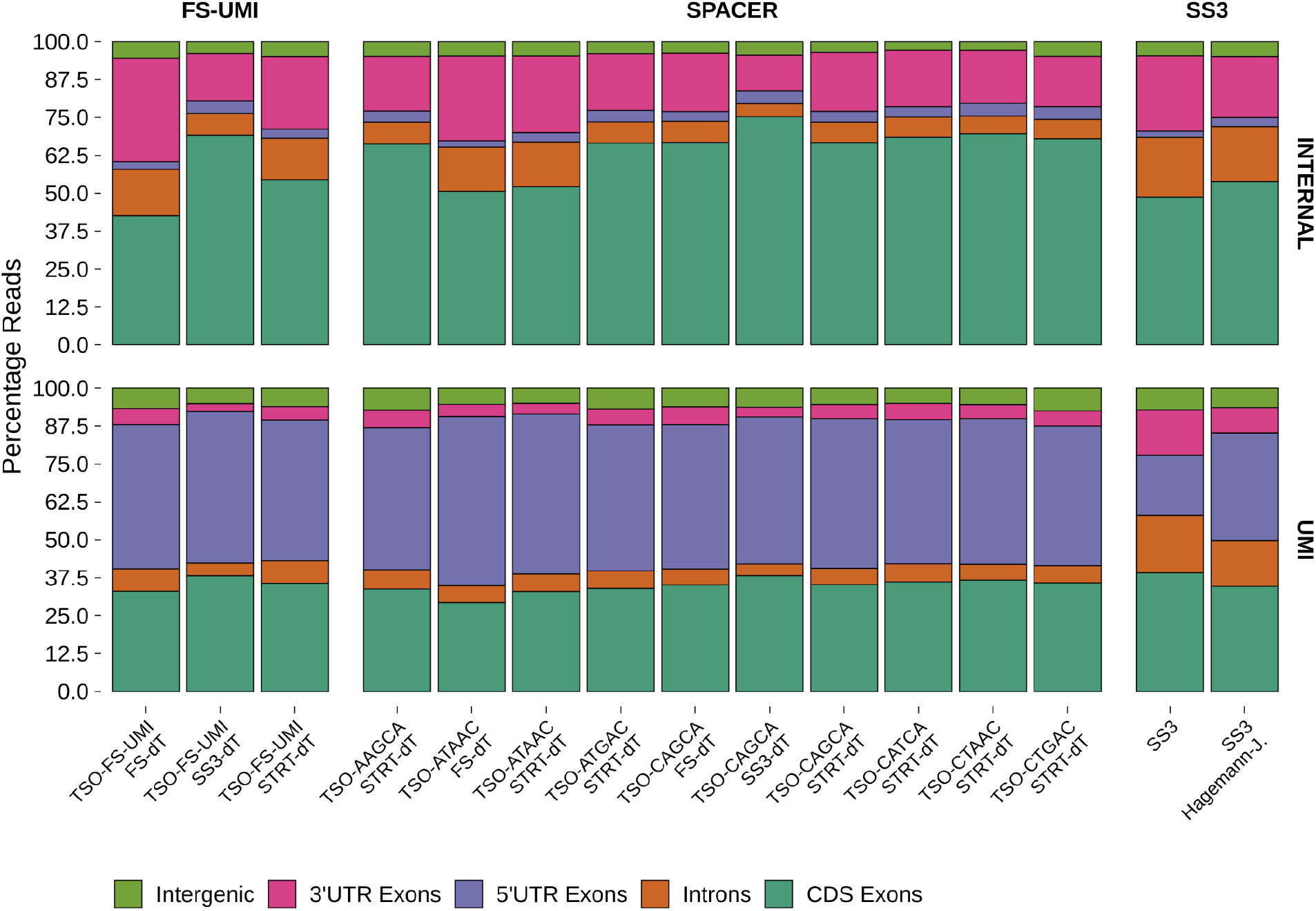
Mapped read distribution in FS-UMI and SS3 protocols. Distribution of mapped internal or UMI-reads between introns, intergenic regions or 3’-UTR / 5’-UTR / Coding sequence (=CDS) exons. Expressed in percentage of read tags and computed using ReSQC.

**Fig. S18.**
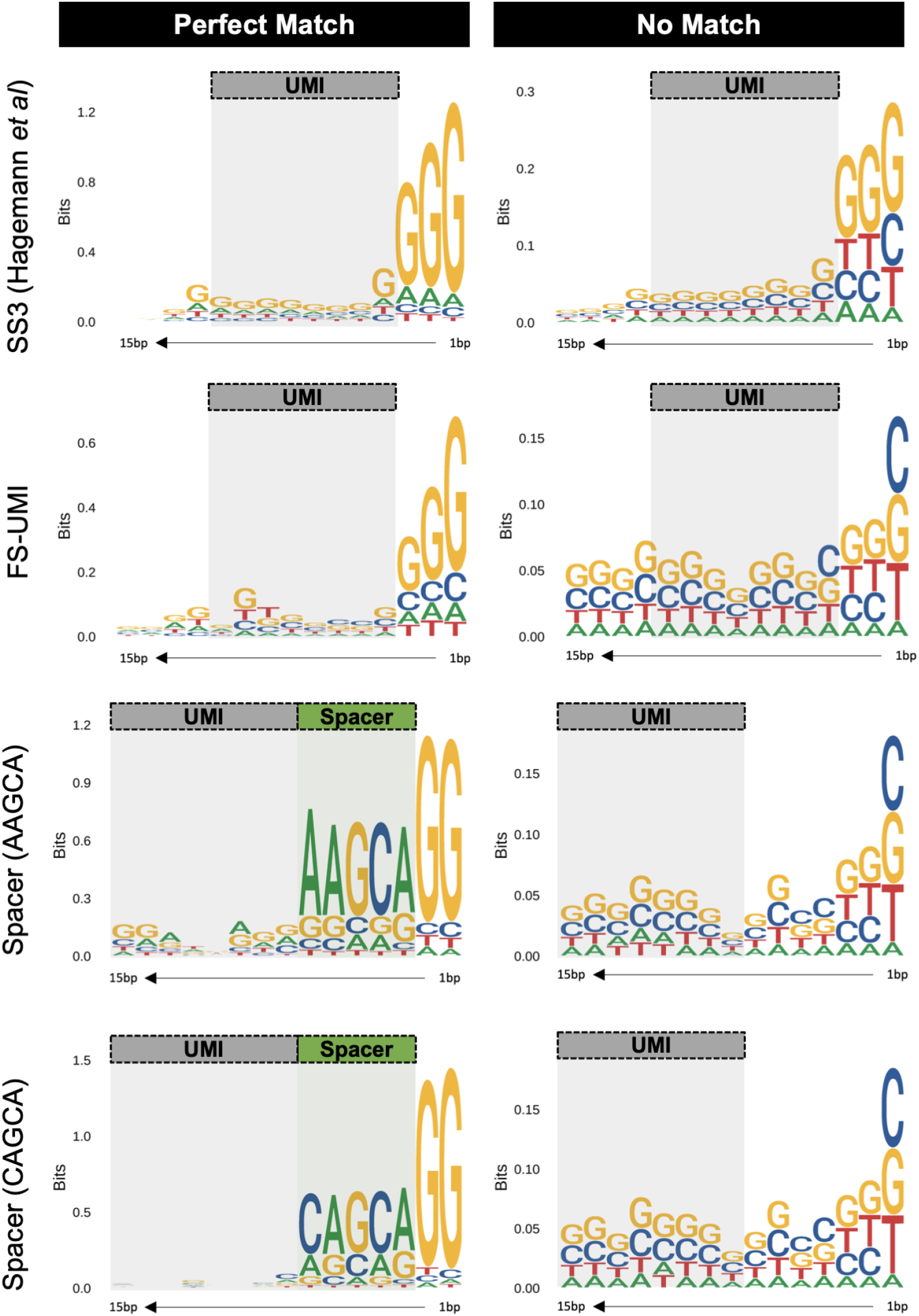
Nucleotide distribution of the genomic sequence adjacent to the read start (in bits). The sequences were split between those displaying a perfect match between the UMI and adjacent sequence (left) and those which did not (right). The expected position of the UMI is highlighted in gray. The expected position of the spacer is highlighted in green. Each row corresponds to one of four representative TSO + oligo-dT conditions.

**Fig S19.**
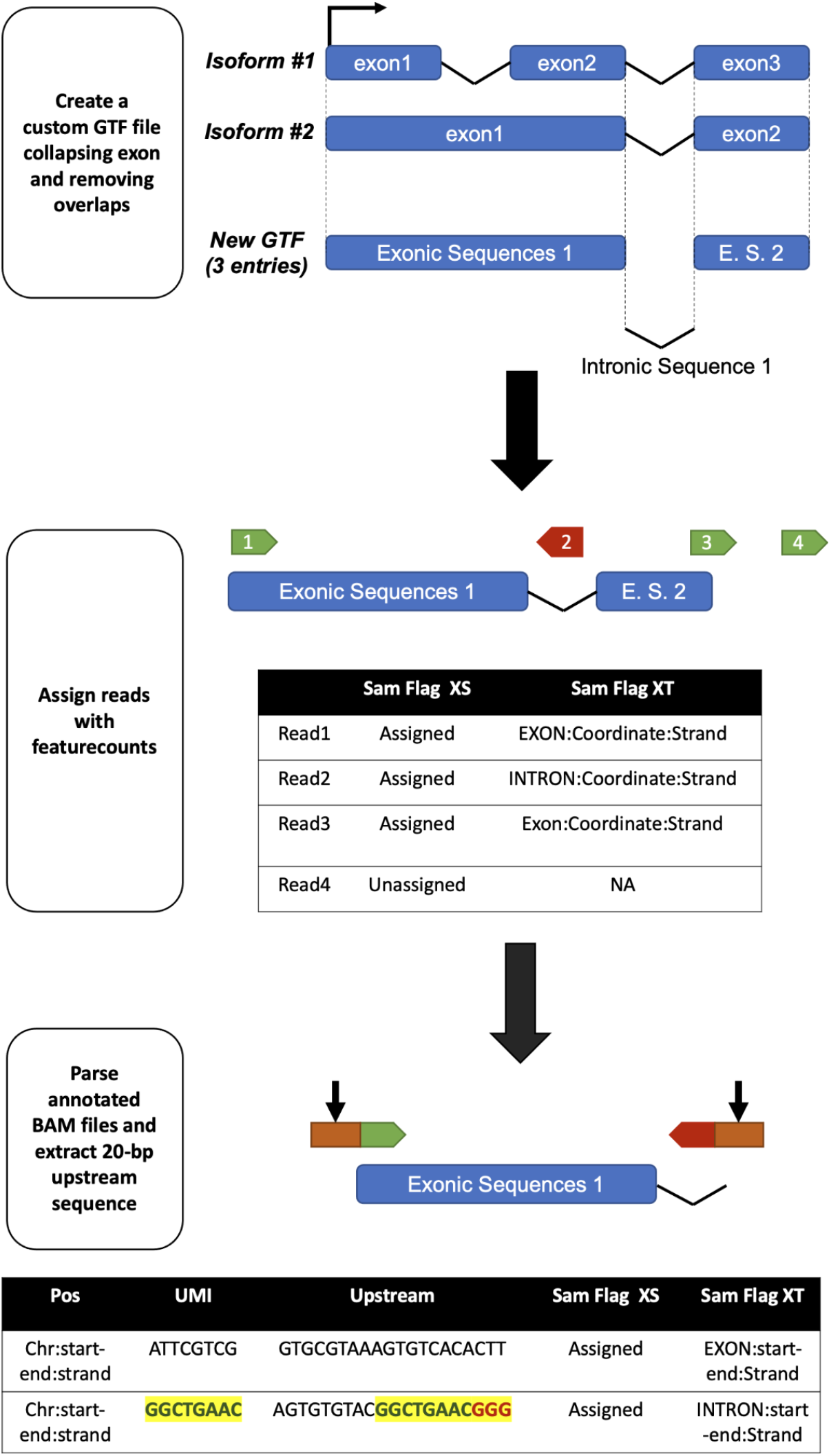
Schematic representation of the strand invasion detection script. First a custom GTF containing collapsed exon-intronic sequences of hg38 protein coding genes is created. Mapped 5’ UMI reads are assigned to a feature using featurecounts. The SAM Flag XT contains the feature coordinates, strand and if the read is mapped unambiguously to an exonic or an intronic sequence. The BAM file is parsed using a custom R script and the 20-bp sequence adjacent to the read start is extracted. This sequence is compared with the UMI sequence, looking for a perfect match.

