## Extended File #2 for "Lightning Fast and Highly Sensitive Full-Length Single-cell sequencing using FLASH-Seq"

### Guidelines - FS Low Amplification

When processing cells using FS “Low-Amplification” (FS-LA) protocol, three important parameters must be taken into account:

- The cell RNA content, which will determine how many pre-amplification cycles are required to generate enough cDNA for the tagmentation while minimizing unmapped / intergenic reads. We currently hypothesise that the excess of intergenic/unmapped reads may originate from a mixture of tagmentation of the genomic DNA (gDNA) and overtagmentation of the messenger RNA (mRNA).
- The reaction volume. As the cDNA is not purified prior to tagmentation, it is important to strike the right balance between diluting enough the salts and additives of the RT-PCR mix and not overdoing it, in order to avoid an unnecessary waste of reagents, which ultimately results in a higher cost per cell. We recommend diluting the unpurified cDNA 10 times for the best results, although lower dilutions might also work.
- The amount of Tn5 needs to be adjusted case by case.

In the following paragraphs we provide some recommendations regarding the adjustment of these parameters.

**cDNA pre-amplification**

To determine the adequate number of PCR cycles we relied on estimating the ratio between the cell mRNA and gDNA. We first estimate the cell mRNA content based on the amount of total RNA recovered from $1x10^6$ cells. For instance, extracting total RNA from HEK 293T cells generates ~16 µg[1] of RNA while hPBMCs gives ~8 µg[1]. Among hPBMCs, dendritic cells are closer to ~4 µg[1]. We then assume the worst and unlikely scenario in which the entire cell gDNA would be available for tagmentation (= 3.1 pg gDNA). Assuming 5% of mRNA and ~60% PCR efficiency (see Fig. 2b) we can draft the following chart:

| | HEK 293T Cells (~16 pg total RNA) | | hPBMCs (~ 4 pg total RNA) | |
|---|---|---|---|---|
| PCR Cycles | mRNA | Ratio mRNA / gDNA | mRNA | Ratio mRNA / gDNA |
| *Starting Material* | 0.8 | 0.3 | 0.2 | 0.1 |
| 1 | 1.3 | 0.4 | 0.3 | 0.1 |
| 2 | 2.0 | 0.7 | 0.5 | 0.2 |
| 3 | 3.3 | 1.1 | 0.8 | 0.3 |
| 4 | 5.2 | 1.7 | 1.3 | 0.4 |
| 5 | 8.4 | 2.7 | 2.1 | 0.7 |
| 6 | 13.4 | 4.3 | 3.4 | 1.1 |
| 7 | 21.5 | 6.9 | 5.4 | 1.7 |
| 8 | 34.4 | 11.1 | 8.6 | 2.8 |
| 9 | 55.0 | 17.7 | 13.7 | 4.4 |
| 10 | 88.0 | 28.4 | 22.0 | 7.1 |
| 11 | 140.7 | 45.4 | 35.2 | 11.3 |
| 12 | 225.2 | **72.6** | 56.3 | 18.2 |
| 13 | 360.3 | 116.2 | 90.1 | 29.1 |
| 14 | 576.5 | 186.0 | 144.1 | 46.5 |
| 15 | 922.3 | 297.5 | 230.6 | 74.4 |
| 16 | 1475.7 | 476.0 | 368.9 | **119.0** |
| 17 | 2361.2 | 761.7 | 590.3 | 190.4 |
| 18 | 3777.9 | 1218.7 | 944.5 | 304.7 |
| 19 | 6044.6 | 1949.9 | 1511.2 | 487.5 |

As shown in the paper, we titrated the number of PCR cycles required to minimize the percentage of intergenic/unmapped reads in HEK 293T cells. These values stabilized around 10-12 PCR cycles. This would be equivalent to a mRNA / gDNA ratio of ~73-fold.

We then moved on to hPBMCs assuming all cells had an mRNA content comparable to the smallest ones in the population (i.e., Dendritic cells, 4 µg). Applying the same reasoning, we settled for 16 PCR cycles, to reach a ratio similar to the one that worked in HEK 293T cells. We obtained high-quality libraries on the first test.

When processing cell types for which no recommendation exists, we advise to perform a first titration experiment to compare the number of intergenic/unmapped/uniquely mapped reads obtained when processing the cells using regular FS (i.e., 19-23 cycles) and FS-LA (12-16 cycles). We foresee that the adequat number of PCR cycles for most cell types will fall between 12 and 16 PCR cycles.

**Tagmentation**

Similarly to other SMART-protocols, the amount of Tn5 used may have to be adjusted to generate sequencing libraries with an insert size between 300 and 600 bp. When working with minute amounts of cDNA, overtagmentation can become an issue, as illustrated below:

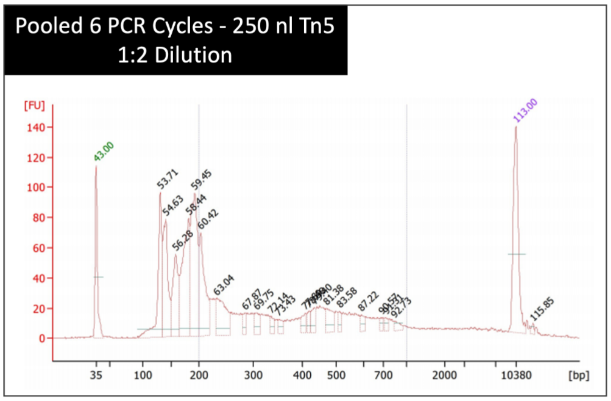


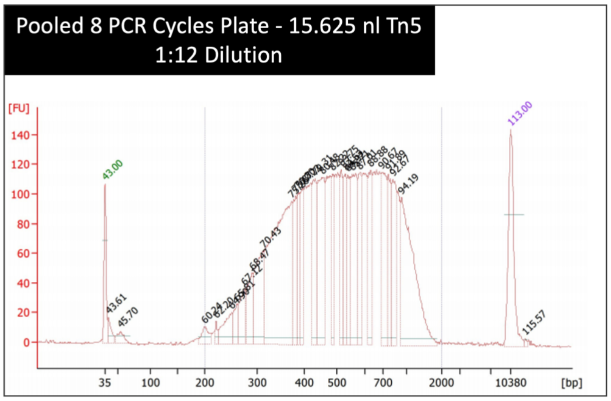


While we typically use 250 nL of our homemade Tn5 to process regular FS (150 pg), values between 8- (31.125 nL) and 16- (15.625 nL) times less Tn5 usually worked well for the low amplification protocol.

**Library Amplification**

We adapted the number of PCR cycles based on the amount of cDNA after the pre-amplification reaction. Libraries from both hPBMCs (16 PCR cycles) and HEK 293T cells (12 PCR cycles) were amplified using 14 PCR cycles. We therefore recommend first testing 14 PCR cycles which should work in most conditions, once the adequate number of PCR cycles is found for the cDNA pre-amplification.

| cDNA pre-amplification (PCR cycles) | Library Amplification |
| --- | --- |
| 4 (HEK 293T) | 24 |
| 6 (HEK 293T) | 22 |
| 8 (HEK 293T) | 18 |
| 10 (HEK 293T) | 16 |
| 12 (HEK 293T) | 14 |
| [redacted] | |
| 16 (hPBMCs) | 14 |

**Reference**


1. https://www.miltenyibiotec.com/_Resources/Persistent/ca9f513c68ed01981bc4d7aa25b01c90db75e6f5/Average_RNA_yields.pdf
