## Extended File #1 for "Lightning Fast and Highly Sensitive Full-Length Single-cell sequencing using FLASH-Seq"

### Extended Information #1

In the following paragraphs we summarize the results of the tests we performed to establish FLASH-Seq (FS) and that we deemed good enough for sequencing. We evaluated the benefits of new additives and reaction conditions, either by using HEK 293T cells or more challenging cells, such as human peripheral blood mononuclear cells (hPBMCs) which in our opinion better reflect a real scientific experiment than cell lines. The cell type used in each experiment (HEK 293T or hPBMCs), and final reaction volume (25 µL or 5 µL) are indicated in square brackets. These tests were aimed at developing a protocol that would be fast to execute and affordable, while providing excellent gene detection even in the most challenging cells (i.e, hPBMCs). Whenever possible we tried to compare gene expression between cells coming from the same plate or same batch to minimize potential batch effects. All comparisons below were performed using 100,000 downsampled raw reads, except when stated otherwise. The gene expression threshold was set to >0 read.

**p-values**: * (<0.05), ** (<0.005), *** (<0.0005)

### Cell Lysis

An ideal lysis buffer would break the cell membrane and free the mRNA content without damaging it or affecting the downstream reactions. We tested the following lysis conditions:

**Triton X-100 1.2% [hPBMCs-5µL], Fig. E1:** Increasing the concentration of Triton X-100 in the lysis buffer resulted in an increased percentage of multi-mapped reads.

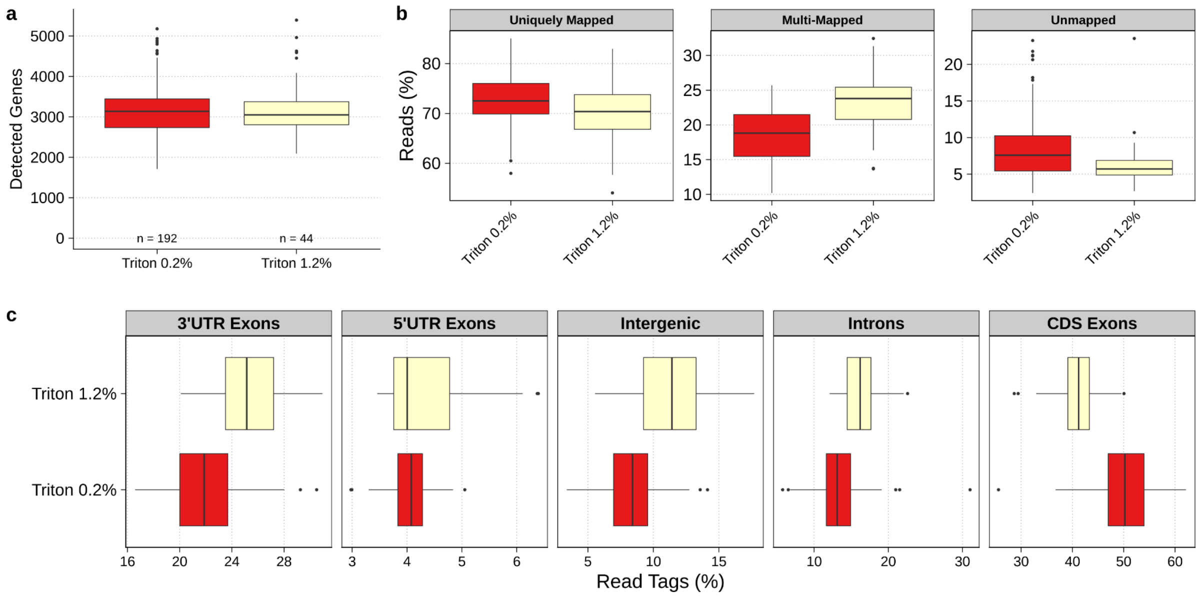


**Fig. E1 | hPBMCs - 5µL - Triton X-100 1.2% a.** Number of detected genes. No significant difference was observed (Mann-Whitney U test). **b.** STAR mapping statistics showing the percentage of uniquely mapped, multi-mapped and unmapped reads. **c.** Distribution of mapped reads between introns, intergenic regions or 3'-UTR / 5'-UTR / Coding sequence (=CDS) exons. Expressed in percentage of read tags and computed using ReSQC.

**Bovine Serum Albumin (BSA, 1 mg/uL) [HEK-25µL/hPBMCs-5µL], Fig. E2 & E3:** BSA is a PCR additive used to mitigate the effect of PCR inhibitors[1] and in single-cell experiments to minimize cell clumping (i.e.,10x Genomics protocols). It also displays crowding properties[2] and can lyse cells when used at high concentration[3]. Replacing Triton X-100 with BSA (1 mg/uL) had different effects in the two tested conditions. In HEK 293T cells (25 µL) it increased the number of detected genes. On the contrary, BSA failed to provide the same benefit in hPBMCs (5 µL). We hypothesize that BSA crowding properties may be beneficial in large reaction volumes but negligible in smaller volumes. It should be noted that BSA strongly interfered with the dispensing process when using the I.DOT. Combining BSA with Triton did not improve the number of detected genes in hPBMCs.

As our main purpose was to develop a reliable and easy-to-implement protocol, we decided to set this condition aside. More tests will be required to determine if this additive can really offer an advantage in particular reaction settings.

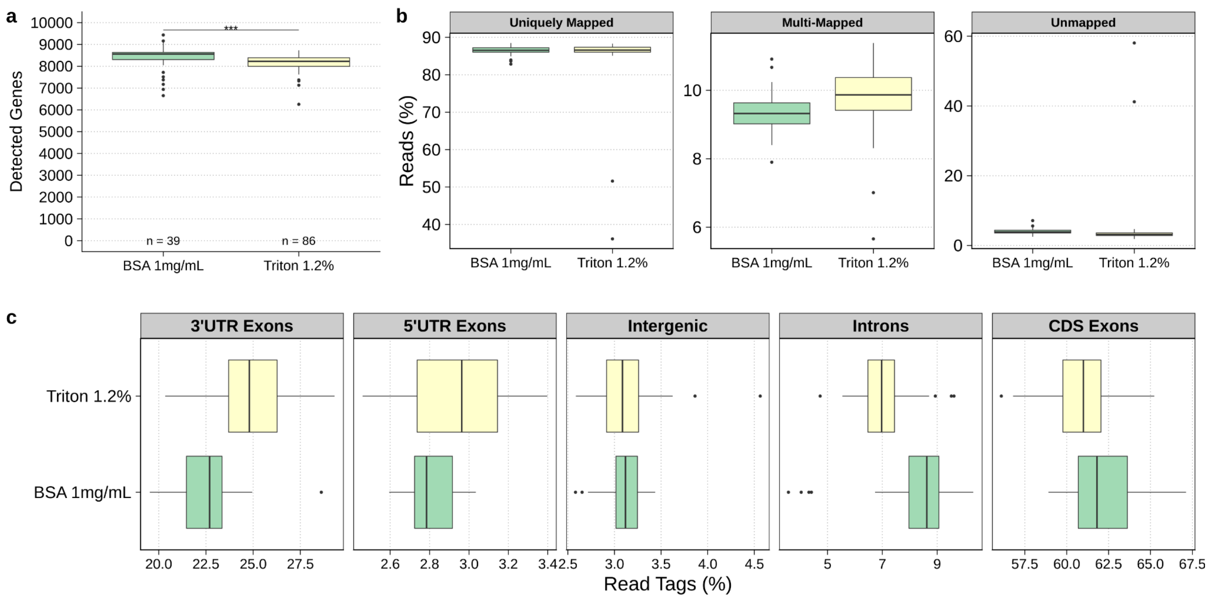


**Fig. E2 | HEK 293T - 25µL - Bovine Serum Albumin (BSA, 1 mg/uL) a.** Number of detected genes (Mann-Whitney U test). **b.** STAR mapping statistics showing the percentage of uniquely mapped, multi-mapped and unmapped reads. **c.** Distribution of mapped reads between introns, intergenic regions or 3'-UTR / 5'-UTR / Coding sequence (=CDS) exons. Expressed in percentage of read tags and computed using ReSQC.

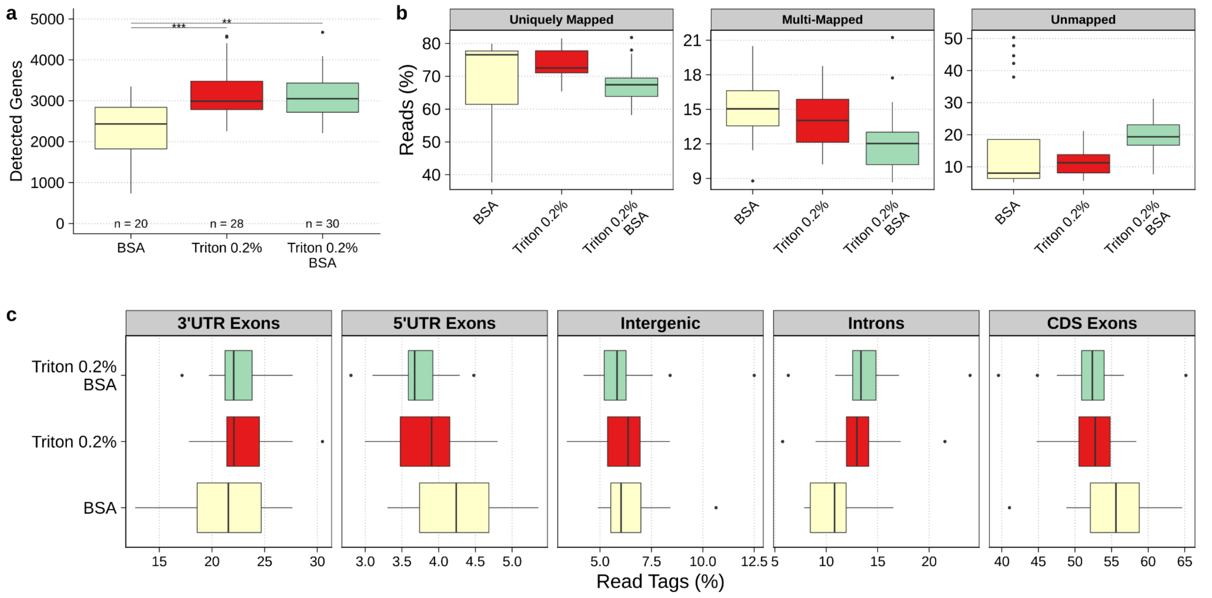


**Fig. E3 | hPBMCs - 25µL - Bovine Serum Albumin (BSA, 1 mg/uL) a.** Number of detected genes (Dunn's test, Bonferonni correction, adj. pval). **b.** STAR mapping statistics showing the percentage of uniquely mapped, multi-mapped and unmapped reads. **c.** Distribution of mapped reads between introns, intergenic regions or 3'-UTR / 5'-UTR / Coding sequence (=CDS) exons. Expressed in percentage of read tags and computed using ReSQC.

**Guanidine Hydrochloride (GuHCl, 250 mM) [HEK-5µL, 100K], Fig. E4:** GuHCl is a chaotropic agent commonly used in DNA/RNA extraction protocols (i.e., Qiagen DNA/RNA extraction kit). Its strong denaturing properties are sufficient to inhibit nucleases, therefore making the addition of RNAse inhibitors in the lysis buffer superfluous, as indirectly confirmed by the average size of pre-amplified cDNA (Fig. E4a). GuHCl could be particularly useful when working with cells undergoing a quick RNA degradation after lysis or which resist Triton X-100 lysis.

Replacing Triton X-100 by 250 mM of GuHCl in the lysis buffer (= 50 mM in final RT-PCR reaction) slightly improved the number of detected genes.

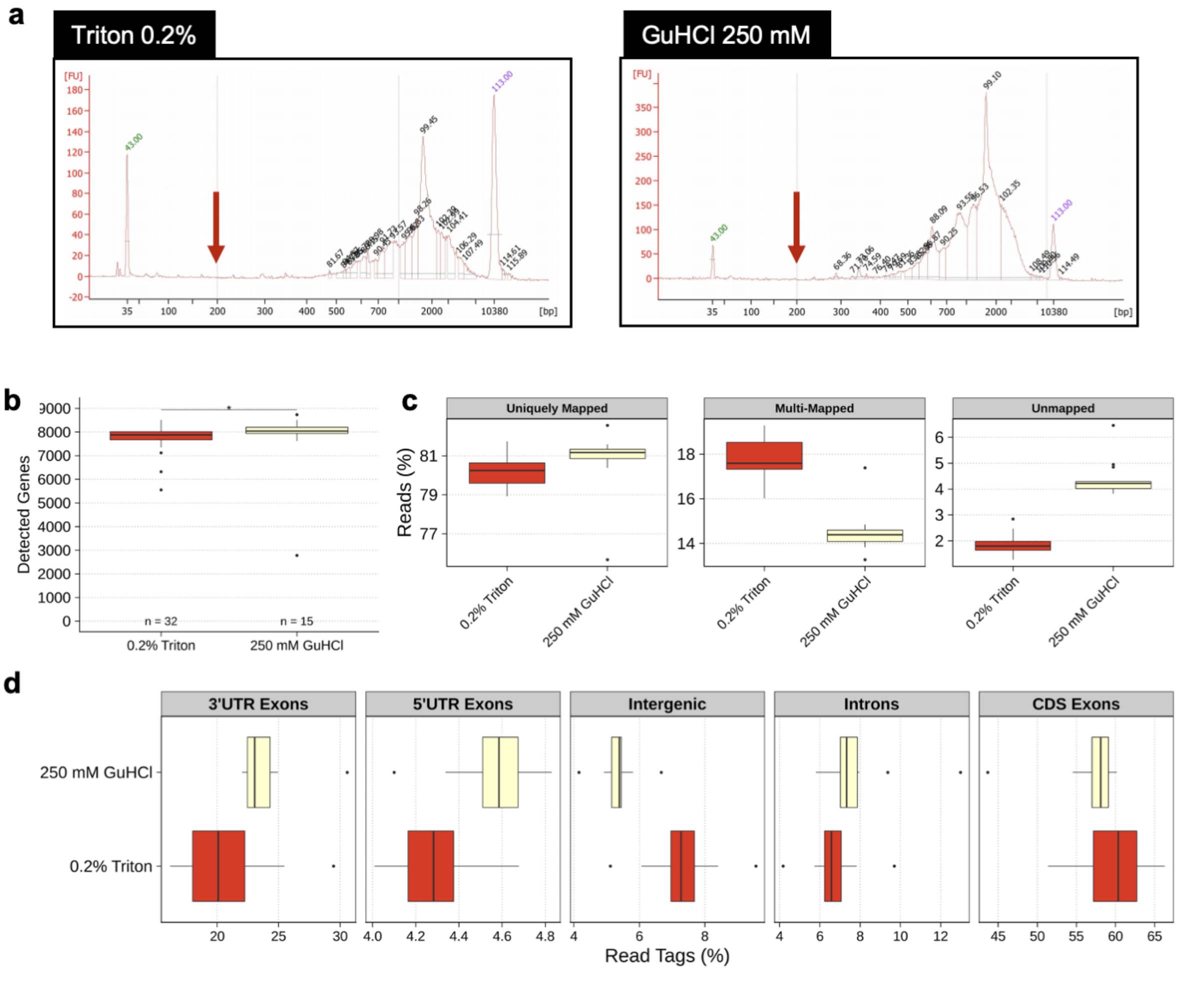


**Fig. E4 | HEK - 5µL - Guanidine Hydrochloride (GuHCl, 250 mM) a.** Bioanalyzer traces of two selected cells lysed with either 0.2% Triton X-100 or 250 mM of GuHCl. Arrows highlight the absence of RNA degradation (<400bp) despite the absence of RNAse inhibitors **b.** Number of detected genes. No significant difference was observed (Mann Withney U test). **c.** STAR mapping statistics showing the percentage of uniquely mapped, multi-mapped and unmapped reads. **d.** Distribution of mapped reads between introns, intergenic regions or 3'-UTR / 5'-UTR / Coding sequence (=CDS) exons. Expressed in percentage of read tags and computed using ReSQC.

#### Denaturation

An initial denaturation step is performed prior to reverse transcription to resolve possible RNA secondary structures. We attempted to improve the denaturation conditions by modifying its duration and temperature.

**3 vs 10 min RNA denaturation (72°C) [hPBMCs-5μL], Fig. E5:** Increasing the denaturation time from 3 to 10 minutes, as reported in other protocols[4], did not affect the number of detected genes. We observed however a small increase in the proportion of intronic reads.

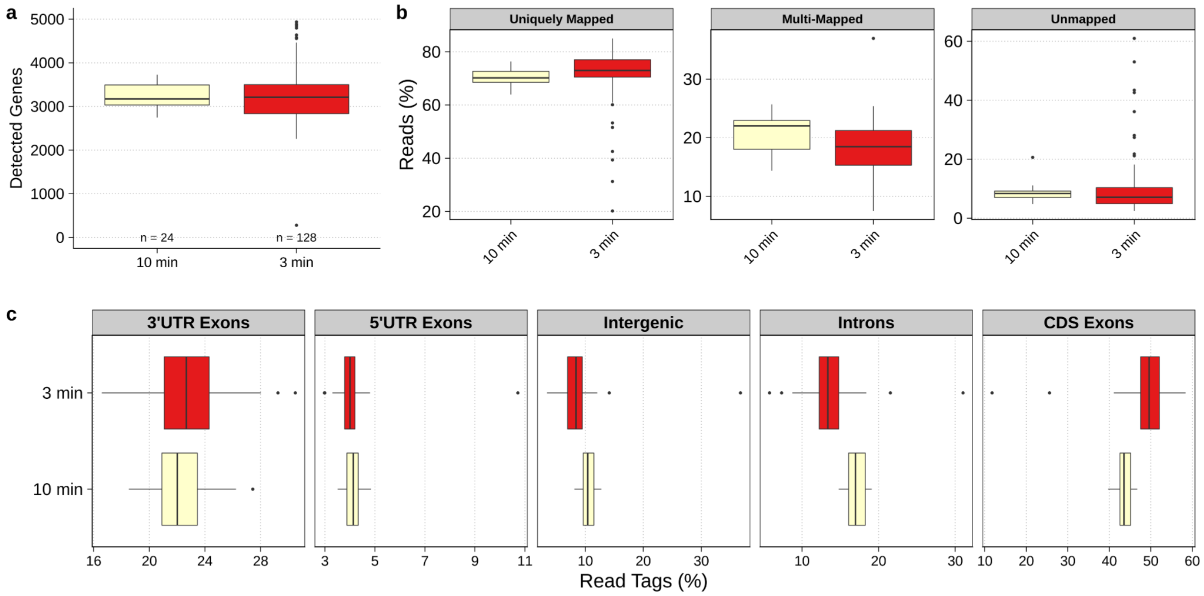


**Fig. E5 | hPBMCs - 5μL - RNA denaturation timing a.** Number of detected genes. No significant difference was observed (Mann-Whitney U test). **b.** STAR mapping statistics showing the percentage of uniquely mapped, multi-mapped and unmapped reads. **c.** Distribution of mapped reads between introns, intergenic regions or 3'-UTR / 5'-UTR / Coding sequence (=CDS) exons. Expressed in percentage of read tags and computed using ReSQC.

**Denaturation temperature (72°C vs 95°C) [hPBMCs-5μL], Fig. E6:** While most scRNA-seq protocols typically perform the denaturation at 72°C, STRT-seq-2i does it at 95°C and reported improved cDNA yields and an increased average cDNA length[5]. In our study, carrying out the denaturation at 95°C resulted in an increased number of multi-mapped reads, a 3'-end gene body coverage bias, a lower average number of detected genes and a striking increase in the proportion of intronic to the detriment of exonic features, compared to the standard 72°C denaturation (Fig. E6). However, when assessing the number of expressed genes using both exonic and intronic features, 95°C denaturation showed greater gene detection (Fig. E6c). Among protein coding genes, the 95°C denaturation detected 2.38-times more genes with only intronic and no exonic reads ($\mu_{onlyIntronics\_72°C}$=429 and $\mu_{onlyIntronics\_95°C}$=1022).

The origin of these intronic reads remains to be confirmed. While we cannot exclude a better denaturation of some mRNAs displaying strong secondary structures or partial lysis of the nuclei releasing pre-mRNA, we currently hypothesize that higher temperatures might result in a mild fragmentation of some mRNAs, as suggested by the 3'-end gene body coverage bias and greater proportion of intergenic reads. Due to these conflicting observations we decided to keep using a standard 72°C denaturation, as RNA fragmentation may significantly bias differential expression, isoform or any UMI-based analysis.

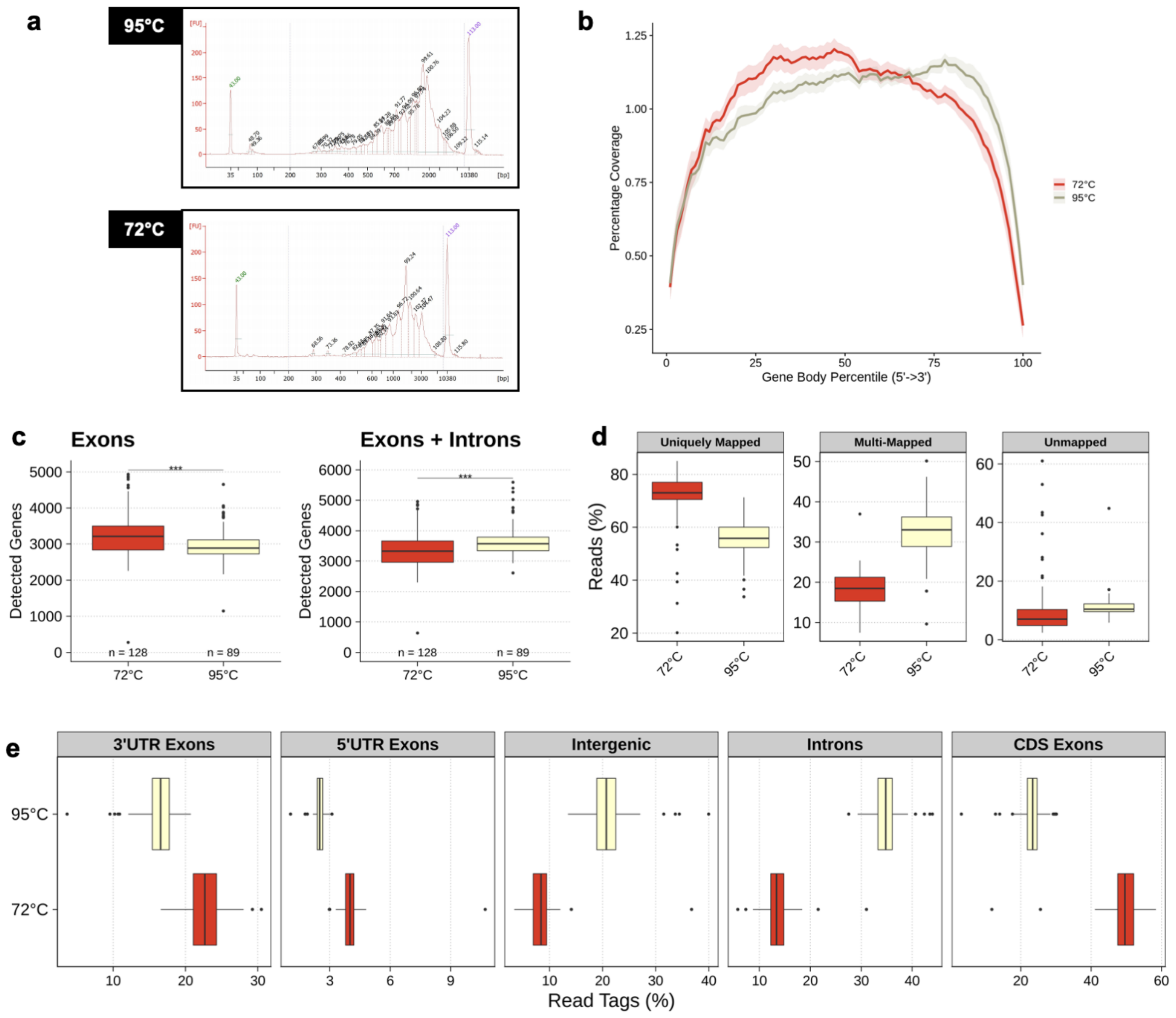


**Fig. E6 | hPBMCs - 5µL - RNA denaturation temperature a.** Bioanalyzer traces of two selected cells with RNA denatured at 95°C or 72°C for 3 minutes. **b.** Gene body coverage **c.** Number of detected genes using exonic (right) or exonic+intronic (left) reads (Mann-Whitney U test). **c.** STAR mapping statistics showing the percentage of uniquely mapped, multi-mapped and unmapped reads. **d.** Distribution of mapped reads between introns, intergenic regions or 3'-UTR / 5'-UTR / Coding sequence (=CDS) exons. Expressed in percentage of read tags and computed using ReSQC.

### Reverse Transcription

During reverse transcription (RT) mRNAs are converted into cDNA by a reverse transcriptase.

**1:5, 1:10, 1:50, FS oligo-dT$_{30}$VN concentration [hPBMCs-5µL], Fig. E7:** When using hPBMCs, FS displayed large amounts of leftover primer dimers which could not be efficiently removed with a magnetic bead cleanup and were long enough to be used as a substrate in the following tagmentation reaction[6,7] (Fig. E7a). FS oligo-dT$_{30}$VN is present in the lysis buffer at a concentration of almost 10 µM. Upon adding the RT-PCR mix (4:1 volume ratio RT-PCR:lysis buffer) the concentration decreases to 2 µM. Reducing the amount of FS oligo-dT$_{30}$VN by 5- or 10-times (0.4 / 0.2 µM final) eliminated the primer dimer peaks in hPBMCs while preserving similar cDNA yields and gene detection (Fig. 7b & 7c). A 50-times reduction in FS oligo-dT$_{30}$VN concentration (0.04 µM final) significantly reduced the cDNA yield and was therefore not pursued further. As larger cells may require larger amounts of FS oligo-dT$_{30}$VN we decided to further use the 0.4 µM final concentration (i.e. 5-times reduction).

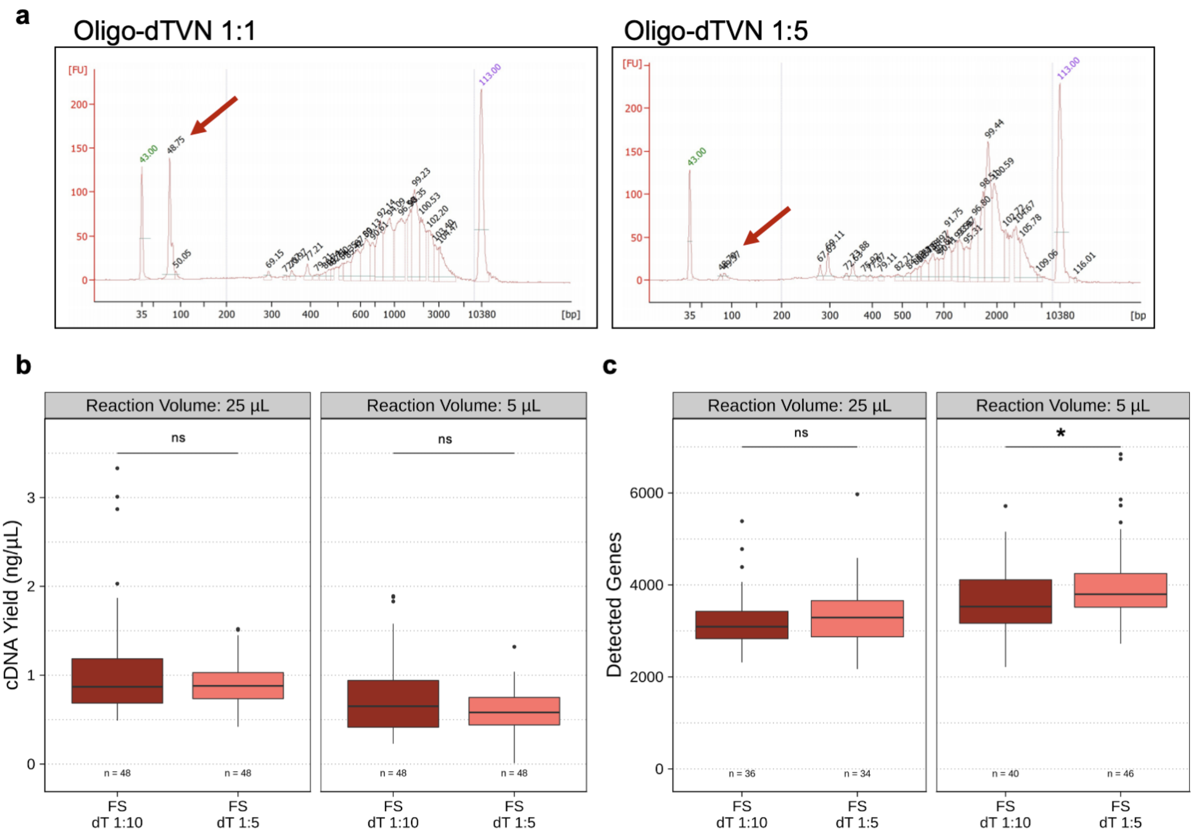

**Fig. E7 | hPBMCs - 5µL or 25µL- FS oligo-dT$_{30}$VN concentration a.** cDNA size distribution of two selected cells. RNA was denatured at 95°C or 72°C for 3 minutes. Final reaction volume of 5µL. The primer dimer peak is highlighted by a red arrow. **b.** FLASH-Seq cDNA yields using 5- or 10-times less FS oligo-dT$_{30}$VN in a 25 or 5 µL reaction (Mann-Whitney U test). **c.** Number of detected genes using 5- or 10-times less FS oligo-dT$_{30}$VN in a 25 or 5 µL reaction (Mann-Whitney U test).

**0, 20, 30, 40 mM NaCl [hPBMCs-5µL], Fig. E8:** NaCl has been shown to improve the activity of the Maxima H- Reverse Transcriptase (ThermoFisher Scientific) in Smart-seq3 protocol[4]. Adding 20 or 30 mM NaCl to the FS mix containing Superscript™ IV did not increase the number of detected genes. 40 mM NaCl inhibited the reaction and was not sequenced.

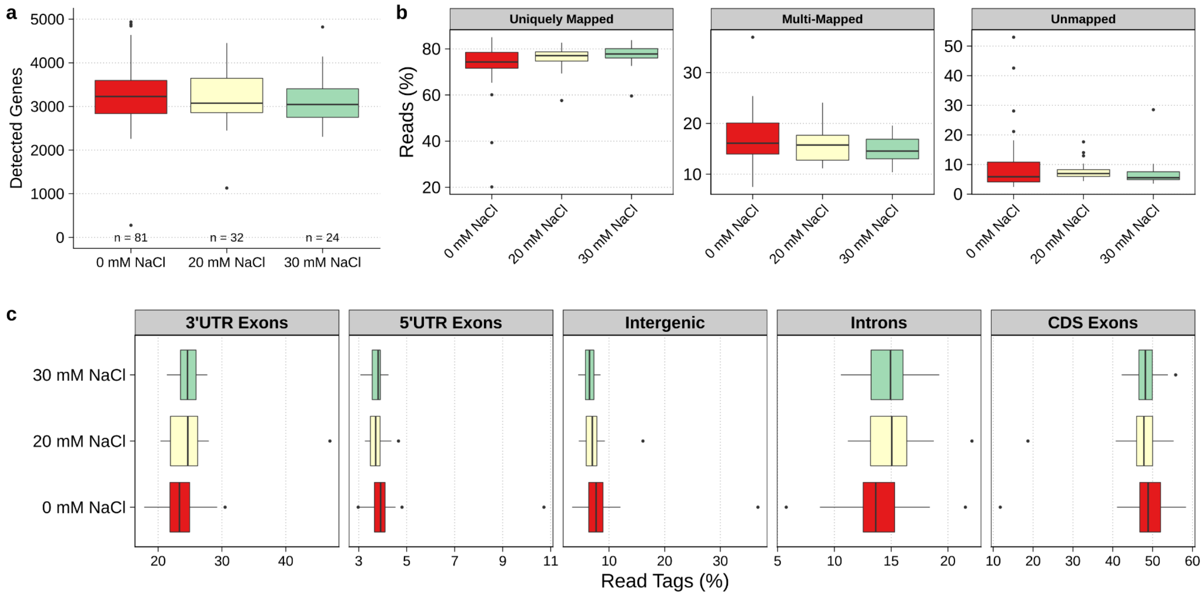


**Fig. E8 | hPBMCs - 25µL - NaCl in RT a.** Number of detected genes (Dunn's test, Bonferonni correction, adj. pval). **b.** STAR mapping statistics showing the percentage of uniquely mapped, multi-mapped and unmapped reads. **c.** Distribution of mapped reads between introns, intergenic regions or 3'-UTR / 5'-UTR / Coding sequence (=CDS) exons. Expressed in percentage of read tags and computed using ReSQC.

**RT Temperature (37°C vs 50°C) [hPBMCs-5µL], Fig. E9:** Moloney Murine Leukemia Virus (MMLV)-derived reverse transcriptases work at a wide range of temperatures (37°C - 55°C). While higher temperatures are generally recommended to resolve RNA secondary structures, we did not observe any significant decrease in the number of detected genes when carrying out the reaction at 37°C. However, the percentage of multi-mapped reads (e.g., ribosomal reads) increased, which could indicate a lower specificity of the poly-A priming of Superscript™ IV at 37°C.

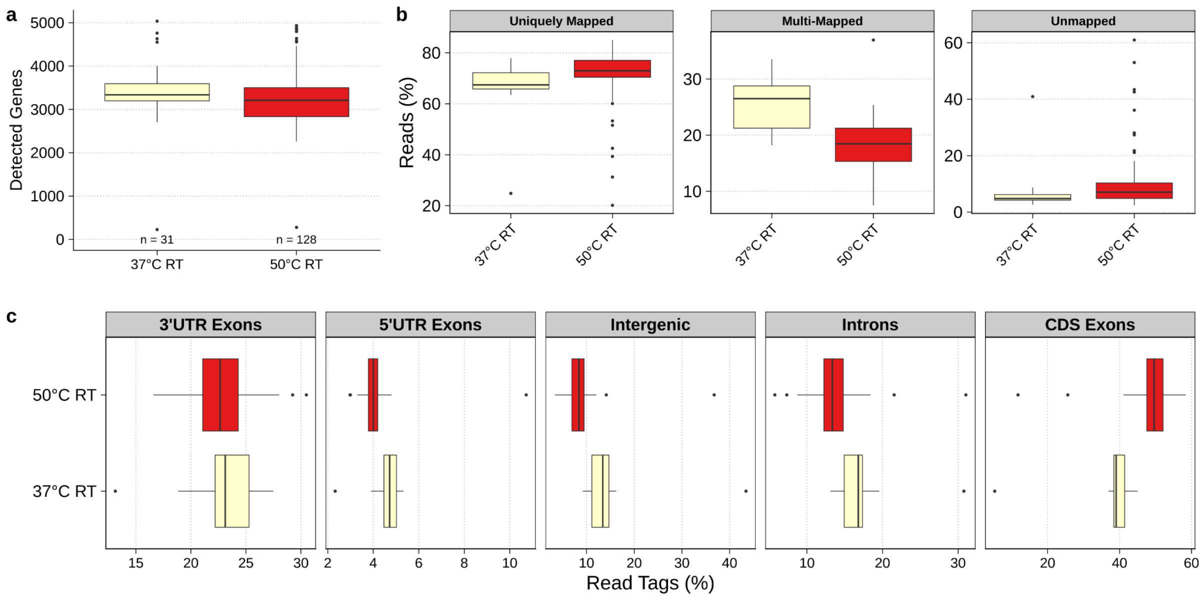


**Fig. E9 | hPBMCs - 5µL - RT Temperature a.** Number of detected genes . No significant difference was observed (Mann-Whitney U test). **b.** STAR mapping statistics showing the percentage of uniquely mapped, multi-mapped and unmapped reads. **c.** Distribution of mapped reads between introns, intergenic regions or 3'-UTR / 5'-UTR / Coding sequence (=CDS) exons. Expressed in percentage of read tags and computed using ReSQC.

**RT Enzyme (Superscript™ IV vs Maxima H-) [hPBMCs-5µL], Fig. E10:** Two of the currently most widely used reverse transcriptases in scRNA-seq are Superscript™ IV and Maxima H Minus. Both perform well in the FS buffer and we did not observe a significant difference in gene detection between them.

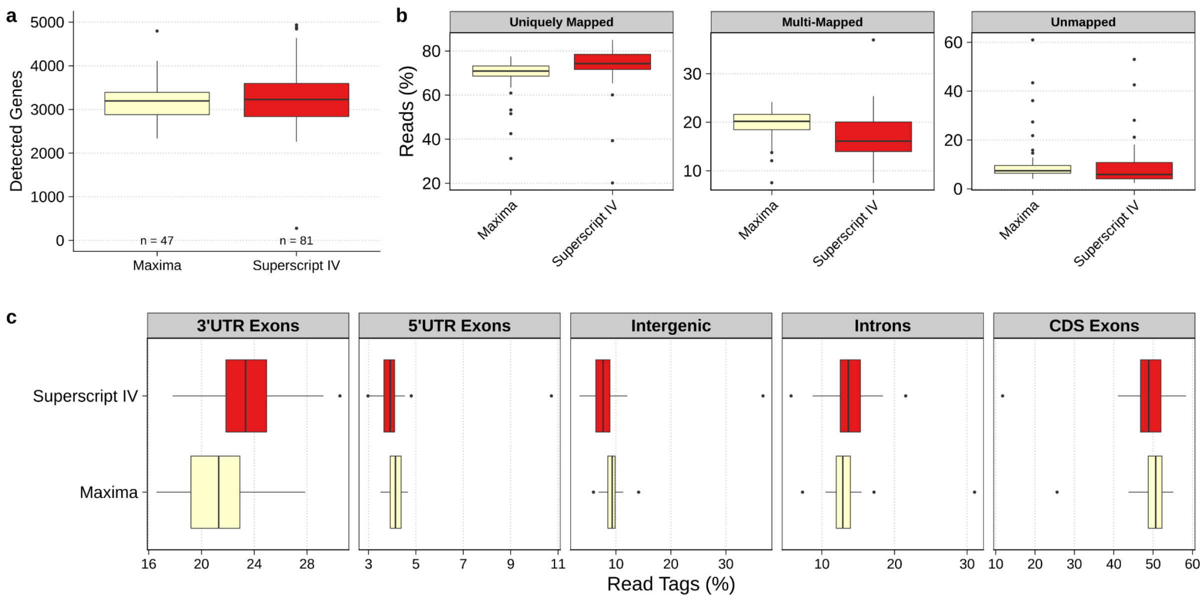

**Fig. E10 | hPBMCs - 5µL - RT Enzyme a.** Number of detected genes. No significant difference was observed (Mann-Whitney U test). **b.** STAR mapping statistics showing the percentage of uniquely mapped, multi-mapped and unmapped reads. **c.** Distribution of mapped reads between introns, intergenic regions or 3'-UTR / 5'-UTR / coding sequence (=CDS) exons. Expressed in percentage of read tags and computed using ReSQC.

**Template-Switching Oligonucleotide (TSO) [hPBMCs-5µL], 250K raw reads, Fig. E11:** Several MMLV-based reverse transcriptases display a template switching activity. Upon reaching the 5´-end of the mRNA molecule, these enzymes add a short stretch of untemplated nucleotides - generally CCC - at the 5'-end of the newly synthesized molecule. These nucleotides are then used as an anchor for the annealing of a second oligo present in the RT mix, the TSO. Reverse transcriptases are then capable of switching template: while initially using the mRNA as template to generate a complementary DNA molecule, they later use the DNA-based template-switching oligonucleotide to generate an additional portion of cDNA, covalently linked to the cDNA derived from the mRNA. The end result is that all cDNA molecules carry a known sequence both at the 5'-end and the 3´-end (the handle sequence located upstream of the poly-T sequence in FS oligo-d$T_{30}$VN primer)[8]. However, the very short and variable length as well as the varying base composition of these untemplated nucleotides makes the annealing very inefficient[9] and a large excess of TSO is usually required to ensure that as many mRNA molecules as possible are efficiently converted to cDNA[10].

Similar to SS2, FS is a highly robust protocol that can accommodate some variations in concentration of all its reagents with negligible efficiency losses except, in our experience, too large shifts in TSO concentration. For this reason, we titrated the amount of TSO required to efficiently process hPBMCs, by comparing the gene detection when using 1-, 1.5-, 2-, 3-, 4- or 5-times more TSO, where 1x TSO corresponds to a final concentration in the RT-PCR reaction of about 2 uM. We observed a linear increase in the number of genes detected with increasing TSO concentrations (median increase = 0-8.74%). However, this improvement was accompanied by a significantly higher proportion of multi-mapped reads, at the expense of the exonic features. The increase in intergenic and multi-mapped reads generated a cumulated excess of up to ~17% of unusable raw reads (4x TSO, about 8 µM). In addition, the increased percentage of intergenic reads may indicate some level of strand-invasion. We therefore decided to not investigate this condition further. We hypothesize that larger cells (i.e., HEK 293T, not tested) may react differently to the variation in TSO concentration due to their higher RNA content.

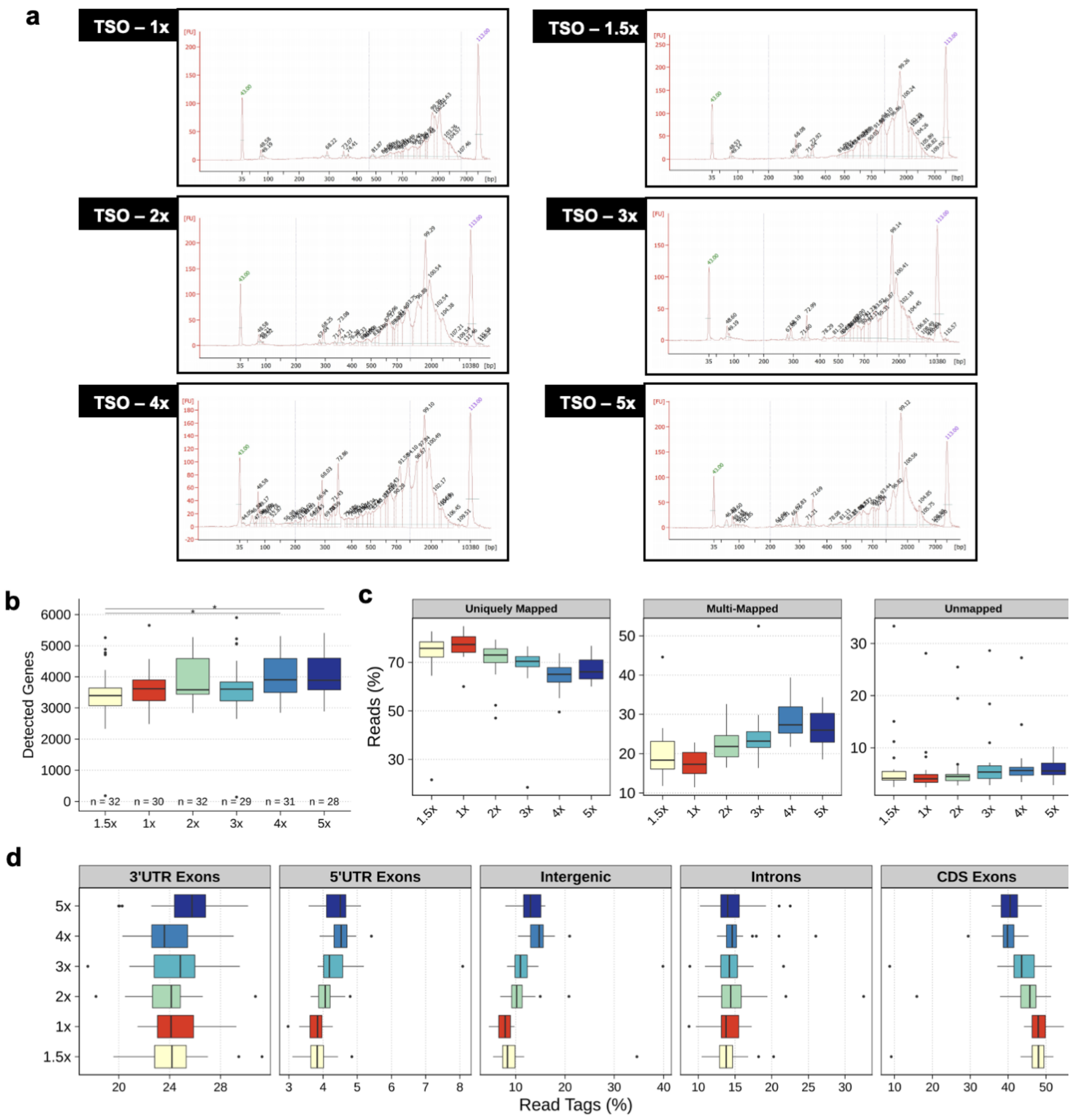


**Fig. E11 | hPBMCs - 5µL - TSO Titration a.** Bioanalyzer traces of selected cells. **b.** Number of detected genes (Dunn's test, Bonferroni correction, adj. pval). **c.** STAR mapping statistics showing the percentage of uniquely mapped, multi-mapped and unmapped reads. **d.** Distribution of mapped reads between introns, intergenic regions or 3'-UTR / 5'-UTR / Coding sequence (=CDS) exons. Expressed in percentage of read tags and computed using ReSQC.

**Ficoll-400 4% v/v [hPBMCs-5µL], Fig. E12:** Ficoll-400 is a highly branched polymer formed by the copolymerization of sucrose and epichlorohydrin which displays crowding properties[2]. Due to its high molecular weight Ficoll-400 mimics the high concentration of molecules in the intracellular environment. Macromolecular crowding is also one of the key factors for the higher efficiency of biological reactions inside cells in comparison to laboratory tubes. The addition of Ficoll-400 to a final concentration of 4% w/v did not improve the reaction. Please note however that, due to volume constraints, betaine (generally present in the RT-PCR at 1 M final concentration) had to be removed to make room for Ficoll-400. Betaine is one of the key additives in the SS2 protocol[10]. These results could suggest that the effects of Ficoll-400 could compensate for the loss of betaine (See Fig. E13).

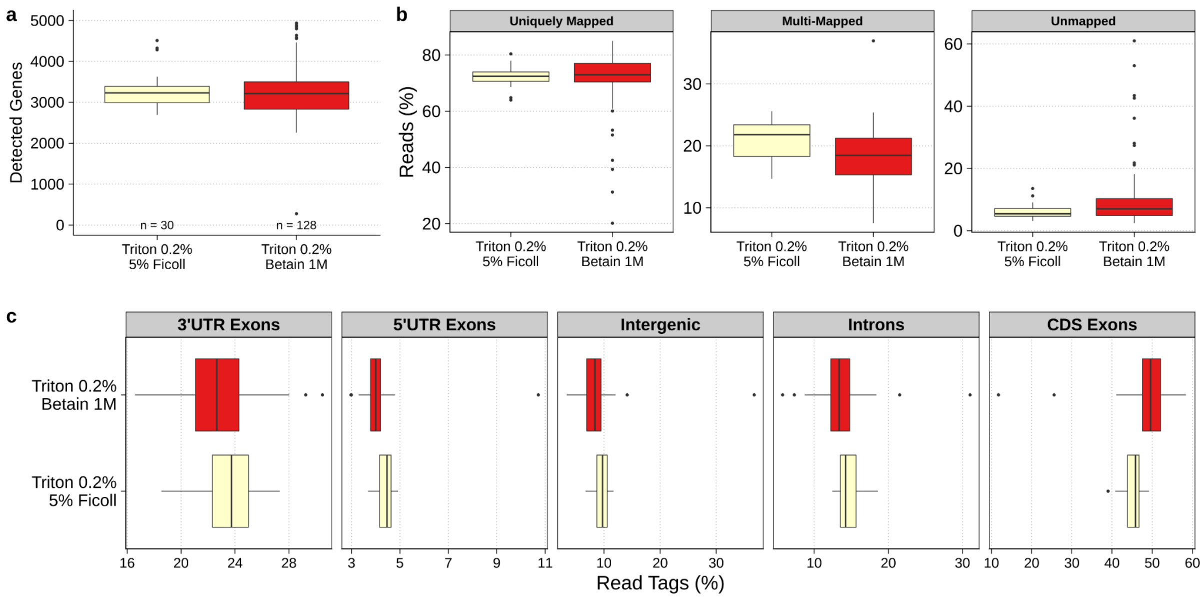


**Fig. E12 | hPBMCs - 5µL - Ficoll-400 4% v/v** **a.** Number of detected genes. No significant difference was observed (Mann-Whitney U test). **b.** STAR mapping statistics showing the percentage of uniquely mapped, multi-mapped and unmapped reads. **c.** Distribution of mapped reads between introns, intergenic regions or 3'-UTR / 5'-UTR / Coding sequence (=CDS) exons. Expressed in percentage of read tags and computed using ReSQC.

**Betaine (1M Betaine vs 0M Betaine) [HEK-5µL], Fig. E13:** Betaine (N,N,N-trimethylglycine) is a methyl group donor which increases RT efficiency, mitigates RNA refolding during RT and acts as a cryoprotectant[10]. The addition of 1 M betaine was one of the key additives introduced by SS2[10]. Surprisingly, we did not observe a significant difference in the number of detected genes when performing FS with 4% v/v Ficoll (without betaine, Fig. E12) and the 1 M betaine control. These initial results suggested that either the addition of 4% v/v Ficoll could compensate for the loss of betaine or that the addition of betaine is no longer needed for FS. To test these hypotheses we processed HEK 293T cells with or without betaine. The number of detected genes decreased when betaine was omitted, showing that its addition is still required in the FS protocol. It also suggests that betaine may be replaced by 4% v/v Ficoll.

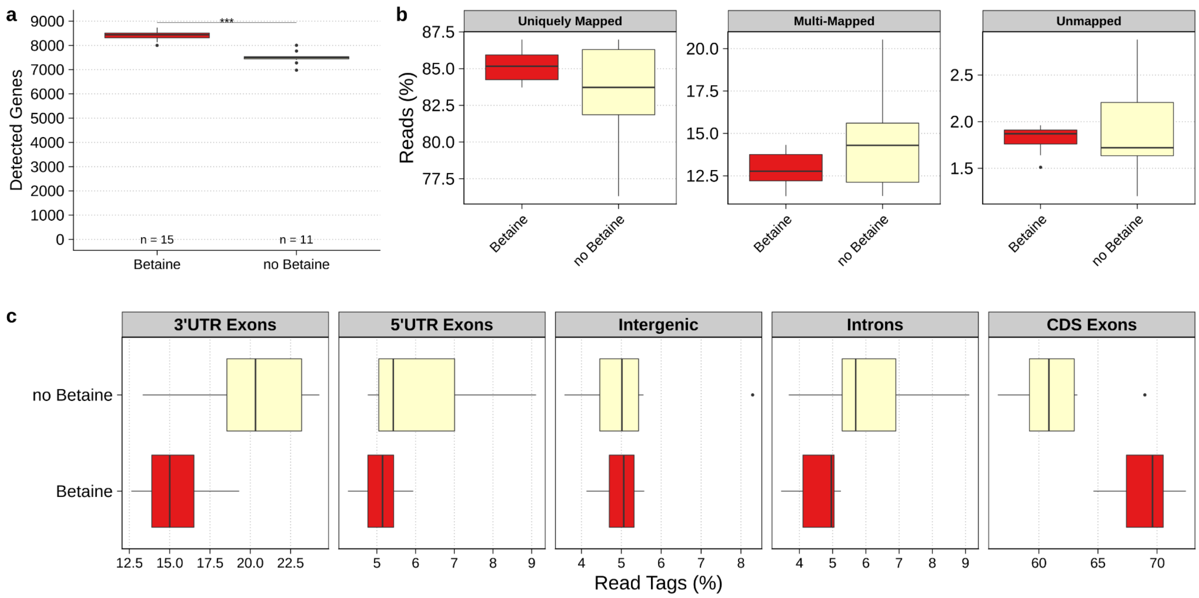


**Fig. E13 | HEK - 5µL - Betaine a.** Number of detected genes. The addition of betaine increases the number of detected genes (Mann-Whitney U test). **b.** STAR mapping statistics showing the percentage of uniquely mapped, multi-mapped and unmapped reads. **c.** Distribution of mapped reads between introns, intergenic regions or 3'-UTR / 5'-UTR / coding sequence (=CDS) exons. Expressed in percentage of read tags and computed using ReSQC.

### cDNA Amplification

As mentioned above, successfully reverse transcribed cDNA molecules contain known sequences both at the 5'- and 3'-end. This allows the amplification of cDNA by PCR using a single primer (i.e. the end of both the TSO and the FS oligo-dT$_{30}$VN primer is identical), in order to reach a concentration compatible with the downstream reactions.

**Pfu DNA Polymerase (0.25-0.375 U/reaction) [hPBMCs-5μL], Fig. E14:** Addition of *Pyrococcus furiosus* DNA Polymerase (a high-fidelity DNA polymerase, NEB) to the RT-PCR mix did not significantly increase the number of detected genes.

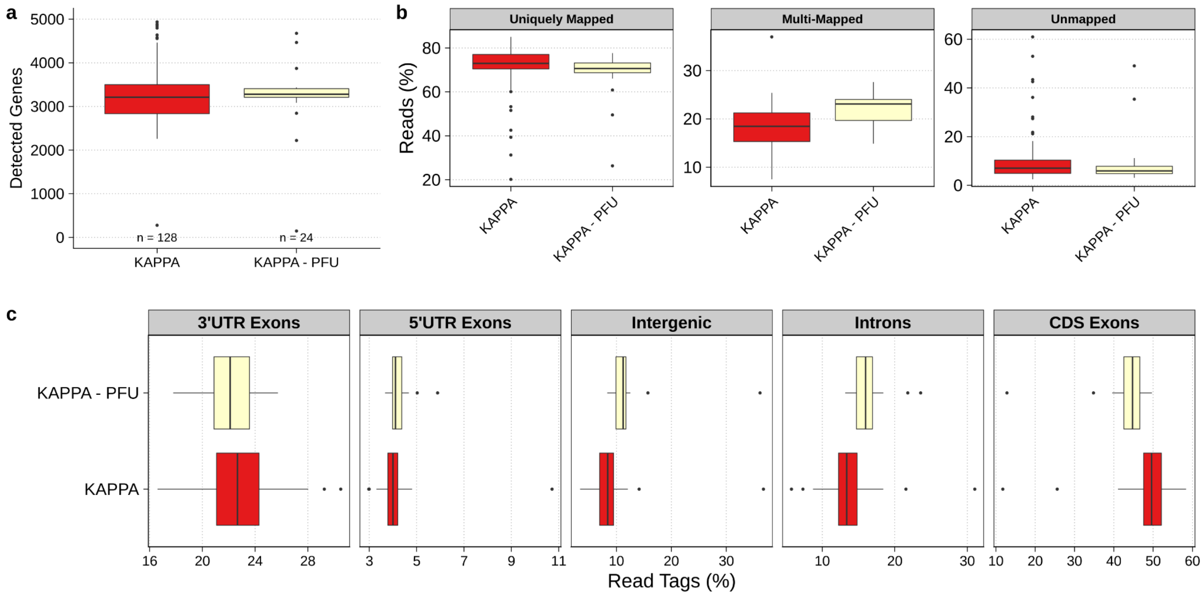


**Fig. E14 | hPBMCs - 5μL - Pfu DNA Polymerase a.** Number of detected genes. No significant difference was observed (Mann-Whitney U test). **b.** STAR mapping statistics showing the percentage of uniquely mapped, multi-mapped and unmapped reads. **c.** Distribution of mapped reads between introns, intergenic regions or 3'-UTR / 5'-UTR / Coding sequence (=CDS) exons. Expressed in percentage of read tags and computed using ReSQC.

**PCR elongation time (4 vs 6 min) [hPBMCs-5μL], Fig. E15:** While other methods (i.e., SS3[4], SSsc [Takara Bio]) perform a 4-minutes elongation step in the pre-amplification reaction, we observed a greater number of detected genes when using 6 minutes.

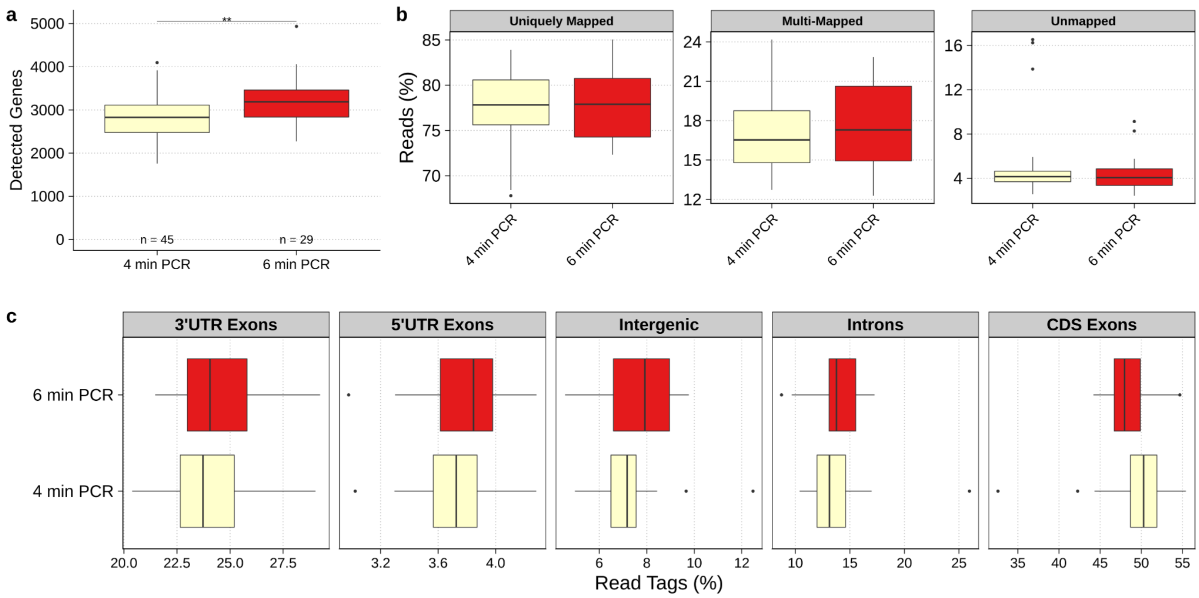


**Fig. E15 | hPBMCs - 5µL - PCR Elongation Time a.** Number of detected genes (Mann-Whitney U test). **b.** STAR mapping statistics showing the percentage of uniquely mapped, multi-mapped and unmapped reads. **c.** Distribution of mapped reads between introns, intergenic regions or 3'-UTR / 5'-UTR / coding sequence (=CDS) exons. Expressed in percentage of read tags and computed using ReSQC.

**Extreme Thermostable Single-Stranded DNA Binding Protein (ET SSB, NEB: 0, 40 ng, 80 ng, 160 ng, 250 ng) [hPBMCs-5µL], Fig. E16:** ET SSB is reported to be a PCR additive that enhances DNA polymerase activity and increases PCR yield. It remains active even at high temperatures and for prolonged periods (i.e., 95°C, 60 minutes). In our hands the addition of ET SSB did not improve cDNA yield nor the number of recovered genes. Using 250 ng ET SSB/samples appeared to inhibit the reaction altogether.

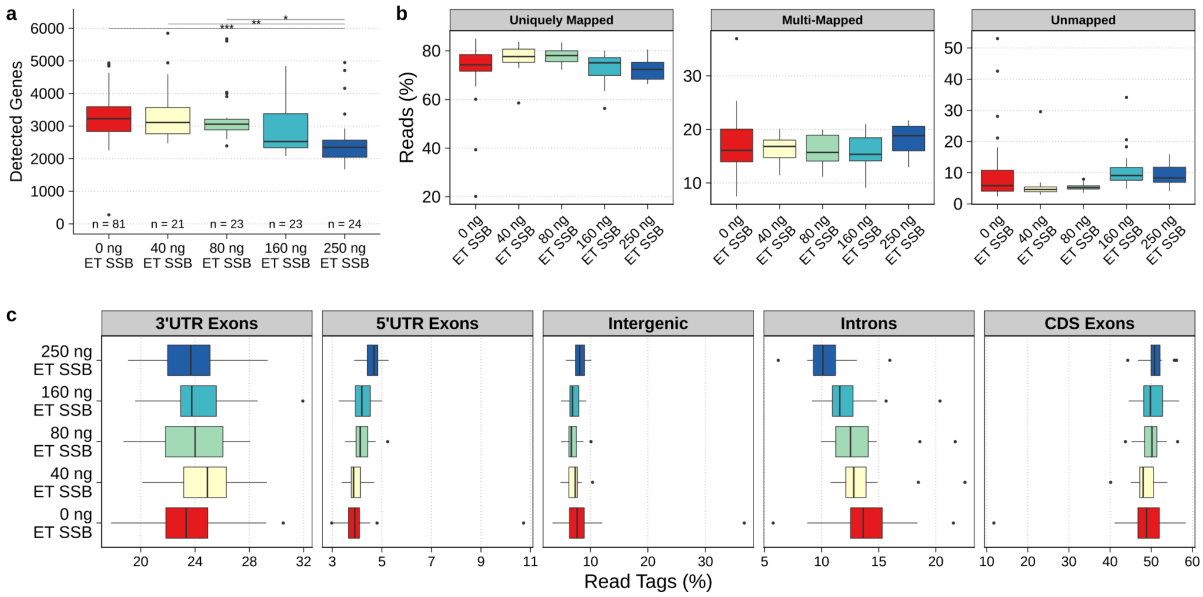

**Fig. E16 | hPBMCs - 5µL - ET SSB Titration a.** Number of detected genes (Dunn's test, Bonferroni correction, adj. pval). **b.** STAR mapping statistics showing the percentage of uniquely mapped, multi-mapped and unmapped reads. **c.** Distribution of mapped reads between introns, intergenic regions or 3'-UTR / 5'-UTR / coding sequence (=CDS) exons. Expressed in percentage of read tags and computed using ReSQC.

**Extra dNTPs (1.2 mM dNTPs vs no additional dNTP) [HEK-5µL], 50K raw reads, Fig. E17:** the dNTPs used in the FS protocol come from two different sources: the lysis buffer mix (final concentration of 1.2 mM in RT-PCR) and the 2x KAPA HiFi HotStart Ready Mix (0.3 mM each in the final 1x mix). The addition of dNTPs to the lysis buffer had previously been shown to stabilize the mRNA during denaturation, however the minimum amount required for an efficient RT reaction was never tested[10].

Therefore, we sequenced HEK 293T cells processed with either 1.5 mM dNTPs (1.2 mM + 0.3 mM in Fig. E17, "Full dNTPs") or just the 0.3 mM from the KAPA Ready Mix (Fig. E17, "No dNTPs"). We observed a significant reduction in the number of detected genes when omitting dNTPs in the lysis buffer, indicating that extra dNTPs must be added to the lysis buffer to achieve the best results.

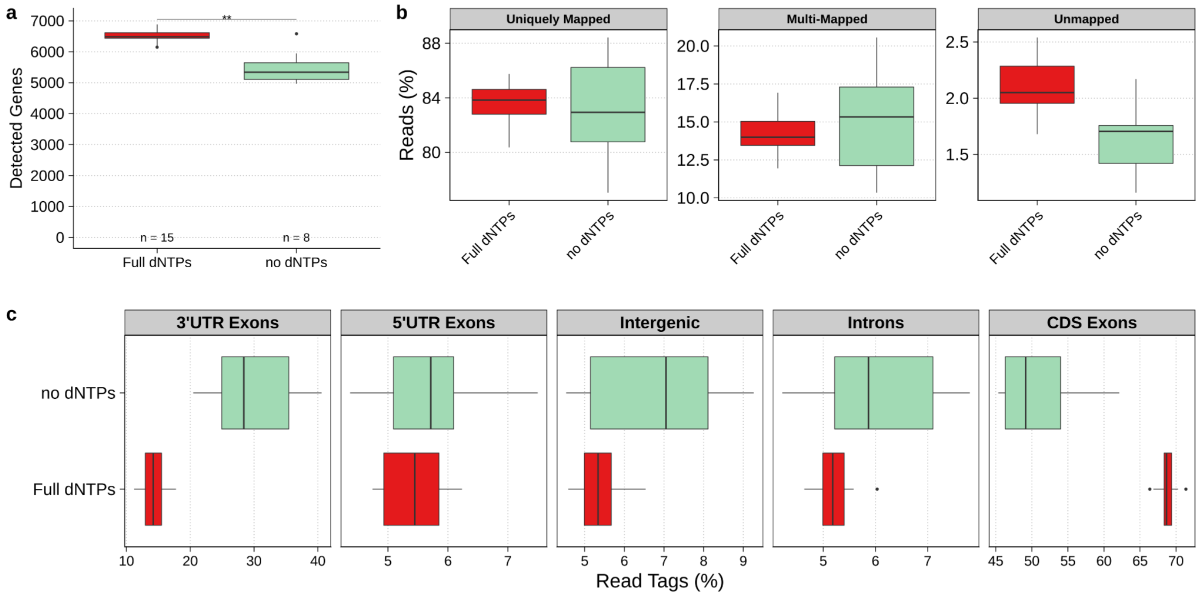


**Fig. E17 | HEK - 5µL - Extra dNTPs a.** Number of detected genes (Mann-Whitney U test). **b.** STAR mapping statistics showing the percentage of uniquely mapped, multi-mapped and unmapped reads. **c.** Distribution of mapped reads between introns, intergenic regions or 3'-UTR / 5'-UTR / coding sequence (=CDS) exons. Expressed in percentage of read tags and computed using ReSQC.

#### T4 gene 32 protein (T4g32p)

T4g32p is a single-strand binding (SSB) protein derived from the T4 bacteriophage, which stabilizes single-strand DNA and RNA and has been shown to enhance the SMART reaction efficiency in the first iteration of the method[11]. More recently, it has been used in the single-cell protocol “RAMDA-seq” to promote strand-displacement and protect newly synthesized cDNA from exonuclease treatment[12]. This protein requires the RT reaction to be carried out at 37°C instead of 50°C. The addition of 5 µg of T4g32p (NEB) to the FS RT-PCR had a dramatic impact on the reaction when it was carried out in a final volume of 25 µL. It increased the cDNA yield by 2.8-fold in HEK 293T cells and by 4.3-fold in hPBMCs (Fig. E18a). However, this was accompanied by a significant increase in shorter cDNA fragments, which could be eliminated in the following bead cleanup only by decreasing the cDNA / magnetic bead ratio from the standard 0.8x to 0.6x (Fig. E18b). The addition of 5 µg T4g32p to the reaction had a positive impact on the number of detected genes in hPBMCs (Fig. E18c). Unfortunately, it also increased the percentage of multi-mapped and unmapped reads (Fig. E18d.). The origin of multi-mapped reads can likely be traced back to the lower temperature at which the RT is carried out compared to standard FS (see Fig. E9), thus coupling a lower stringency with a higher reaction efficiency. However, we were unable to assess the source of the unmapped reads. We did not observe an increase in cDNA yield when performing the RT reaction at 50°C in the presence of T4g32p (data not shown), suggesting that T4g32p is inactivated at higher temperatures. Interestingly, T4g32p increased only mildly the cDNA yield when the reaction was carried out in a smaller volume (5 µL), with no improvements in the gene detection (see T4g32p titration in hPBMCs, Fig. E19). We made several attempts to modify the reaction conditions without any apparent improvement on the number of detected genes. We therefore concluded that the benefits of miniaturization may compensate for the effect of T4g32p.

Finally, T4g32p increases the cost per 96-cells by ~230$ which could be better used in sequencing deeper each cell. We therefore do not recommend its use.

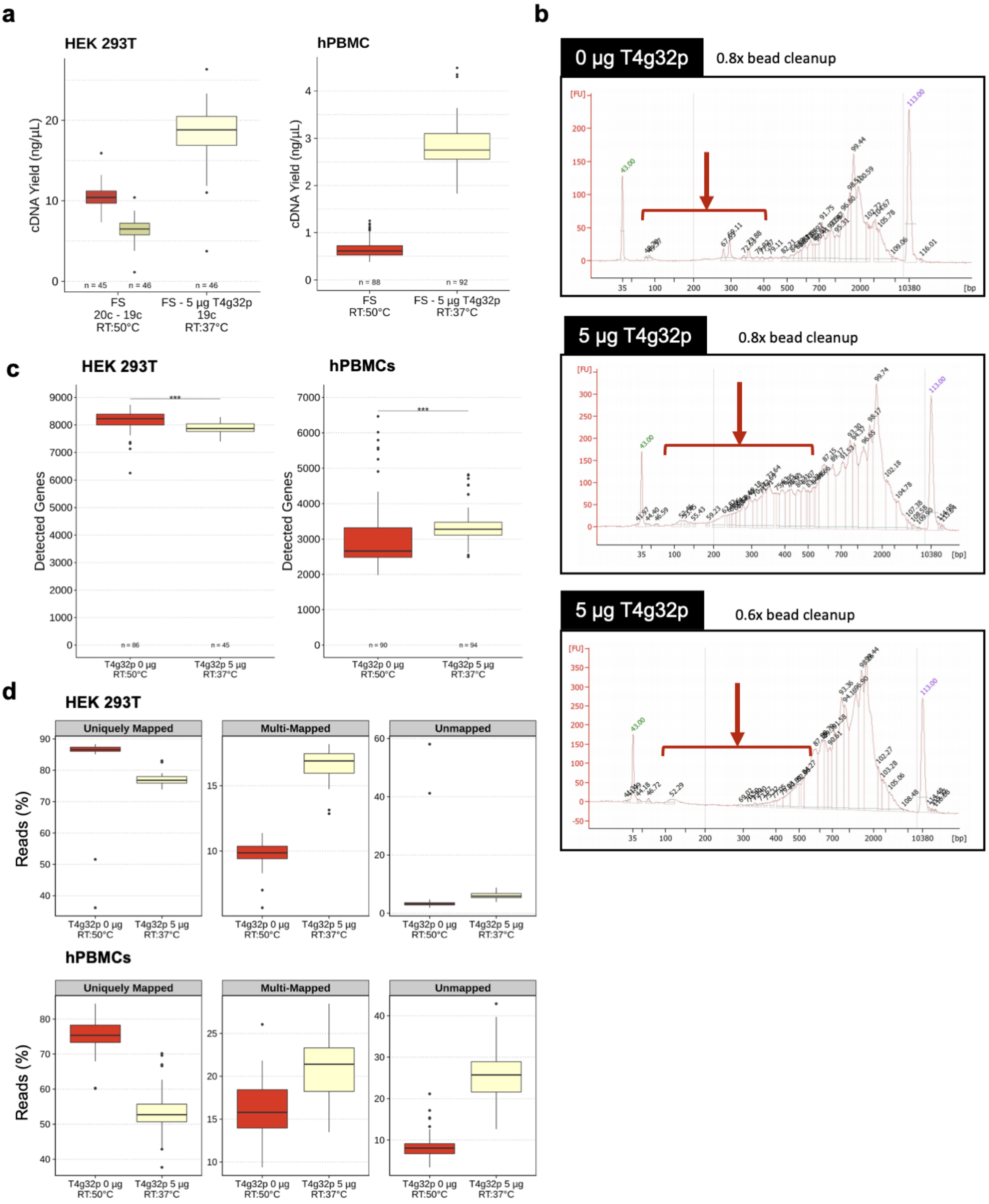


**Fig. E18 | hPBMCs & HEK 293T - 25 µL - Effect of T4g32p on FS reaction. a.** FS cDNA yield of HEK 293T (left) or hPBMCs (right) processed in the presence or absence of 5 µg T4g32p. Final reaction volume of 25 µL. **b.** Bioanalyzer traces of selected cells processed in the absence of T4g32p (top) or with 5 µg T4g32p and cleaned using a ratio of magnetic beads / cDNA of either 0.8x (mid) or 0.6x (bottom). Red arrows highlight the excess of short fragments (<550bp). **c.** Number of detected genes in either HEK 293T (left) or hPBMCs (right) in the presence or absence of T4g32p protein (Mann-Whitney U test). Final reaction volume of 25 µL. **d.** STAR mapping statistics showing the percentage of uniquely mapped, multi-mapped and unmapped reads in either HEK 293T (left) or hPBMCs (right) in the presence or absence of T4g32p protein.

**T4g32p Titration [hPBMCs-5µL], Fig. E19:** The amount of T4g32p was titrated in a 5 µL reaction by adding 0.125, 0.25, 0.50, 0.75 or 1 µg of T4g32p to the reaction and comparing the results with regular FS (no T4g32p, RT-50°C). In contrast with the 25 µL reaction volume, the addition of T4g32p in a smaller reaction volume did not significantly increase the number of detected genes. These results seem to suggest that in a final reaction volume of 5 µL, the addition of T4g32p may not be required.

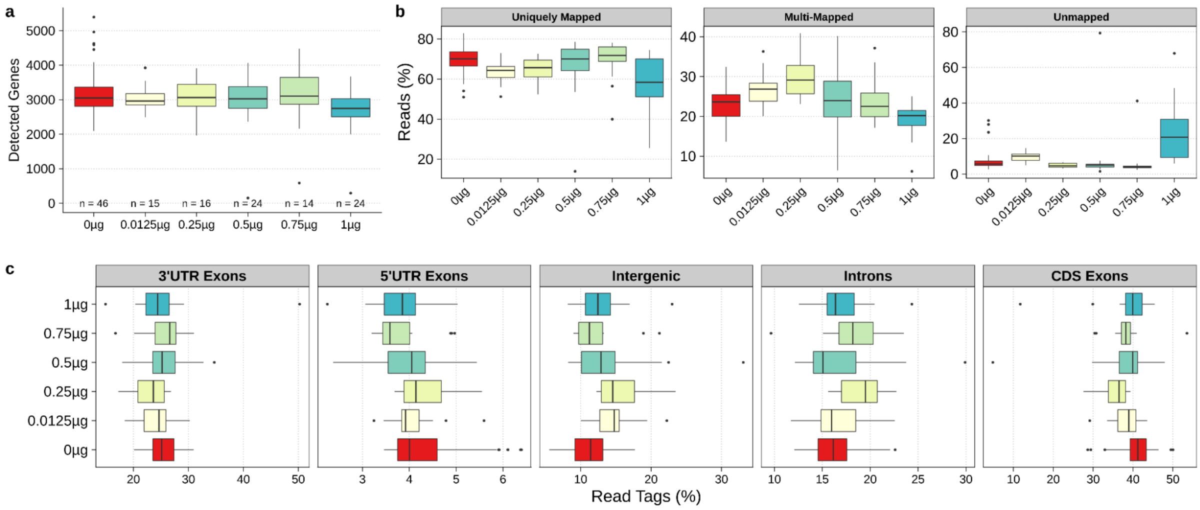


**Fig. E19 | hPBMCs - 5µL - T4g32p Titration a.** Number of detected genes (Dunn's test, Bonferroni correction, adj. pval). **c.** STAR mapping statistics showing the percentage of uniquely mapped, multi-mapped and unmapped reads. **d.** Distribution of mapped reads between introns, intergenic regions or 3'-UTR / 5'-UTR / coding sequence (=CDS) exons. Expressed in percentage of read tags and computed using ReSQC.

**T4g32p 0.25 µg + FS oligo-dT$_{30}$VN (1/5 or 1/10 compared the standard FS [1.8µM]) [hPBMCs-5µL], Fig. E20:** Similarly to the reaction in absence of T4g32p, we did not observe a difference in terms of number of detected genes when using 5- or 10-times less FS oligo-dT$_{30}$VN. We decided to continue working with 5-times less FS oligo-dT$_{30}$VN as it ensures a more complete usage of the primers in RT without undesired side-effects (i.e. presence of a primer dimer peak) after clean-up.

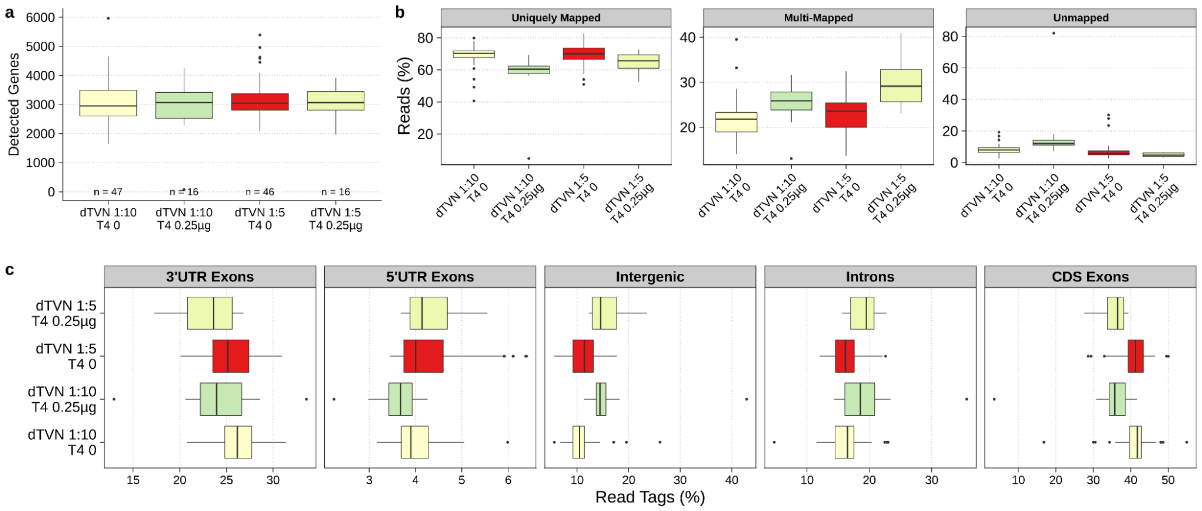


**Fig. E20 | hPBMCs - 5µL - T4g32p and FS oligo-dT$_{30}$VN a.** Number of detected genes (Dunn's test, Bonferroni correction, adj. pval). **c.** STAR mapping statistics showing the percentage of uniquely mapped, multi-mapped and unmapped reads. **d.** Distribution of mapped reads between introns, intergenic regions or 3'-UTR / 5'-UTR / coding sequence (=CDS) exons. Expressed in percentage of read tags and computed using ReSQC.

**T4g32p 0.25/0.50/0.75 µg + Maxima H- or Superscript IV [hPBMCs-5µL], Fig. E21:** Similarly to the reaction in absence of T4g32p, we did not observe a significant difference between the two reverse transcriptases.

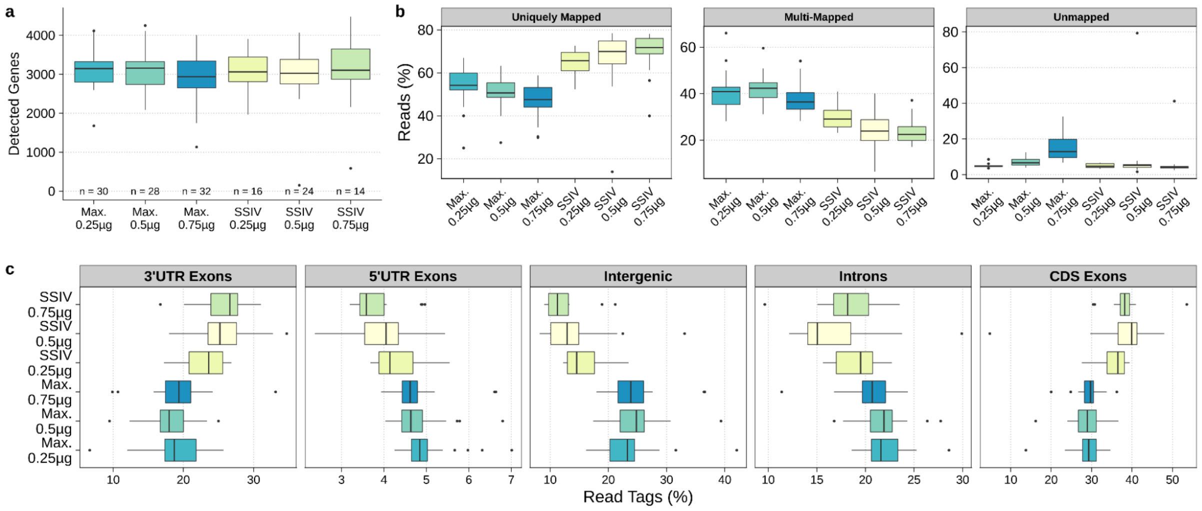


**Fig. E21 | hPBMCs - 5µL - T4g32p Titration and Retrotranscriptase a.** Number of detected genes (Dunn's test, Bonferroni correction, adj. pval). **c.** STAR mapping statistics showing the percentage of uniquely mapped, multi-mapped and unmapped reads. **d.** Distribution of mapped reads between introns, intergenic regions or 3'-UTR / 5'-UTR / coding sequence (=CDS) exons. Expressed in percentage of read tags and computed using ReSQC.

**T4g32p 0.25/0.5/0.75 µg + variable RT duration (30 or 60 minutes) [hPBMCs-5µL], Fig. E22:** We observed a trend towards a lower number of detected genes when performing the RT for 30 minutes with T4g32p compared to 60 minutes. It should be noted that the 30 minutes RT were indexed using plexWell standard input kit (SeqWell) while the 60 minutes RT were indexed using Nextera XT (Illumina).

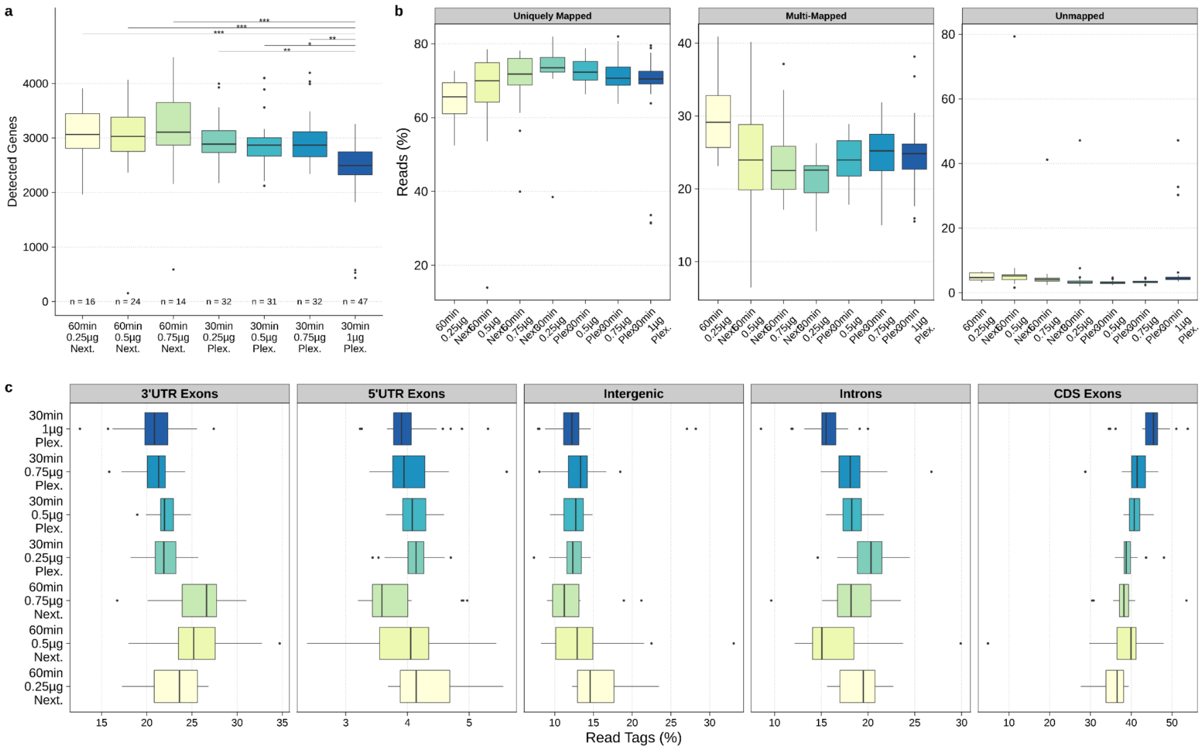


**Fig. E22 | hPBMCs - 5µL - T4g32p Titration and RT Duration a.** Number of detected genes (Dunn's test, Bonferroni correction, adj. pval). **c.** STAR mapping statistics showing the percentage of uniquely mapped, multi-mapped and unmapped reads. **d.** Distribution of mapped reads between introns, intergenic regions or 3'-UTR / 5'-UTR / coding sequence (=CDS) exons. Expressed in percentage of read tags and computed using ReSQC.

**T4g32p 0.25/0.5/1 µg + Triton X-100 1.2% or 0.2% [hPBMCs-5µL], Fig. E23:** We observed greater gene detection when using 0.2% Triton X-100.

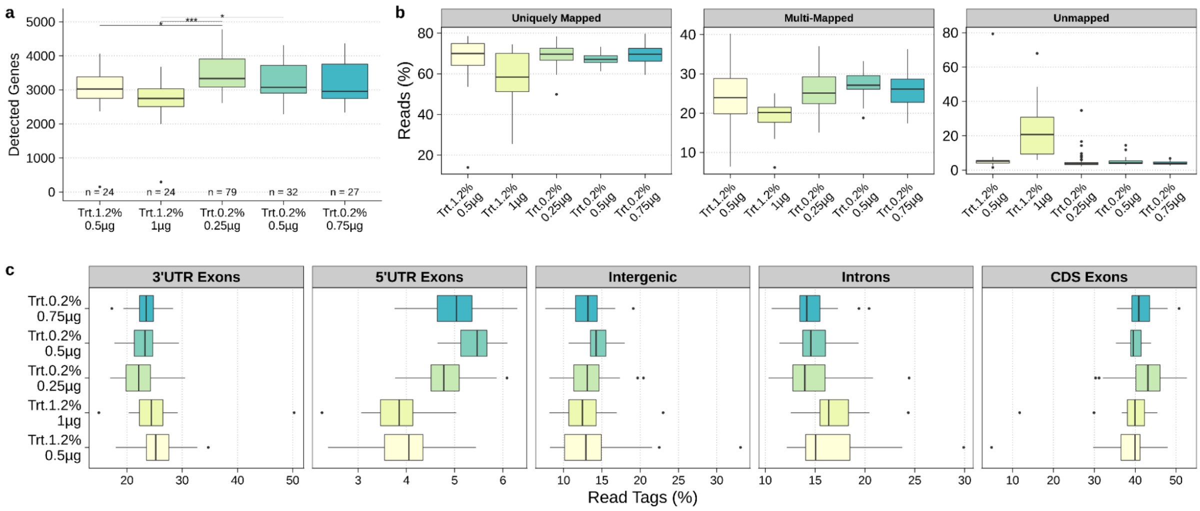


**Fig. E23 | hPBMCs - 5µL - T4g32p Titration and Lysis Buffer** **a.** Number of detected genes (Dunn's test, Bonferroni correction, adj. pval). **c.** STAR mapping statistics showing the percentage of uniquely mapped, multi-mapped and unmapped reads. **d.** Distribution of mapped reads between introns, intergenic regions or 3'-UTR / 5'-UTR / Coding sequence (=CDS) exons. Expressed in percentage of read tags and computed using ReSQC.

**T4g32p 0.25 µg + TSO (1x, 2x, 3x, 4x) [hPBMCs-5µL], Fig. E24:** As in the no-T4g32p condition, we observed a linear relationship between the increase of TSO and the number of detected genes. This effect appeared to be less pronounced than in the no-T4g32p condition (i.e., adj. pval > 0.05) but was still associated with an increased number of multi-mapped and intergenic reads. These observations suggest that increasing the amount of TSO in the reaction is not beneficial.

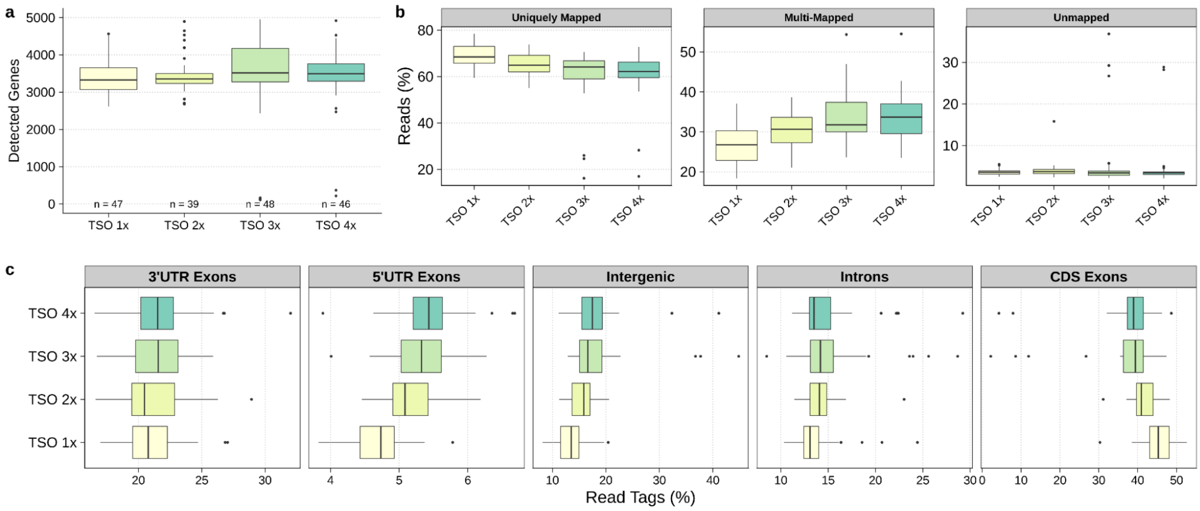

**Fig. E24 | hPBMCs - 5µL - T4g32p and TSO Titration** **a.** Number of detected genes (Dunn's test, Bonferroni correction, adj. pval). **b.** STAR mapping statistics showing the percentage of uniquely mapped, multi-mapped and unmapped reads. **c.** Distribution of mapped reads between introns, intergenic regions or 3'-UTR / 5'-UTR / coding sequence (=CDS) exons. Expressed in percentage of read tags and computed using ReSQC.

**Homemade T4g32p [hPBMCs-25µL], Fig. E25:** We had observed a higher percentage of unmapped reads when using T4g32p. To rule out contaminations by bacterial DNA/RNA related to the T4g32p production, we used a homemade T4g32p which had been thoroughly cleaned by dialysis and for which we had more control on the potential left-over contaminants. To minimize any plate-to-plate variations, in the same 96-well plate we processed 48 cells to which we added 5 µg of T4g32p and 48 cells according to the standard FS protocol. Both RT reactions were carried out at 37°C. Similarly to the commercial enzyme, we observed a higher percentage of multi-mapped, unmapped and intergenic reads when using T4g32p.

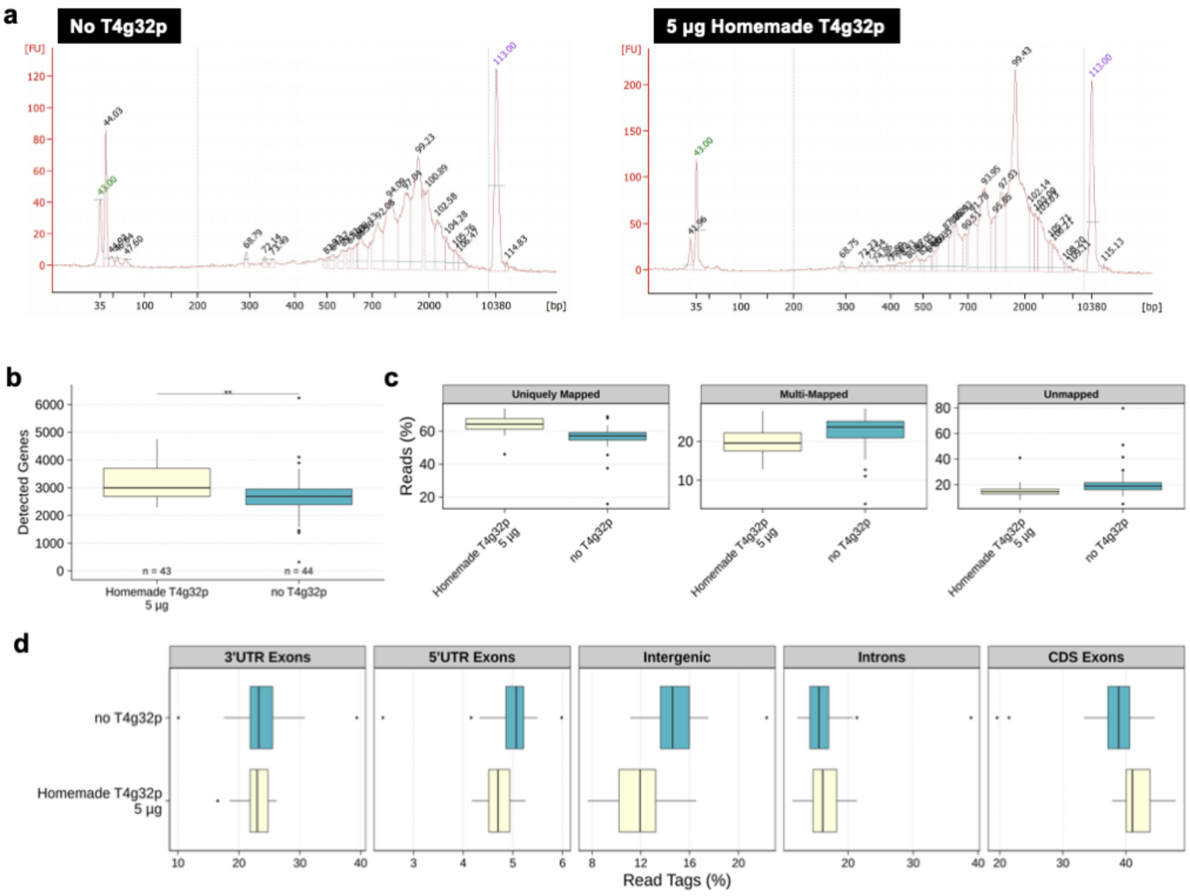

**Fig. E25 | hPBMCs - 25µL - Homemade T4g32p a.** cDNA size distribution **b.** Number of detected genes (Mann-Whitney U test). **c.** STAR mapping statistics showing the percentage of uniquely mapped, multi-mapped and unmapped reads. **d.** Distribution of mapped reads between introns, intergenic regions or 3'-UTR / 5'-UTR / coding sequence (=CDS) exons. Expressed in percentage of read tags and computed using ReSQC.

### Conclusion:

The initial development of FLASH-Seq is the result of years of observations accumulated after developing Smart-seq2[10]. Building on this knowledge, we explored dozens of conditions before settling on the final protocol described in the main text of this study. As it is perhaps to be expected, the vast majority of these changes did not bring any significant benefit in terms of data quality and/or overall cost per-cell. Our results seem to indicate that the improvement of single-cell protocols through additives and optimized reaction conditions may be reaching its limit. A new generation of enzymes with higher processivity and capable of handling sub-picogram levels of RNA is therefore urgently needed to unravel the very lowly expressed genes.

Interestingly, as exemplified by T4g32p, carrying out the reactions in smaller volumes tended to overshadow the effect of several additives that had an impact in larger volumes. It appears that the benefits provided by miniaturization often compensate for those provided by most additives. In addition, we also observed significant differences in working with large (i.e., HEK 293T) vs small cells (i.e., hPBMCs), highlighting the need to perform the tests directly in the conditions which better reflect the "real" experiment a user has in mind.

In conclusion, several additives/reaction conditions such as variable TSO concentrations or different lysis buffer (i.e. BSA or GuHCl) may provide circumstantial benefits and will likely require more in-depth testing to find an appropriate place for them in the FS protocol. We hope that these results can provide valuable information to researchers and help build the next generation of single-cell sequencing methods.

#### Bibliography Extended File #1
